# Postzygotic barriers persist despite ongoing introgression in hybridizing *Mimulus* species

**DOI:** 10.1101/2023.08.05.552095

**Authors:** Samuel J. Mantel, Andrea L. Sweigart

**Author notes:** Corresponding author: Samuel Mantel, 120 E. Green St. Athens, GA 30602.

## Abstract

- The evolution of postzygotic isolation is thought to be a key step in maintaining species boundaries upon secondary contact, yet the dynamics and persistence of hybrid incompatibilities in sympatric species are not well understood.
- Here, we explore these issues using genetic mapping in three populations of recombinant inbred lines between naturally hybridizing monkeyflowers *Mimulus guttatus* and *M. nasutus* from the sympatric Catherine Creek population.
- The three *M. guttatus* founders differ dramatically in admixture history. Comparative genetic mapping also reveals three putative inversions segregating among the *M. guttatus* founders, two due to admixture. We observe strong, genome-wide transmission ratio distortion, but patterns are highly variable among populations. Some distortion is explained by epistatic selection favoring parental genotypes, but tests of inter-chromosomal linkage disequilibrium also reveal multiple candidate Dobzhansky-Muller incompatibilities. We also map several genetic loci for hybrid fertility, including two interacting pairs coinciding with peaks of distortion.
- Remarkably, in this limited sample of *M. guttatus*, we discover abundant segregating variation for hybrid incompatibilities with *M. nasutus,* suggesting this population harbors diverse contributors to postzygotic isolation. Moreover, even with substantial admixture, hybrid incompatibilities between *Mimulus* species persist, suggesting postzygotic isolation might be a potent force in maintaining species barriers in this system.

## Introduction

Despite often being conceptualized as a single event, it has long been understood that speciation is a complex and dynamic process (Darwin, 1859; Dobzhansky, 1937). As new species form (generally in allopatry), the cumulative impacts of adaptive divergence, genetic drift, and the sorting of ancestral variation can lead to the evolution of reproductive isolation, which limits gene flow between species when they come into secondary contact (Dobzhansky, 1937; Mayr, 1942; Coyne & Orr, 2004). Nevertheless, gene flow between recognized species – at rates high enough to have substantial genomic consequences – is not uncommon, even when strong reproductive isolating barriers exist (Nolte *et al*., 2009; Teeter *et al*., 2010; Ellegren *et al*., 2012; Renaut *et al*., 2013; Larson *et al*., 2014; Delmore *et al*., 2015; Caeiro-Dias *et al*., 2023). At the extremes, hybridization can break down established barriers causing previously isolated lineages to fuse (Taylor *et al*., 2006) or strengthen reproductive isolation through reinforcement (Servedio & Kirkpatrick, 1997; Jaenike et al., 2006; Hopkins & Rausher, 2012), but theory also suggests partial reproductive isolation can sometimes evolve as a stable optimum (Servedio & Hermisson, 2020). Studies of naturally hybridizing species can provide a window into the causes of these alternative evolutionary outcomes, allowing us to identify and measure the impact of the genetic loci involved in reproductive isolation.

One powerful approach to address these questions is experimental hybridization of incompletely isolated species that occur together in secondary sympatry. In particular, the use of late-generation hybrids such as recombinant inbred lines (RILs) can enable replication of wild, admixed genotypes and facilitate genetic mapping of barrier loci under controlled, environmental conditions. This approach should be especially fruitful for identifying loci involved in postzygotic isolation because strongly incompatible allele combinations are likely to be purged during RIL formation. Indeed, selection against hybrid incompatibilities can be a major source of transmission ratio distortion (TRD) in RIL populations (Colomé-Tatché & Johannes, 2016). Although there are often additional contributors to TRD in experimental hybrid populations (reviewed in Fishman & McIntosh, 2019), hybrid incompatibilities are expected to leave a particular signature of distortion, generating both local peaks and linkage disequilibrium between unlinked loci caused by underrepresented or absent multi-locus genotypes. This signature will likely be particularly apparent in RILs, which have the advantage of multiple generations of recombination, increasing power to detect and localize incompatibilities by amplifying their effects (and revealing recessive phenotypes) through inbreeding (Moyle & Nakazato, 2008).

Despite the clear value of experimental hybridization, and RILs in particular, for illuminating the genetic causes of reproductive isolation, inferences to natural populations can be challenging because such studies are often based on crosses between only two individuals or inbred lines. This limitation might be serious for investigations of postzygotic isolation between species in the early stages of divergence, as hybrid incompatibilities are often polymorphic (Sweigart *et al*., 2007; Good *et al*., 2008; Cutter, 2012), particularly in populations with ongoing hybridization (Zuellig & Sweigart, 2018a). Variation in chromosomal structure is also a common feature of natural populations (Hoffmann & Rieseberg, 2008), and polymorphic inversions are often associated with adaptive differences (Feder *et al*., 2003; Lowry & Willis, 2010; Jones *et al*., 2012) and reproductive isolation (Noor *et al*., 2001b; Sotola *et al*., 2023). One way to capture a more representative set of natural variation is to generate a multi-parent mapping population, for instance by performing multiple biparental crosses to a common founder (Yu *et al*., 2008). Although this strategy is now routinely used in plant breeding programs (Scott *et al*., 2020) and to dissect the genetics of complex traits (Aylor *et al*., 2011; Long *et al*., 2014) it has rarely been applied to studies of speciation.

In this study we investigate patterns and mechanisms of reproductive isolation in three independent RIL populations constructed from crosses between *Mimulus guttatus* and *M. nasutus* (also referred to as *Erythranthe guttata* and *E. nasuta*: see Lowry *et al*., 2019 and Nesom *et al*., 2019). These two species are closely related (∼200 KYA diverged: Brandvain *et al*., 2014), and broadly allopatric across western North America, but occur in secondary sympatry at many sites in their shared range. One such sympatric site occurs at Catherine Creek (CAC) in Washington state, a system of ephemeral seeps and streams that flow into the Columbia River. These *Mimulus* species have diverged in mating system (*M. guttatus* is large flowered and predominantly outcrossing, whereas *M. nasutus* is a small flowered and primarily selfing), which acts as a strong premating barrier where the two species co-occur (Martin & Willis, 2007). At Catherine Creek, divergent flowering phenology (Fishman *et al*., 2014a; Kenney & Sweigart, 2016) and microhabitat adaptation (Mantel & Sweigart, 2019) are also important contributors to premating isolation. Despite these substantial barriers to interspecific mating, studies of population genomic variation at Catherine Creek have revealed clear signatures of historical and contemporary hybridization (Kenney & Sweigart, 2016), resulting in largely unidirectional introgression from *M. nasutus* into *M. guttatus* (Sweigart & Willis, 2003; Brandvain *et al*., 2014; Kenney & Sweigart, 2016). Even in the absence of interspecific gene flow, *M. guttatus* populations are exceptionally genetically diverse (Flagel *et al*., 2014; Puzey *et al*., 2017) and maintain substantial variation for complex phenotypes (Troth et al., 2018).

Here, we attempt to capture and leverage this diversity at Catherine Creek to understand the patterns and mechanisms of hybrid breakdown in naturally hybridizing species. We generate three independent RIL populations by crossing distinct CAC *M. guttatus* progenitors to a common *M. nasutus* parent. Using this powerful crossing design, we discover that differences in admixture have cascading effects on structural variation and reproductive barriers. Moreover, even in this small sample of three *M. guttatus* lines, we find segregating variation for hybrid incompatibilities with *M. nasutus*, suggesting this population harbors diverse contributors to postzygotic isolation. Finally, we discover that even with substantial admixture, hybrid incompatibilities between *Mimulus* species persist, suggesting postzygotic isolation might be a potent force in maintaining species barriers in the face of ongoing hybridization.

## Materials and Methods

### Plant Material and Crossing Design

The plants used in this study are derived from a collection of inbred lines (>10 generations inbred, single seed decent) produced from wild-collected seeds from Catherine Creek (CAC). We produced three parallel RIL populations from crosses between these inbred lines of CAC *M. guttatus* and *M. nasutus*. We selected three previously studied CAC *M. guttatus* inbred lines (CAC110, CAC162, and CAC415; TableS1; Mantel & Sweigart, 2019) with stereotypic *M. guttatus* morphological and phenological characteristics and which produced fertile F1s with *M. nasutus* (∼72-99% viable pollen). One *M. nasutus* inbred line (CAC9) served as a common maternal parent, which we emasculated before pollen development, and crossed by hand with pollen from each of the *M guttatus* inbred lines, producing three F1 hybrids. For each of these F1 hybrids, we self-fertilized a single plant to generate three corresponding F2 populations (*N* = 500 plants each). We performed several additional rounds of self-fertilization using single-seed-descent to produce three populations of F6 or F7 recombinant inbred lines (RILs; Figure S1), generating a total of 368 RILs (151 CAC110 RILs, 128 CAC162 RILs, and 89 CAC415 RILs), much lower than the initial 1500 due to line extinction during the inbreeding process.

### Library Preparation, Sequencing, and Processing

We grew one individual of each of the 368 RILs (F6 generation or later) for tissue collection and DNA extraction in the greenhouse at the University of Georgia, conditions described in Supporting Information Methods S1. We collected bud and meristem tissue from each individual into 96-well plates on dry ice, stored them at −80°C, and extracted genomic DNA using a modified CTAB method in a 96-well format (dx.doi.org/10.17504/protocols.io.bgv6jw9e). We then constructed ddRADseq libraries using a modified version of RAD Capture (Ali *et al*., 2016), “BestRAD” (dx.doi.org/10.17504/protocols.io.6awhafe, Supporting Information Methods S1). We prepared libraries for, and sequenced the same RILs twice, each on two partial lanes of Illumina HiSeq4000 for paired end 150-bp reads at the Genome Sequencing Facility at Duke University. We also extracted genomic DNA from bud and leaf tissue of the three parental CAC *M. guttatus* lines using a modified CTAB method as above. Whole genome sequence (WGS) libraries were prepared and sequenced on a partial lane of Illumina HiSeq4000 for paired end 150-bp reads at the Genome Sequencing Facility at Duke University.

We aligned WGS reads to the *Mimulus guttatus* IM62 v3 refence assembly (phytozome.org), IM: Iron Mountain, an allopatric *M. guttatus* population in Oregon, ∼200km south of CAC), called SNPs, removed heterozygous calls and filtered sites to a minimum depth of 3x, and a maximum depth of 2 standard deviations above the mean. To aid in genotyping *M. guttatus* and *M. nasutus* SNPs across the genome, we identified a full list of diagnostic SNPs (i.e., those homozygous for different alleles in each of the parental pairs, CAC9 & CAC110, CAC9 & CAC162, and CAC9 & CAC415). We joint genotyped each pair of parental lines, quality filtered the called SNPs, and retained only biallelic SNPs, extracting a list of high-quality, diagnostic SNPs for each pair of parental lines. Details of sequence data processing are described in Supporting Information Methods S1.

Following sequencing, we demultiplexed the ddRADseq data and aligned reads to the *Mimulus guttatus* IM62 v3 reference assembly. At this stage, six samples were removed from further analysis (4 from the CAC162 RILs, and 2 from the CAC415 RILs) due to low similarity between duplicate BAMs produced from the two independent ddRAD library preparations. We then merged the remaining duplicate BAMs for a total of 362 RIL individuals (151 CAC110 RILs, 124 CAC162 RILs, and 87 CAC415 RILs). We called SNPs, joint genotyped each RIL population with its corresponding parental lines, and quality filtered called SNPs. Finally, we removed non-diagnostic sites between each pair of parental lines.

To assess levels of residual heterozygosity in the RIL populations, we filtered the three RIL VCFs as above and requiring a minimum depth of 10x. Heterozygosity was low and uniform across the genome: CAC110 RILs: 1.95%, N=13,975; CAC162 RILs: 2.34%, N=33,340; CAC415 RILs: 2.79%, N=23,178 and consistent with the Mendelian expectation (∼1.5-3%, in F6/F7 RILs).

### Assessment of Introgression in Parental Lines

As previous studies have shown that admixture between *M. guttatus* and *M. nasutus* is common at CAC (Brandvain *et al*., 2014; Kenney & Sweigart, 2016) we first checked the selected CAC *M. guttatus* parental lines for evidence of introgression from *M. nasutus*. To coarsely identify putative regions of introgression in *M. guttatus* parental lines, we calculated *Dxy* between CAC9 and the three parental *M. guttatus* lines in non-overlapping 50-kb windows containing 5000 or more called sites. To more concretely assess evidence of *M. nasutus* introgression in the parental CAC *M. guttatus* lines, we used the newly generated WGS data, plus WGS data from 7 other inbred *Mimulus* accessions downloaded from the SRA (https://www.ncbi.nlm.nih.gov/sra, Table S1): the parental *M. nasutus* line CAC9, 3 non-CAC *M. nasutus* lines, the 3 parental *M. guttatus* lines from CAC, 5 additional non-parental *M. guttatus* lines from CAC, and 2 non-CAC *M. guttatus* lines). We processed reads from additional (non-parental) lines as above and joint genotyped all lines together. We filtered called SNPs based on quality as above, and retained only biallelic SNPs.

We identified putative regions of introgression using a Hidden Markov Model (HMM, modified from Durbin *et al*., 1998 and developed in Brandvain *et al*., 2014. As an input for the model, we identified fourfold degenerate sites and non-degenerate sites in the IM62 v3 reference assembly (https://github.com/tsackton/linked-selection/tree/master/misc_scripts). With these lists and the VCF of high-quality filtered SNPs in parental and non-parental *Mimulus* lines described above, we counted the total number of genotyped sites, genotyped non-degenerate sites, and genotyped fourfold degenerate sites in 1-kb windows in each line. We then counted the number of those sites that differed in their SNP call between each pairwise comparison of lines. With these counts and the HMM, we calculated posterior probabilities of *M. nasutus* and *M. guttatus* ancestry across the genomes of the parental CAC *M. guttatus* lines. We considered 1-kb windows with a calculated posterior probability of *M. nasutus* ancestry >0.95 to be windows showing evidence of introgression.

### RIL Genotyping and Linkage Mapping

We genotyped the RILs in non-overlapping 50-kb windows; windows with >75% *M. guttatus* calls were coded as *M. guttatus* homozygotes, 25-75% as heterozygotes, and <25% as M. nasutus homozygotes. Windows with no SNP calls were coded as missing data. To fill these gaps, we used Python scripts from the processing steps of the program GOOGA (Flagel *et al*., 2019) to calculate genotype probabilities for each window in each RIL individual. In GOOGA, we set the expected genotype frequencies to 1:1 for alternative parental homozygotes (heterozygotes are excluded given that they are <3% of the genotypes in this advanced RIL population). We then used the program to estimate genotype error rates and recombination rates for each windowed genotype for each RIL. We dropped 39 RILs from the dataset due to excessive missing data (stemming from low read coverage) because they produced high (e1 or e2 > 0.2) estimated error rates in GOOGA (8 from the CAC110 RILs, 26 from the CAC162 RILs, and 5 from the CAC415 RILs). Finally, we used GOOGA’s HMM to estimate genotype probabilities for each 50-kb window of the remaining RILs and converted to hard calls excluding windows with probabilities <0.75. Of the original 368 RILs, this resulted in genotype data for 323 of them (143 of the original 151 from the CAC110 RILs, 98/128 from the CAC162 RILs, and 82/89 from the CAC415 RILs). We used LepMAP3 (Rastas, 2017) to construct a linkage map for each of the three RIL populations. Linkage mapping details are described in Supporting Information Methods S1 and Table S2. We produced three linkage maps containing 3,828, 4,389 and 4,379 markers, retaining 794, 606, and 690 that mapped to recombination-informative sites (for the CAC110, CAC162, and CAC415 RIL populations, respectively).

### Assessment of Transmission Ratio Distortion and Linkage Disequilibrium

Following linkage mapping, we calculated genotype frequencies at each marker in each map and identified regions of transmission ratio distortion (TRD) by testing for deviations from the Mendelian 1:1 expectation with χ^2^ tests. We used both an uncorrected α = 0.01 and a more stringent Bonferonni-corrected ⍺ (unique ⍺ for each chromosome in each RIL population depending on the number of markers tested; Table S3). To investigate whether peaks of TRD were associated with major two-locus hybrid incompatibilities we used the package pegas (Paradis, 2010) to characterize pairwise linkage disequilibrium (LD) between all markers with unique map positions. To minimize false positives driven by the large number of pairwise comparisons, we identified putative interacting loci associated with inter-chromosomal LD based on r values greater than the 95% percentile (CAC110 RILs: r > 0.37, CAC162 RILs: r > 0.41, CAC415 RILs: r > 0.39). At these interacting loci we then performed likelihood ratio tests comparing observed genotype ratios to expected ratios (calculated by cross tabulation of observed allele frequencies), and describe epistatic interactions where *p*<0.0001.

### Assessment of Hybrid Male and Female Sterility

To measure male fertility, we collected pollen from three flowers (second flower pair or later) of each RIL individual. We suspended the pollen of each flower separately in lactophenol-aniline blue stain (Kearns & Inouye, 1993) and inspected the pollen grains using a compound microscope. For each flower, we estimated pollen fertility by calculating the proportion of darkly stained (viable) pollen grains from a haphazardly selected group of >100 within a field of view in of the microscope. When fertility estimates from flowers of the same plant differed by more than 0.2, pollen from up to three more flowers was collected and counted. We then estimated individual pollen viability as the average value from all collected flowers. Because the distribution of pollen viability in each RIL population was left-skewed, we logit-transformed the data to more closely match a normal distribution.

To estimate hybrid female sterility, we intercrossed random pairs of RILs in at least duplicate, and collected all seeds produced. We counted the number of seeds produced per fruit for each cross made and ran a linear mixed model (LMM, using the lmer command in the ‘lme4’ package). In addition to maternal parent genotype (fixed effect), the model included average viable pollen proportion of the paternal parent (as calculated above, fixed effect), a unique RIX identifier nested within RIX cross type (CAC110 RIL x CAC162 RIL, CAC162 RIL x CAC415 RIL, etc.; random effect) and the year that the cross was made (crosses were made during two separate grow-outs in separate years; random effect). We then calculated least square means of female fertility (using seeds per fruit as a proxy) for each RIL genotype using the ‘emmeans’ package. This method produced distributions with a long right tail, so we square root transformed the data to more closely match a normal distribution.

### QTL Mapping

Given the small sample sizes of each RIL population, we took a multistep approach to genetically map hybrid male and female sterility. For each trait in each RIL population, we first used r/QTL (Broman & Sen, 2009; Arends *et al*., 2010) to perform 10 iterations of composite interval mapping (CIM; Zeng, 1993, 1994; Jansen & Stam, 1994). Because CIM requires no missing data, missing genotypes are imputed before the scan is run, leading to potential differences among iterations. We set lenient LOD thresholds (α = 0.15) for QTL significance, by performing 5,000 permutations, and calculated 1.5-LOD intervals (‘lodint’ command) for each significant peak position in any of the 10 iterations. We then performed single-QTL scans using each QTL identified with CIM as an interactive covariate (‘intcovar’ option) in separate runs. We again we set LOD thresholds (α = 0.15, 10,000 permutations) for interacting QTL significance, and calculated 1.5-LOD intervals for each significant peak position. With these identified peaks, we then fit a multiple-QTL model (‘fitqtl’ command; HK method) and directly estimated additive QTL effects (at each peak position) and identified epistatic effects. We then removed any nonsignificant (p > 0.05) effects, and re-fit the model.

## Results

### Admixture in M. guttatus parental lines

All *M. guttatus* lines tested show clear evidence of admixture from *M. nasutus*, but the genomic location and proportion of introgression vary considerably across lines (Figure 1, Figure S2, Figure S3). Of the three parents, CAC110 is the most admixed with ∼24% of its genome showing very low genetic divergence (*Dxy*) and a high posterior probability of recent coalescence with *M. nasutus* via probabilistic inference (HMM, see Methods; Brandvain *et al*., 2014). At least some of this introgression in CAC110 appears to be recent, with large portions of entire chromosome arms showing evidence of introgression (Figure S2). Although the largest inferred contiguous blocks in CAC110 are only ∼1.1Mb (Figure S4b), the HMM may sometimes break up true contiguous blocks of introgression (or incorrectly identify short regions with low *Dxy* as admixture blocks – see Brandvain et al. 2014). Given the improbability of inheriting a large number of small introgressions immediately adjacent to one another, we think it likely that several regions in CAC110 (on Chromosomes 1, 8, 11, and 13 – Figure 1 and Figure S2) represent true contiguous blocks of admixture. In both CAC162 and CAC415, the overall proportion of admixture is lower (∼9% in each) and blocks are generally smaller and more evenly dispersed throughout the genome (Figure S2; Figure S4b), consistent with introgression events that occurred in the more distant past. Even in CAC110, however, there are many very small regions of inferred introgression (Figure S4b), suggesting *M. guttatus* genomes at Catherine Creek are shaped by a long history of introgression from *M. nasutus*.

**Figure 1.**
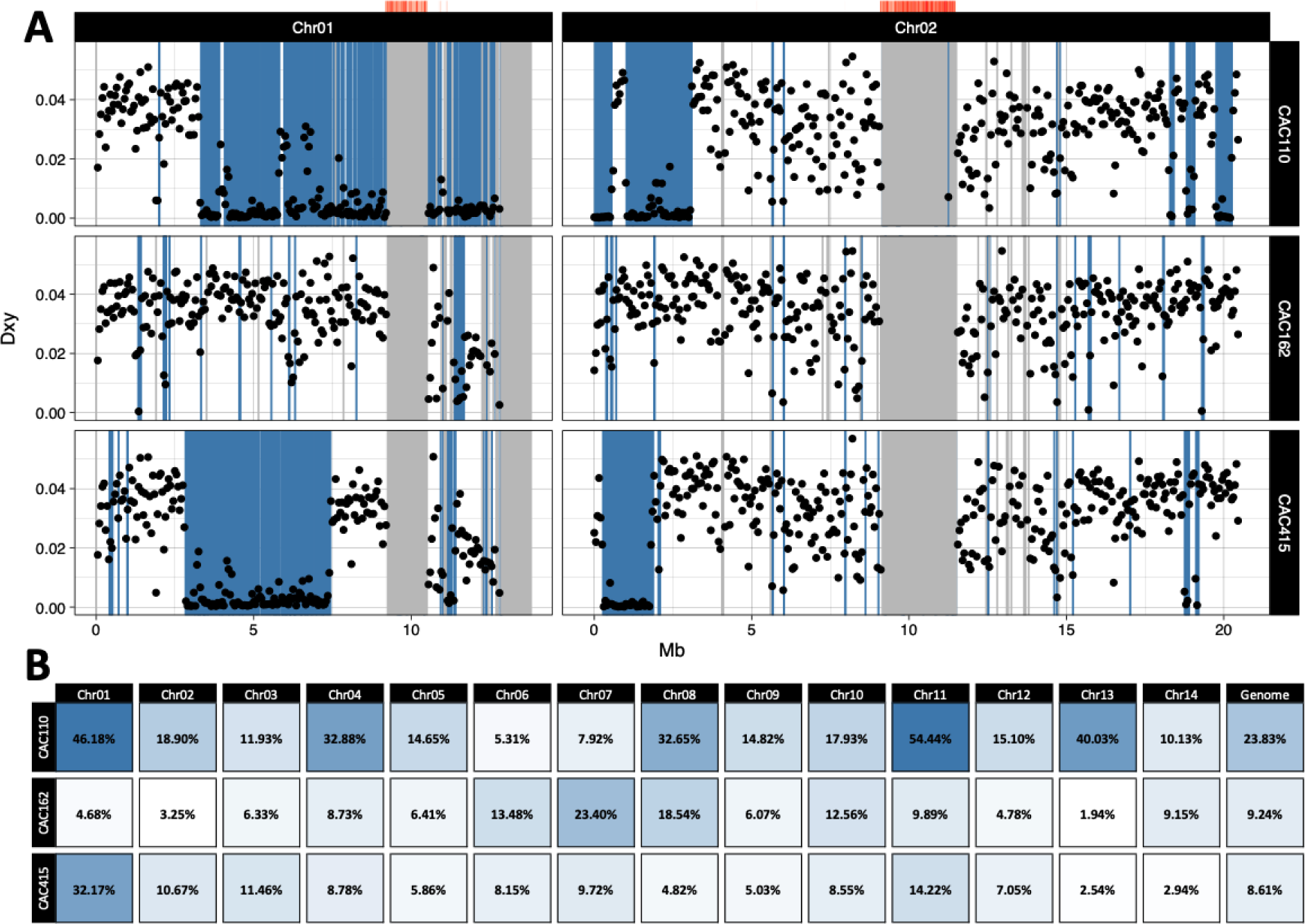
Introgression from *M. nasutus* into sympatric *M. guttatus.* **(a)** *Dxy* calculated in 50kb windows across the first two chromosomes of the IM62 v3 *Mimulus guttatus* reference genome, between the parental lines of the three RIL populations (CAC9 v. CAC110, CAC9 v. CAC162, and CAC9 v. CAC415). Blue shaded regions indicate putatively admixed regions in the three focal CAC *M. guttatus* lines tested, diagnosed by a greater than a 95% posterior probability of *M. nasutus* ancestry (in 1kb windows) as inferred via the HMM (i.e. more recent coalescence with included *M. nasutus* samples than with included *M. guttatus* samples). Grey shaded regions indicate missing data (fewer than 5000 genotyped sites within any 50kb window). Red heatmaps above each chromosome indicate the density of centromeric repeats in the same 50kb windows. All chromosomes are shown in Figure S2. **(b)** Heatmap showing the proportion of *M. nasutus* ancestry in each chromosome, and across the genome, of each of the three CAC *M. guttatus* lines used as parents of the RIL populations, as estimated as the proportion of 1kb windows with a greater than a 95% posterior probability of *M. nasutus* ancestry as inferred via the HMM. Darker blue shading indicates a higher proportion of *M. nasutus* ancestry.

### Comparative linkage mapping

Across most of the genome, the three RIL linkage maps are colinear with similar lengths (Table 1), and marker order generally follows the physical order of the *M. guttatus* IM62 v3 reference assembly (Figure 2; Figure S5). However, we identified three clear exceptions to this pattern, where maps differ due to parental variation in putative inversions, characterized by a large number of physically dispersed markers mapping to the same genetic position. Two of these putative inversions are on LG1 (*inv1.1* and *inv1.2;* Figure S6) and are potentially unique to Catherine Creek (or locally distributed), having not been reported in other crosses with these *Mimulus* species (Fishman *et al*., 2014b; Flagel *et al*., 2019). In both cases, the CAC9 *M. nasutus* parent carries the ancestral orientation and the *M. guttatus* parents segregate for the derived variant. At *inv1.1*, recombination is suppressed in the CAC110 and CAC415 RILs, but not in the CAC162 RIL, indicating collinearity between the CAC9 *M. nasutus* and CAC162 *M. guttatus* parents (Figure 2; Figure S6). At *inv1.2*, on the other hand, the genetic maps suggest CAC9 is collinear only with CAC110 (Figure 2; Figure S6). Remarkably, this *M. guttatus* parent also carries a large block of introgression spanning this entire region (Figure 1), suggesting that gene flow from *M. nasutus* has re-introduced the ancestral variant into CAC110. We discovered a similar situation on LG13, where introgression from *M. nasutus* accounts for collinearity between CAC9 and CAC110 at *inv13* (recombination rates in the CAC110 RIL are much higher in this region than in the CAC162 and CAC415 RILs; Figure S5). Additionally, *inv13* might have arisen in *M. nasutus*, as it also appears to be present in the reference genome assembly of *M. nasutus* SF5 (an accession derived from Shear’s Falls, an allopatric *M. nasutus* population ∼70 km from CAC), but not in other recently assembled *M. guttatus* genomes (phytozome.org). Taken together, these mapping results point to introgression from *M. nasutus* as a major source of large-scale structural variation at Catherine Creek, responsible both for restoring ancestral variants and for introducing new derived inversions into *M. guttatus*.

**Figure 2.**
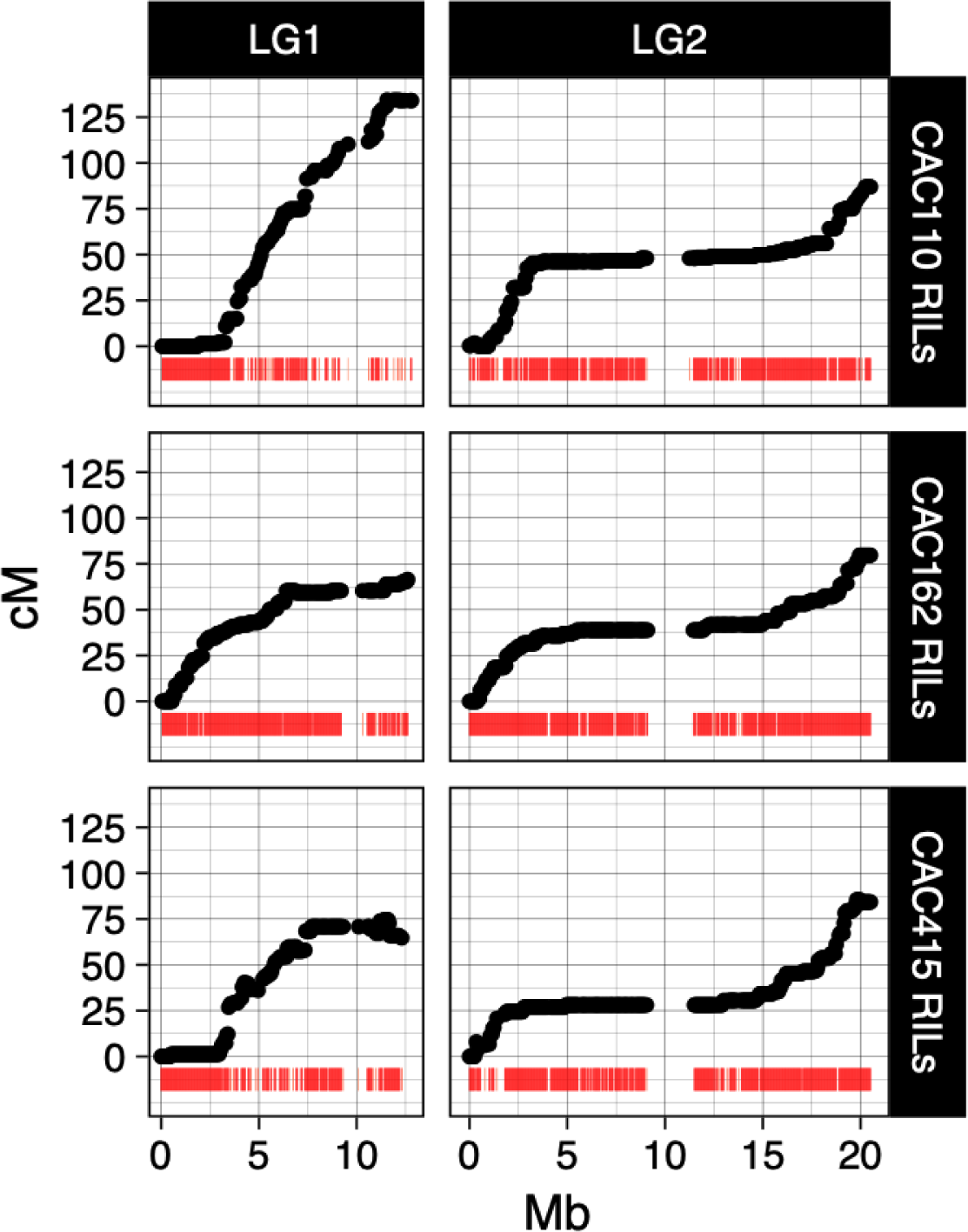
Map position (cM) versus physical marker positions (Mb) for the first two chromosomes of the IM62 v3 *Mimulus guttatus* reference genome in the three RIL populations. Red heatmaps below each curve indicate the density of markers, measured as the distance in Mb to the nearest neighboring marker. Low SNP density, and the subsequent inability to genotype markers in areas of the *M. guttatus* parental genomes showing evidence of introgression from *M. nasutus,* manifests as regions of low marker density on some chromosomes. All linkage groups are shown in Figure S5.

**Table 1.**
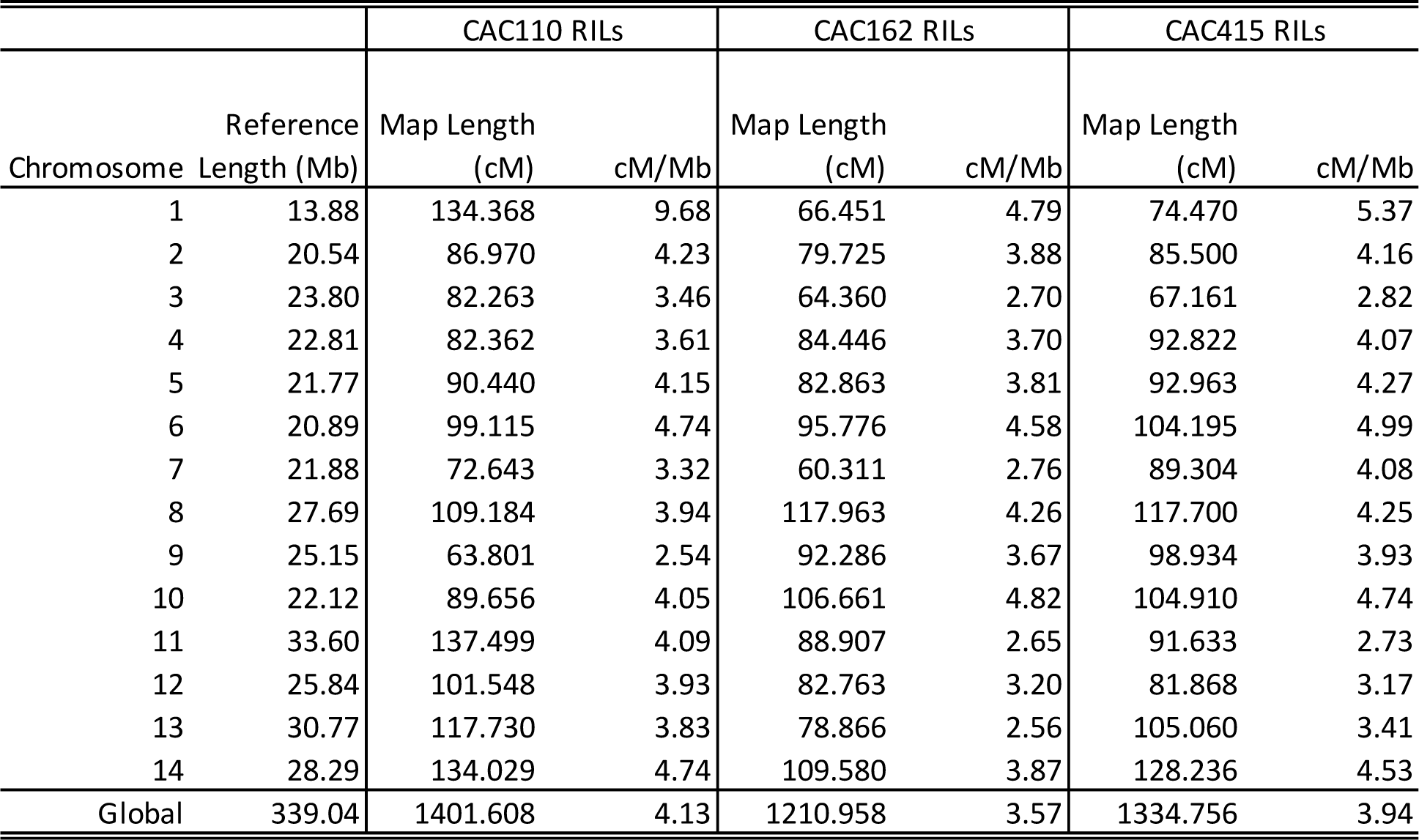
Summary of linkage map length and intrascaffold recombination rate estimates. Total map length and intrascaffold recombination rate estimates for each chromosome, plus total map length and average recombination rate in each of the three RIL populations.

|  |  | CAC110 RILs |  | CAC162 RILs |  | CAC415 RILs |  |
| --- | --- | --- | --- | --- | --- | --- | --- |
| Chromosome | Reference Length (Mb) | Map Length (cM) | cM/Mb | Map Length (cM) | cM/Mb | Map Length (cM) | cM/Mb |
| 1 | 13.88 | 134.368 | 9.68 | 66.451 | 4.79 | 74.470 | 5.37 |
| 2 | 20.54 | 86.970 | 4.23 | 79.725 | 3.88 | 85.500 | 4.16 |
| 3 | 23.80 | 82.263 | 3.46 | 64.360 | 2.70 | 67.161 | 2.82 |
| 4 | 22.81 | 82.362 | 3.61 | 84.446 | 3.70 | 92.822 | 4.07 |
| 5 | 21.77 | 90.440 | 4.15 | 82.863 | 3.81 | 92.963 | 4.27 |
| 6 | 20.89 | 99.115 | 4.74 | 95.776 | 4.58 | 104.195 | 4.99 |
| 7 | 21.88 | 72.643 | 3.32 | 60.311 | 2.76 | 89.304 | 4.08 |
| 8 | 27.69 | 109.184 | 3.94 | 117.963 | 4.26 | 117.700 | 4.25 |
| 9 | 25.15 | 63.801 | 2.54 | 92.286 | 3.67 | 98.934 | 3.93 |
| 10 | 22.12 | 89.656 | 4.05 | 106.661 | 4.82 | 104.910 | 4.74 |
| 11 | 33.60 | 137.499 | 4.09 | 88.907 | 2.65 | 91.633 | 2.73 |
| 12 | 25.84 | 101.548 | 3.93 | 82.763 | 3.20 | 81.868 | 3.17 |
| 13 | 30.77 | 117.730 | 3.83 | 78.866 | 2.56 | 105.060 | 3.41 |
| 14 | 28.29 | 134.029 | 4.74 | 109.580 | 3.87 | 128.236 | 4.53 |
| Global | 339.04 | 1401.608 | 4.13 | 1210.958 | 3.57 | 1334.756 | 3.94 |

In addition to the effects of structural variation, we observed some instances of map length variation that appear to be associated only with parental differences in introgression. In the CAC110 RILs, introgressed regions on LGs 1, 11, 12, and 13 have elevated genetic distances relative to the same (non-introgressed) regions in the other two RIL populations (Table 1, Figure 2, Figure S5). Rather than reflecting a biological difference in recombination frequency between introgressed and non-introgressed regions, we attribute this effect to technical differences arising from the much lower marker densities in introgressed regions, which are highly similar to the CAC9 parent (Table 1, Figure 2, Figure S5).

### Genome-wide patterns of transmission ratio distortion (TRD)

With the exception of LG9, we observed strong TRD in at least one RIL population on all linkage groups (Figure 3, Figure S7, Table S3). Globally, patterns of TRD are highly variable across RIL populations. In the CAC110 and CAC415 RILs, the majority of distorted markers show an excess of *M. guttatus* genotypes (∼59% in CAC110 and ∼80% in CAC415), whereas in the CAC162 RILs, there is a strong deficit of *M. guttatus* across the genome (∼89% of distorted markers show an excess of *M. nasutus*). In a few cases, the same region is even distorted in opposite directions (e.g., at the beginning of LG4: *M. guttatus* excess in CAC110 RILs, *M. nasutus* excess in CAC162 RILs; Figure 3, Figure S7). Despite this general variability, there are a few instances of shared distortion in multiple RIL populations, potentially implying the same causal mechanisms. Most notably, the proximal end of LG14 shows an excess of *M. guttatus* alleles in all three populations. There is also an excess of *M. guttatus* alleles on LGs 1 and 4 and an excess of *M. nasutus* alleles on LGs 8, 12, and 13 in two of the three RIL populations (CAC110 and CAC162). These regions of shared distortion are largely confined to segments of the genome that are non-introgressed in parental lines. In addition to local peaks of TRD among homozygous marker genotypes, we also investigated levels of residual heterozygosity in the three RIL populations. At the SNP level (RIL windowed genotypes exclude heterozygous calls; see Methods), we observed low levels of heterozygosity (∼1.5-3%) consistent with Mendelian expectations in F6/F7 RILs. Residual heterozygosity was broadly uniform across the genome, with no clear regions enriched for heterozygous SNP calls.

**Figure 3.**
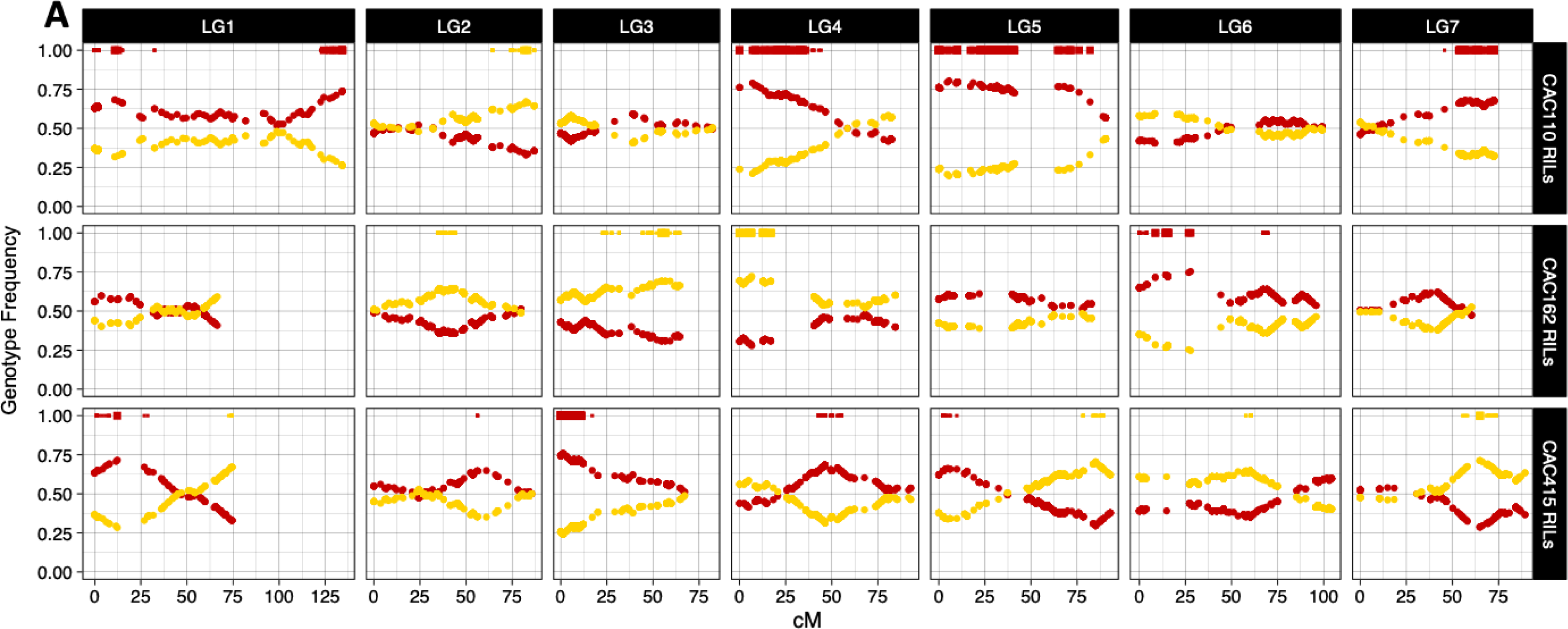
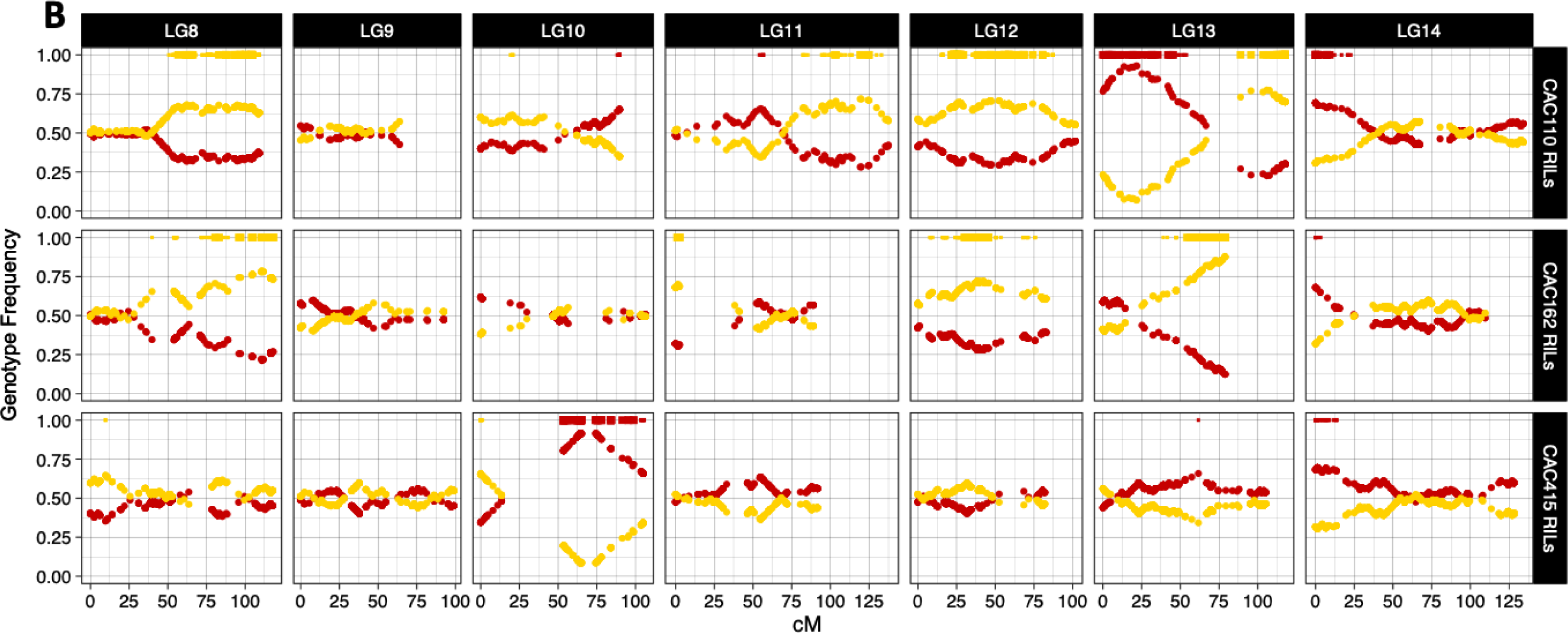
Transmission ratio distortion by map position, (a) Linkage group (LG) 1-7, (b) LG8-LG14. Genotype frequencies in windowed 50kb markers across the 14 linkage groups of the three RIL populations. Red dots indicate homozygous *M. guttatus* genotypes, yellow dots, homozygous *M. nasutus* genotypes. Bars at the top of each plot indicate regions showing significant transmission ratio distortion from the Mendelian 1:1 expectation by χ^2^ tests at ⍺ = 0.01 (thin bars), and a more stringent Bonferonni-corrected ⍺ (thick bars, unique ⍺ for each chromosome in each RIL population depending on the number of markers tested, see Table S3 for values). Bar color indicates an excess of *M. guttatus* (red), or *M. nasutus* (yellow) genotypes. Markers are ordered by genetic position within each map (cM).

### Hybrid incompatibilities as a source of TRD

In each RIL population, we identified several instances of two-locus epistasis (10 in the CAC110 RILs, 4 in the CAC162 RILs, 9 in the CAC415 RILs: Table 2; Table S5). All but one of these 23 cases are due to a deficit of heterospecific allele combinations, consistent with the action of hybrid incompatibilities reducing viability or fertility during RIL formation. In 16 of these cases, the underlying mechanism seems to be epistatic selection favoring the same parental genotypes – that is, we observed either an excess of *M. guttatus* genotypes or *M. nasutus* genotypes at *both* loci. In six cases, however, there are clear deficits of only one of the two heterospecific genotype combinations *and* the expected single-locus distortion stemming from selection against the incompatible genotype at each locus (Table 2). This asymmetry is a hallmark of Dobzhansky-Muller incompatibilities. In four of the six cases, loci involved in these incompatibilities occur exclusively in non-introgressed regions. However, in two of the pairs, one or both loci reside in introgressed blocks, suggesting there may also be segregating incompatibilities within CAC *M. nasutus*. As with overall patterns of TRD, regions involved in two-locus epistasis are largely population specific (although in some cases, we likely have limited power to detect shared epistasis due to small sample sizes and genetic background effects).

**Table 2.**
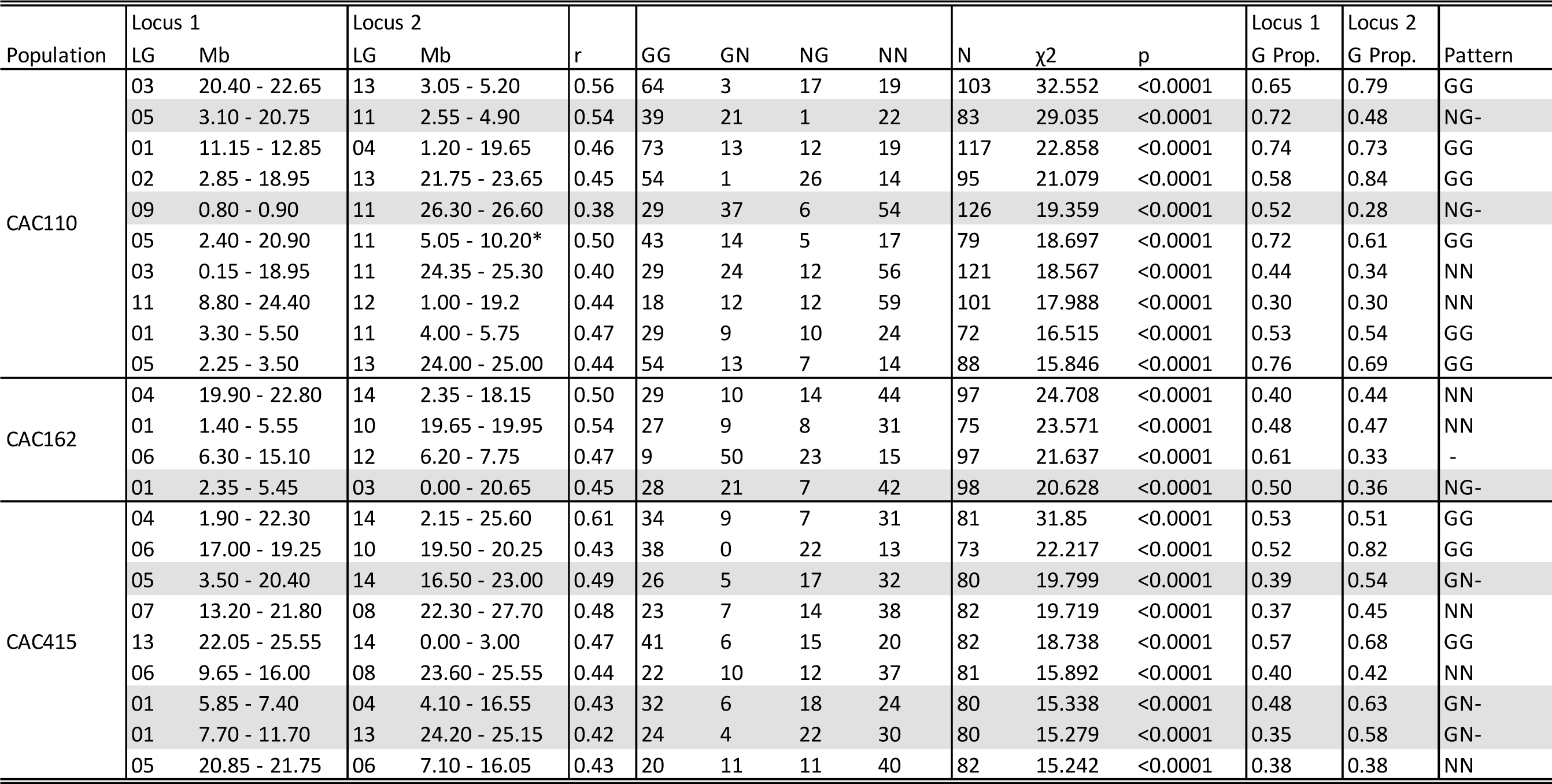
Positions of two-locus epistasis. Physical (Mb) positions of pairs of loci on each linkage group (LG) in the three RIL populations that show LD (r) in the 95^th^ percentile, as well as Chi-Square statistics from likelihood ratio tests with p<0.0001. Counts of each genotypic class in the population are shown. Loci marked with an asterisk have peak r and χ^2^ values at adjacent markers. Single locus proportions of *M. guttatus* alleles (G Prop.) were used to determine if the pattern of TRD at each epistatic locus could be explained by a two-locus hybrid incompatibility (NG- or GN-), or if epistatic selection favoring the same parental genotypes better explains the pattern (GG or NN). The best candidates for two locus hybrid incompatibilities are shaded grey. See Table S5 for full list of epistatic loci in LD 95^th^ percentile.

### Genetics of hybrid fertility traits

Despite six generations of selfing, all three RIL populations exhibited substantial variation in male fertility. Some of this variation may be due to inbreeding depression in *M. guttatus* parental lines, which showed lower pollen viability than the *M. nasutus* parent (Figure S14). However, heterospecific interactions are likely also involved, as all three RIL populations include a considerable fraction of individuals (∼10-22%) with pollen viability <50% (well below the *M. guttatus* average of 72%; Figure S14). In the CAC110 RIL population, we mapped five pollen viability QTLs accounting for ∼36% of variance, including two separate epistatic pairs (Table 3; Figure 4). One of these epistatic pairs leads to reduced male fertility in RILs with *M. nasutus* alleles on LG4 and *M. guttatus* alleles on LG14 (Figure S15a). The pattern of TRD at these same loci (excess of *M. guttatus* on LG4 and *M. nasutus* alleles on LG14; Figure 3) might suggest the action of a gametic incompatibility that reduces pollen viability, but other genetic mechanisms are also possible (e.g., a sporophytic incompatibility that reduces pollen viability to the point of preventing effective self-fertilization/seed production). The second epistatic pair in the CAC110 RIL population – between QTLs on LG12 and LG14 – also arises due to a negative heterospecific interaction (RILs homozygous for *M. nasutus* alleles on LG12 and *M. guttatus* on LG14 have the lowest fertility; Figure S15c), although *M. nasutus* genotypes have higher pollen fertility at each QTL individually (Table 3). At the remaining QTL for male fertility on LG11, *M. nasutus* alleles result in lower pollen viabilities (opposite to parental values), suggesting there could be additional unmapped loci that interact with this QTL to decrease male fertility. In the CAC162 RIL population, we mapped two QTLs for pollen viability (accounting for ∼28% of the variance), but because we detected no epistasis between them and *M. nasutus* alleles increased fertility at both, these effects might be due to inbreeding depression or to incompatibilities with additional loci below our detection thresholds. We mapped no QTLs for male fertility in the CAC415 RIL population.

**Figure 4.**
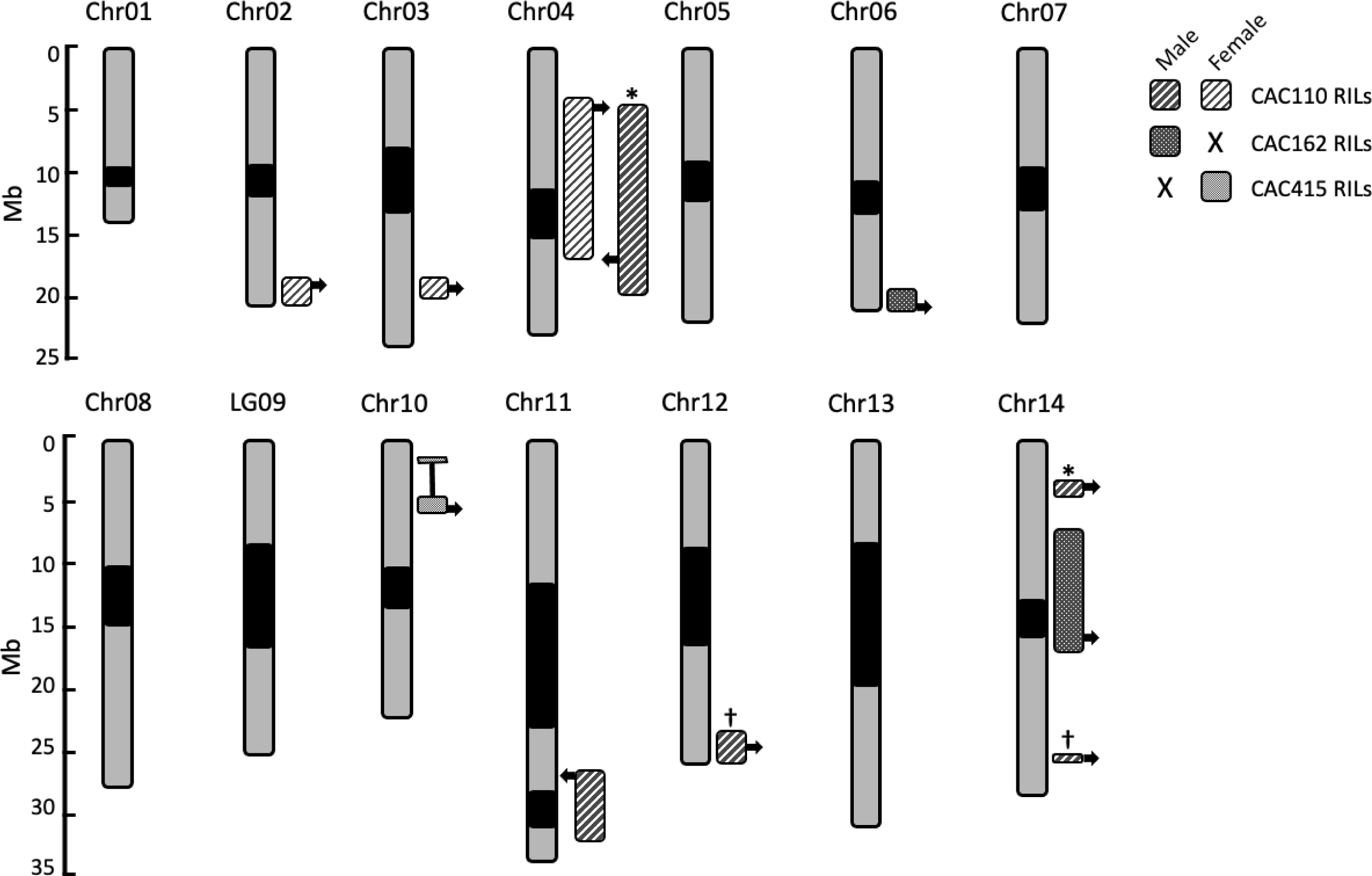
Male and female fertility QTL locations in each mapping population. Physical position (Mb) of the 1.5-LOD interval of each significant QTL is shown. Phenotype and RIL population indicated in key. Arrows indicate the position of peak marker(s). Arrows pointing left indicate the *M. guttatus* allele has the lower value, arrows pointing right indicate the *M. nasutus* allele has the lower value. QTLs with significant epistatic effects are indicated with shared symbols. The 1.5-LOD interval of the QTL underlying female fertility on Chr10 in the CAC415 RILs spans a breakpoint of the known derived inversion in the reference *M. guttatus* line IM62, leading to a disjunct interval. Centromere locations are indicated in black.

**Table 3.**
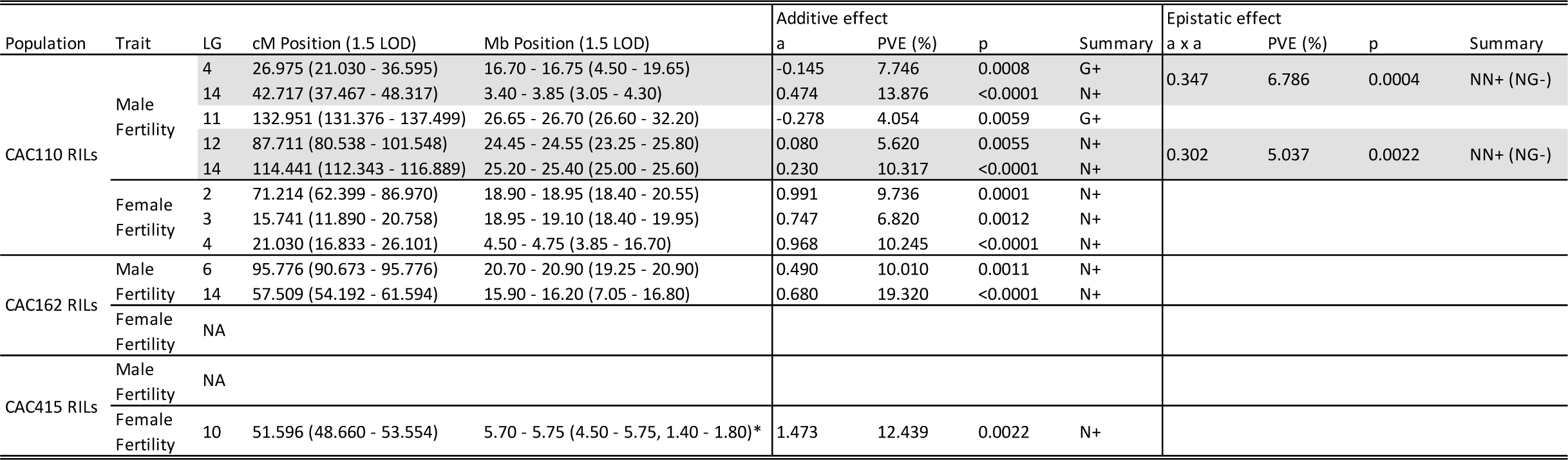
Male and female fertility QTL locations and effects in each mapping population. Linkage group (LG), map (cM), and physical (Mb) position of each significant QTL is shown. Additive effects (a) are negative when the *M. guttatus* allele has the lower value, indicated in the summary column. QTLs with significant epistatic effects are shaded together. Percentage of RIL population variance (PVE) is shown for each additive and epistatic effect. The 1.5-LOD interval of the QTLs underlying female fertility on LG10 in the CAC415 RILs, marked with an asterisk, spans a breakpoint of the known derived inversion in the reference *M. guttatus* line IM62, leading to a disjunct interval. Effect plots are shown in Figures S15-S18.

For female fertility, all three *M. guttatus* parents had much lower fruit production than the *M. nasutus* parent, suggesting substantial inbreeding depression affecting this trait (Figure S14). Accordingly, all QTLs we identified – three in the CAC110 RIL and one in the CAC415 RIL – were additive with *M. nasutus* alleles increasing fertility (Table 3).

## Discussion

Closely related species living in secondary sympatry, especially those displaying incomplete reproductive isolation and still experiencing gene flow, provide an opportunity to investigate the evolution and maintenance of reproductive barriers in nature. Here we construct three RIL populations in one such system: hybridizing sister species *Mimulus guttatus* and *M. nasutus* from the sympatric site Catherine Creek. Our multi-parent crossing design allows for a uniquely powerful assessment of postzygotic barriers in a naturally admixed and highly genetically diverse population. Overall, patterns of TRD and long-range LD, along with interacting QTLs for hybrid sterility, provide evidence of multiple segregating hybrid incompatibilities between these naturally hybridizing species. Additionally, these RILs are now a permanent resource to be used in research on adaptation and speciation in *Mimulus*, facilitating future experiments to identify the genes involved in hybrid incompatibilities and understand their ecological context across environmental conditions.

### Introgression in sympatry: M. nasutus ancestry in CAC M. guttatus

Consistent with previous work showing signatures of ongoing and historical introgression at Catherine Creek (Brandvain *et al*., 2014; Kenney & Sweigart, 2016), we identified many clear tracks of admixture from *M. nasutus* across the genomes of the *M. guttatus* parental lines (despite having chosen these lines for their species-diagnostic traits). Despite pervasive admixture, these studies also showed a genome-wide, negative correlation between recombination rate and divergence between species at Catherine Creek (Brandvain *et al*., 2014; Kenney & Sweigart, 2016). This same negative correlation has been discovered in several other systems (Begun & Aquadro, 1993; Takahashi *et al*., 2004; Keinan & Reich, 2010) and is evidence that, in general, natural selection acts against *M. nasutus* ancestry in *M. guttatus* genetic backgrounds. Consistent with this idea, many regions of introgression in the three *M. guttatus* are unique to a single line. Despite this general pattern, two regions of introgression, at the ends of chromosomes 3 and 4, are present in all three *M. guttatus* lines. These same two regions also show elevated levels of gene flow (relative to the genome-wide average) in wild admixed individuals at CAC (Figure 4, Kenney & Sweigart, 2016) suggesting they might carry loci that are universally adaptive (Suarez-Gonzalez *et al*., 2018) or that spread through selfish processes (Lindholm & Price, 2016).

A key question at Catherine Creek is whether introgression is continuously renewed through recent hybridization or instead reflects the remnants of past events. Nearly one-quarter of the CAC110 genome shows evidence of *M. nasutus* ancestry, the expectation for a second-generation (backcross) hybrid. Although this line does contain large genomic regions with primarily *M. nasutus* ancestry (Figure S4), the largest actual block of introgression detected by the HMM is only ∼1.1 Mb, much smaller than what is expected in an early hybrid. In some cases, there might be technical reasons for these breaks in introgression – genotyping errors, for instance, or underestimates by the HMM (see Brandvain *et al*., 2014, Text S1). However, we note that use of variants in these regions largely improves the RIL genetic maps, indicating that the breaks might often have biological causes. For example, crosses between hybrid individuals might combine previously independent admixture events into larger, nearly contiguous blocks. At Catherine Creek, we find evidence of a stable admixed population in certain microsites over a period of more than 10 years (Farnitano et al., unpublished results), suggesting such mating events might be quite frequent. Additionally, some of these apparent breaks in introgression could be due to variation in *M. nasutus*, which has an average nucleotide diversity of ∼1% (Brandvain et al. 2014). Although most of this *M. nasutus* genetic variation is likely segregating between rather than within populations (Sweigart & Willis, 2003), the long history of introgression at Catherine Creek (Brandvain *et al*., 2014; Kenney & Sweigart, 2016) might inflate diversity between contemporary and remnants of past CAC *M. nasutus* samples (the latter contained in introgressions). Finally, although introgression at Catherine Creek is highly asymmetric, interspecific gene flow does occasionally go in the opposite direction, introducing *M. guttatus* alleles into *M. nasutus* (Brandvain *et al*., 2014). Such variation in *M. nasutus* could also contribute to small blocks of *M. guttatus* ancestry identified within larger blocks of introgression.

### M. nasutus as a source of structural variation in sympatric M. guttatus

From this small sample of only three *M. guttatus* lines, we identified three previously unknown structural variants segregating within *M. guttatus* at CAC, two of which appear to have been introduced into *M. guttatus* from *M. nasutus.* Unlike reciprocal translocations, which can cause F1 fertility problems through direct impacts on chromosomal segregation (Stathos & Fishman, 2014; Sotola *et al*., 2023), inversions often do not directly induce underdominant hybrid sterility in plants (Fishman & Sweigart, 2018; Huang & Rieseberg, 2020; Zhang *et al*., 2021). Nevertheless, loci for reproductive isolation have often been shown to map to inversions (Noor *et al*., 2001a,b; Livingstone & Rieseberg, 2004; Huang & Rieseberg, 2020; Todesco *et al*., 2020), potentially because, in species still connected by some degree of gene flow, they prevent selection from purging incompatibilities (Noor et al. 2001b) or lock together multiple linked variants involved in divergent adaptation (Rieseberg et al. 2001, Kirkpatrick & Barton 2006). Although none of the hybrid sterility QTLs we discovered map to any of the three inversions, several putative incompatibilities do (Table 2, Figure S6), suggesting structural variation might play a role in maintaining postzygotic isolation at Catherine Creek. Going forward, it will also be important to determine whether traits involved in divergent ecological adaptation between CAC *M. nasutus* and *M. guttatus* (Mantel & Sweigart, 2019) map to any of these inversions as has been shown in ecotypes of *M. guttatus* (Lowry & Willis 2010) and sunflowers (Todesco et al. 2020).

Whatever their functional and fitness effects may be, the three inversions are likely to have a large impact on patterns of admixture at Catherine Creek. In the most admixed of the three *M. guttatus* parents (CAC110), roughly one-quarter of the introgression is within the boundaries of the *inv1.2* and *inv13* inversions (Figure S2, Figure S6). An open question is whether inverted regions display elevated levels of divergence as expected if they disproportionately contribute to reproductive isolation and adaptive divergence (Noor et al. 2001, Rieseberg et al. 2001). If so, taking these regions into account might actually strengthen the previously discovered negative relationship between absolute divergence and local recombination (Brandvain *et al*., 2014; Kenney & Sweigart, 2016), which relied on estimates of recombination from genetic maps between non-CAC accessions.

### Variation in transmission ratio distortion and postzygotic isolation across experimental hybrid populations

Previous comparisons of TRD in *Mimulus* crosses between a common *M. guttatus* parent (the annual line IM62) and either an allopatric *M. nasutus* line (SF) or a perennial *M. guttatus* line (DUN) showed strong, but largely non-overlapping patterns of TRD (Hall & Willis 2005). In this study, too, we find that patterns of TRD are highly variable across the three RIL populations, although differences here can be attributed only to diversity within CAC *M. guttatus*. It is well known that individual populations of *M. guttatus* contain a tremendous amount of genetic variation (Flagel *et al*., 2014; Puzey *et al*., 2017) often with important phenotypic effects (Willis, 1992; Mojica *et al*., 2012; Monnahan & Kelly, 2017). This study suggests this variation might also have profound effects on reproductive barriers between species.

Although TRD may arise through several mechanisms (Fishman & McIntosh 2019), only some of them are expected to produce the polymorphism we observe among RIL populations. A consistent signal of TRD is expected in regions affected by purifying selection against weakly deleterious *M. nasutus* alleles (i.e., hybridization load: Moran *et al*., 2021) or by environmental selection favoring *M. nasutus* alleles at loci responsive to the greenhouse growth conditions. On the other hand, environmental selection favoring *M. guttatus* alleles (including epistatic selection – see Table 2) will produce a consistent signal only if all three CAC *M. guttatus* parents share the same functional alleles at causal loci. Similarly, inbreeding depression in *M. guttatus* might be expected to produce variable patterns of TRD given that it is often caused by rare deleterious alleles (Willis, 1999a,b). Because meiotic drivers are generally expected to fix rapidly in the absence of associated fitness costs (Burt & Trivers, 1998), they might seem unlikely to contribute to variable TRD. However, one such well-characterized driver – the centromeric *D* allele in *M. guttatus* – is polymorphic in the Iron Mountain population where it was discovered, and linked to alleles that reduce ferility (Fishman & Saunders, 2008). Thus, even selfish segregation distorters might contribute to the variable patterns of TRD we observe in the CAC RILs.

Another well-known cause of distortion – and the one with the most relevance for speciation – is the reduced viability of gametes or zygotes due to hybrid incompatibilities (Leppälä *et al*., 2013; Kerwin & Sweigart, 2017). During RIL construction, gametic incompatibilities may be particularly likely to persist, as even RILs with fewer viable ovules or pollen grains can be successfully self-fertilized. Gametic incompatibilities affecting male and female fertility have been identified in crosses between *M. guttatus* and *M. nasutus* (Kerwin & Sweigart 2017) and between *M. parishii* and *M. cardinalis* (Sotola *et al*., 2023), and the genotypic ratios observed at many putative two-locus incompatibilities at CAC (Table 2) are consistent with this mechanism of action. On the other hand, given that QTLs associated with reductions in pollen viability (Table 3) do not map to distorted regions, it seems likely that these incompatibilities disrupt male fertility in the diploid sporophyte. Whatever their mechanism, a key point for this study is that hybrid incompatibilities between closely related species are often polymorphic (López-Fernández & Bolnick, 2007; Cutter, 2012; Corbett-Detig *et al*., 2013; Charron *et al*., 2014; Servedio & Hermisson, 2020). Moreover, in *Mimulus*, hybrid incompatibility loci often vary *within* populations (Christie & Macnair, 1987; Sweigart *et al*., 2007; Sweigart & Flagel, 2015; Zuellig & Sweigart, 2018a). In one case of duplicate genes that cause hybrid seedling inviability between sympatric *M. guttatus* and *M. nasutus* at Don Pedro Reservoir (DPR) in California (Zuellig & Sweigart, 2018b), at least some of the polymorphism seems to be due to introgression from *M. nasutus* into *M. guttatus* (Zuellig & Sweigart, 2018a). Given the large amount of admixture observed in CAC *M. guttatus*, it seems highly probable that, like at DPR, interspecific gene flow contributes to polymorphism in hybrid incompatibilities at Catherine Creek.

Despite this general variability, there is one region at the end of chromosome 14 consistently distorted in all three RIL populations (with an excess of *M. guttatus* alleles), which may point to a conserved mechanism. This same region actually showed the opposite pattern of distortion (an excess of *M. nasutus* alleles) in F_2_ hybrids between IM62 *M. guttatus* and SF *M. nasutus* (Fishman *et al*., 2001, 2008), suggesting the underlying mechanism is not shared species-wide and might be unique to CAC.

## Conclusions

Overall, these RIL populations provide evidence of diverse contributors to postzygotic isolation between *Mimulus* species, suggesting long-term persistence of hybrids at Catherine Creek may be unlikely. Even highly admixed *M. guttatus* individuals carry alleles at multiple loci that are incompatible with *M. nasutus*. This substantial load of hybrid incompatibilities may help explain the observed cline of *M. nasutus* introgression in wild admixed individuals at Catherine Creek (Kenney & Sweigart, 2016), which seems to suggest persistent backcrossing to *M. guttatus* and purging of *M. nasutus* ancestry. Moreover, our findings of genome-wide TRD and several retained fertility QTLs in the RILs supports previous evidence for broad, genome-wide selection against *M. nasutus* ancestry in an *M. guttatus* background (Brandvain *et al*., 2014; Kenney & Sweigart, 2016). Thus, despite a long history of admixture at Catherine Creek, intrinsic barriers between *Mimulus* species remain pervasive, limiting the homogenizing effects of gene flow and, potentially, playing a key role in the maintenance of species.

## Author Contributions

Research conceived and designed by SJM and ALS. Data collected and analyzed by SJM. Manuscript written by SJM and ALS.

## Data Accessibility

All whole genome sequence data of inbred lines and RADseq data of RILs will be deposited at the NCBI Sequence Read Archive upon acceptance. Other data and scripts will be made available from the Dryad Digital Repository.

## Acknowledgements

We thank Amanda Kenney for seed collection at Catherine Creek. We are also grateful to Aaliyah Stoudemire, Violet Iyahne, Suni Thakur, Brennen Anderson, and Selma Kajtazovic for assistance with RIL creation, and Michaela Yarbough, Inam Jameel, and Mohamed Elgallad for help with RIL crossing and fertility phenotyping. We thank greenhouse staff, Greg Cousins, Kevin Tanner, Noah Hentherinton, and Mike Boyd for plant care. Jill Anderson, Lisa Donovan, Bob Schmitz, and Mike White provided helpful discussions and guidance. Makenzie Whitener, Matt Farnitano, and Natalie Gonzalez provided valuable comments on a previous draft. This work was supported by a National Science Foundation grant [IOS-1827645 to ALS]; [DEB-1856223 to ALS]; funds from the University of Georgia Research Foundation [to ALS]; the National Institute of General Medical Sciences of the National Institute of Health [T32GM007103 to SJM]; and the UGA Genetics Jan and Kirby Alton Graduate Fellowship [to SJM].

