## Supplemental Methods, Figures, and Tables for "Postzygotic barriers persist despite ongoing introgression in hybridizing *Mimulus* species"

**Supporting Information**

Methods S1

*Plant Growth Conditions*

We planted seeds onto moist Fafard 3-B potting mix (Sun Gro Horticulture, Agawam, MA) in 2.5” pots, and cold stratified them for seven days at 4°C before moving them to a Conviron growth chamber (16-hr days; 23°C days/16°C nights) for germination. We transplanted seedlings into individual cells of 96-cell plug trays filled with potting mix and moved them to the University of Georgia greenhouses under 16-hr supplemental light, 23°C days/16°C nights.

*BestRAD Library Preparation*

Briefly, we digested 50-100 ng of genomic DNA with BfaI and PstI, and used the ‘NEBNext Ultra II’ library preparation kits for Illumina (New England BioLabs, Ipswich, MA) to ligate 48 unique barcoded i5 adaptors to each sample, isolated tagged fragments, and pooled samples into 8 pools of 48 or fewer samples. We attached a unique ‘NEBNext Adaptor for Illumina’ i7 adaptor to each pool of samples, and enriched the samples with PCR. We size-selected the libraries to 300-900bp using BluePippin 2% agarose cassettes (Sage Science, Beverly, MA) and pooled the libraries in equimolar concentrations following quantification via qPCR. (dx.doi.org/10.17504/protocols.io.6awhafe).

*Processing of Sequence Data*

After trimming Illumina adaptors from reads using Trimmomatic (v0.36; Bolger *et al.*, 2014), we aligned WGS reads to the *Mimulus guttatus* IM62 v3 refence assembly (phytozome.org, IM: Iron Mountain, an allopatric *M. guttatus* population in Oregon, ~200km south of CAC) using BWA-MEM (v0.7.17; Li & Durbin, 2009; Li, 2013). We then removed reads with alignment qualities <Q29 with SAMtools view (v1.6; Li *et al.*, 2009). Using Picard (v2.16.0; broadinstitute.github.io/picard/), we added read groups, checked mate-pair information, and removed PCR duplicates. We used SAMtools view (v0.7.17; Li *et al.*, 2009) to remove paired reads mapped to different contigs. Finally, using GATK HaplotypeCaller (v3.8.1; Van der Auwera & O'Connor, 2020) in GVCF mode, we called and filtered SNPs, removing heterozygous calls and retaining sites with a minimum depth of 3x, and a maximum depth of 2 standard deviations above the mean (calculated with Qualimap2 v2.2.1; Okonechnikov *et al.*, 2016) for each line. To aid in genotyping *M. guttatus* and *M. nasutus* SNPs across the genome, we identified a full list of diagnostic SNPs (i.e., those homozygous for different alleles in each of the parental pairs, CAC9 & CAC110, CAC9 & CAC162, and CAC9 & CAC415). We joint genotyped each pair of parental lines using GATK GenotypeGVCFs (v3.8.1; Van der Auwera & O'Connor, 2020), quality filtered the called SNPs (QD < 2.0, FS > 60.0, MQ < 40.0, MQRankSum < -12.5, ReadPosRankSum < -8.0), and retained only biallelic SNPs using VariantFiltration and SelectVariants. From these three VCFs, we extracted a list of high-quality, diagnostic SNPs with VCFtools (v0.1.16; Danecek *et al.*, 2011).

Following sequencing, we demultiplexed the ddRADseq data by i7 index and used a custom Python script (dx.doi.org/10.17504/protocols.io.bjnbkman) to remove PCR duplicates from each library using the unique molecular barcode of the i5 adaptor. We demultiplexed each library to individual samples using Flip2BeRAD (github.com/tylerhether/Flip2BeRAD) and Stacks process_radtags (v2.55; Catchen *et al.*, 2011, 2013). Next, we removed Illumina adaptors and aligned reads to the *Mimulus guttatus* IM62 v3 reference assembly. At this stage, six samples were removed from further analysis (4 from the CAC162 RILs, and 2 from the CAC415 RILs) due to low similarity between duplicate BAMs produced from the two independent ddRAD library preparations. We then merged the remaining duplicate BAMs with SAMtools merge (v0.7.17; Li *et al.*, 2009) for a total of 362 RIL individuals (151 CAC110 RILs, 124 CAC162 RILs, and 87 CAC415 RILs). We used Picard and SAMtools to process reads as above (with the exception of MarkDuplicates, as PCR duplicates were already removed). We called SNPs, joint genotyped each RIL population with its corresponding parental lines, and filtered called SNPs as described above. Finally, we used VariantFiltration and SelectVariants in GATK (v3.8.1; Van der Auwera & O'Connor, 2020) to remove sites with only parental coverage and used VCFtools (v0.1.16; Danecek *et al.*, 2011) to remove non-diagnostic sites between each pair of parental lines.

*Linkage Mapping*

We used LepMAP3 (Rastas, 2017) to construct a linkage map for each of the three RIL populations. We used the affx2post.awk script to calculate posterior genotype probabilities, and then ran ParentCall2 with default parameters. We ran SeparateChromosomes2 to group markers from the same chromosome, followed by OrderMarkers2 (grandparentPhase=1, useKosambi=1, sexAveraged=1, 10 iterations/per chromosome) to create a linkage map for each chromosome of the reference genome (details in Table S2). Of the 10 iterations of OrderMarkers2, (or using the physical order of markers in the reference genome if order was poorly resolved), we selected the order with the highest likelihood for each chromosome (Table

S2). Finally, we manually removed standalone markers whose physical position differed greatly from their calculated genetic positions.

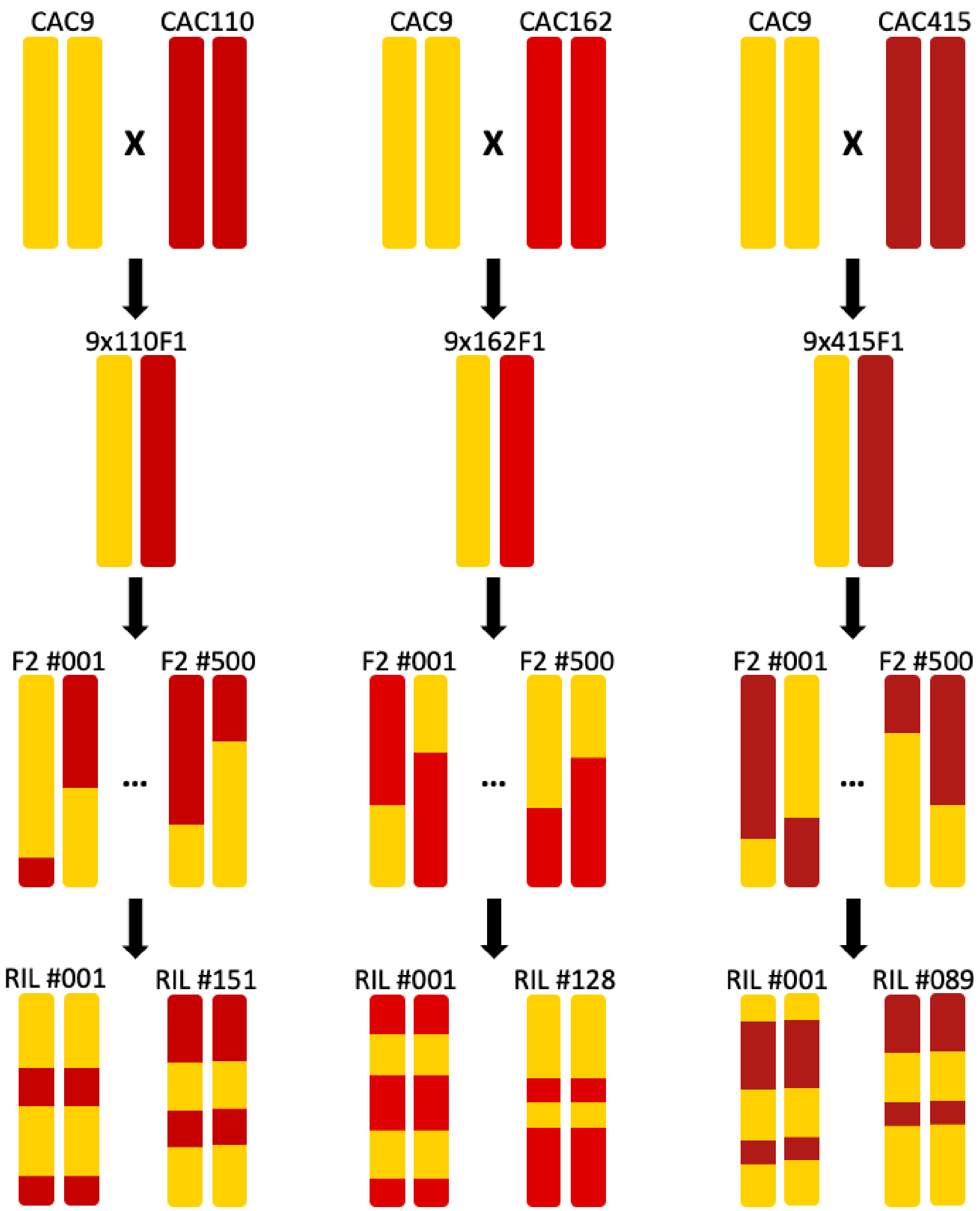


**Figure S1. Recombinant inbred line (RIL) population crossing designs.** We constructed three parallel RIL populations by crossing maternal *M. nasutus* line CAC9 to three paternal *M. guttatus* lines: CAC110, CAC162, and CAC415. We self-pollinated one F1 from each cross, producing a population of 500 F2s. We self-pollinated each F2 for at least 4 more generations with single seed decent to produce three RIL populations of F6 (or later generation) inbred individuals.

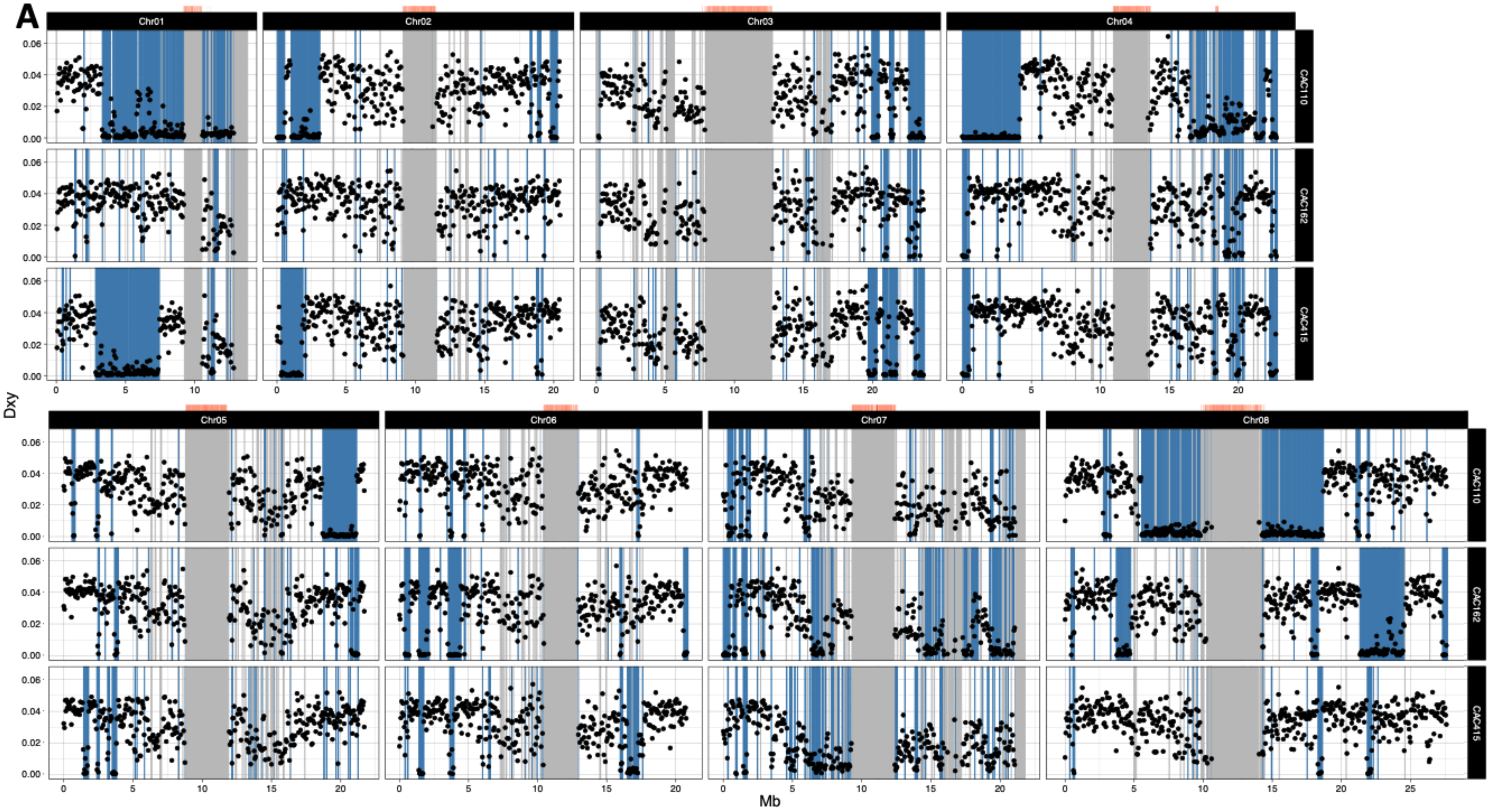
A
Chr01
Chr02
Chr03
Chr04
Chr05
Chr06
Chr07
Chr08
CAC110
CAC162
CAC415
Dxy
Mb

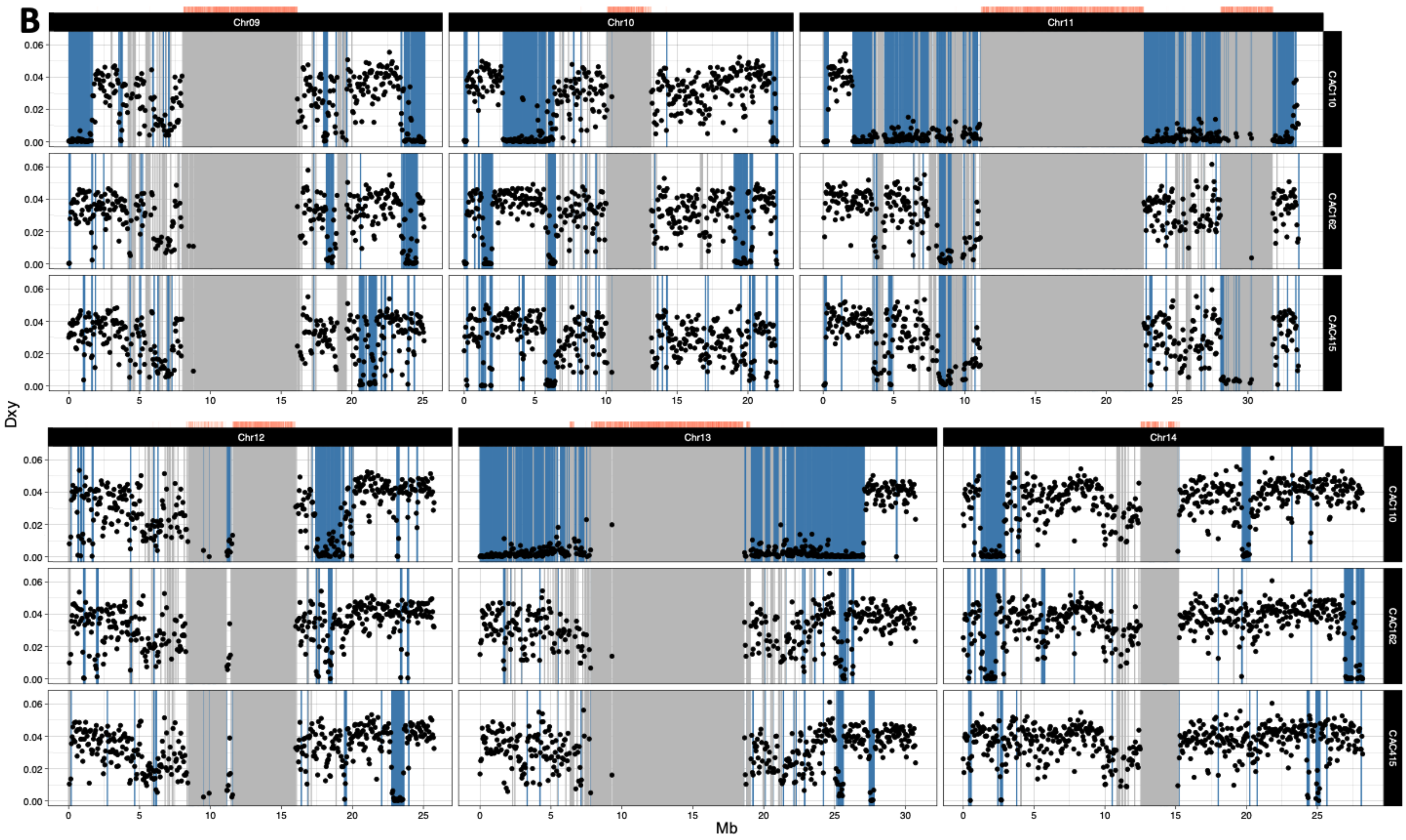


**Figure S2. Introgression from *M. nasutus* into sympatric *M. guttatus*. (a) Chromosome (Chr) 01-08, (b) Chr09-Chr14.** Dxy calculated in 50kb windows across each chromosome of the IM62 v3 *Mimulus guttatus* reference genome, between the parental lines of the three RIL populations (CAC9 v. CAC110, CAC9 v. CAC162, and CAC9 v. CAC415). Blue shaded regions indicate putatively admixed regions of the three focal CAC *M. guttatus* lines tested, diagnosed by a greater than a 95% posterior probability of *M. nasutus* ancestry (in 1kb windows) as inferred via the HMM (i.e. more recent coalescence with included *M. nasutus* samples than with included *M. guttatus* samples). Grey shaded regions indicate missing data (fewer than 5000 genotyped sites within any 50kb window). Red heatmaps above each chromosome indicate the density of centromeric repeats in the same 50kb windows.

| | Chr01 | Chr02 | Chr03 | Chr04 | Chr05 | Chr06 | Chr07 | Chr08 | Chr09 | Chr10 | Chr11 | Chr12 | Chr13 | Chr14 | Genome |
|---|---|---|---|---|---|---|---|---|---|---|---|---|---|---|---|
| CAC6 | 3.05% | 14.40% | 8.65% | 23.31% | 10.22% | 12.11% | 10.09% | 7.39% | 10.24% | 8.95% | 43.70% | 10.73% | 3.99% | 2.62% | 12.90% |
| CAC110 | 46.18% | 18.90% | 11.93% | 32.88% | 14.65% | 5.31% | 7.92% | 32.65% | 14.82% | 17.93% | 54.44% | 15.10% | 40.03% | 10.13% | 23.83% |
| CAC112 | 3.29% | 21.46% | 15.58% | 23.00% | 14.15% | 27.49% | 10.61% | 44.21% | 8.28% | 46.45% | 35.22% | 6.99% | 3.18% | 14.91% | 20.17% |
| CAC134 | 4.44% | 5.40% | 8.24% | 8.03% | 4.57% | 8.63% | 14.42% | 7.49% | 16.61% | 10.35% | 15.82% | 3.57% | 2.46% | 11.48% | 8.92% |
| CAC141 | 3.67% | 5.55% | 5.80% | 7.58% | 8.64% | 9.75% | 20.69% | 10.06% | 9.93% | 13.41% | 9.65% | 7.52% | 2.30% | 9.38% | 8.85% |
| CAC162 | 4.68% | 3.25% | 6.33% | 8.73% | 6.41% | 13.48% | 23.40% | 18.54% | 6.07% | 12.56% | 9.89% | 4.78% | 1.94% | 9.15% | 9.24% |
| CAC262 | 47.84% | 10.07% | 17.30% | 10.48% | 3.88% | 21.00% | 12.26% | 44.58% | 10.21% | 7.21% | 29.25% | 51.32% | 20.80% | 6.07% | 20.90% |
| CAC415 | 32.17% | 10.67% | 11.46% | 8.78% | 5.86% | 8.15% | 9.72% | 4.82% | 5.03% | 8.55% | 14.22% | 7.05% | 2.54% | 2.94% | 8.61% |

**Figure S3.** Heatmap showing the proportion of *M. nasutus* ancestry in each chromosome, and across the genome, of each of the CAC *M. guttatus* lines, estimated as the proportion of 1kb windows with a greater than a 95% posterior probability of *M. nasutus* ancestry as inferred via the HMM. Darker blue shading indicates a higher proportion of *M. nasutus* ancestry.

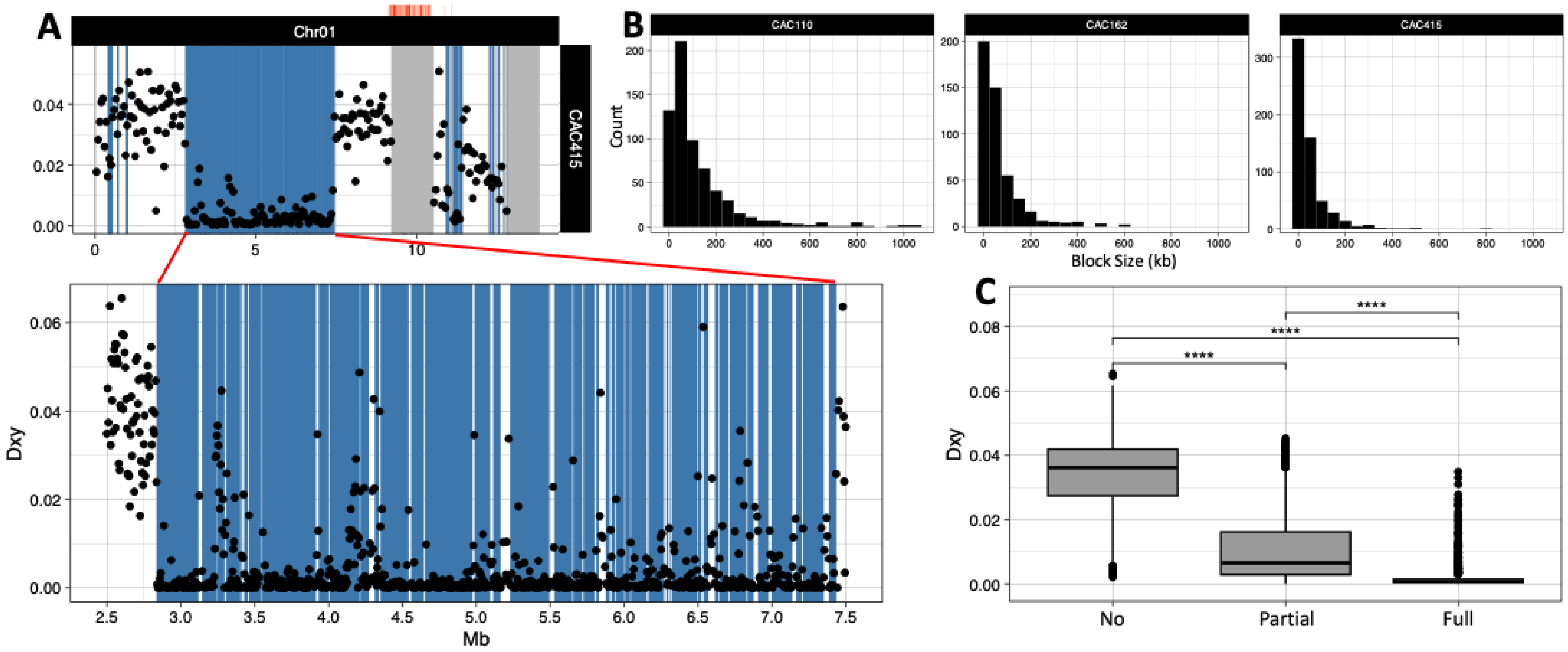


**Figure S4. Block size and characteristics of introgression from *M. nasutus* into sympatric *M. guttatus.* (a)** A fine scale look at diagnosed blocks of introgression on the first chromosome in sympatric *M. guttatus* line CAC415. Top: Dxy calculated over 50kb windows between CAC9 and CAC415. Grey shaded regions indicate missing data (fewer than 5000 genotyped sites within any 50kb window). Red heatmaps above the chromosome indicate the density of centromeric repeats in the same 50kb windows. Blue shaded regions indicate putatively admixed regions diagnosed by a greater than a 95% posterior probability of *M. nasutus* ancestry (in 1kb windows) as inferred via the HMM (i.e. more recent coalescence with included *M. nasutus* samples than with included *M. guttatus* samples). The shaded region between ~3 and 7.5 Mb is shown in finer detail, with Dxy calculated in 5kb windows across this region. **(b)** Histograms showing the length of each inferred block of *M. nasutus* introgression in each of the sympatric *M. guttatus* lines used as parents for the three RIL populations. CAC110 average block length: ~125kb, n=645 blocks, CAC162: ~66kb, n=476, CAC415: ~48bp, n=603. **(c)** Dxy in each 50kb window in the three sympatric *M. guttatus* lines used as parents for the three RIL populations binned by if that window has no inferred *M. nasutus* introgression within it (No), part of the window contains inferred introgression (Partial), or the entire window is covered by an inferred block of introgression (Full). Wilcox test, **** p<0.0001.

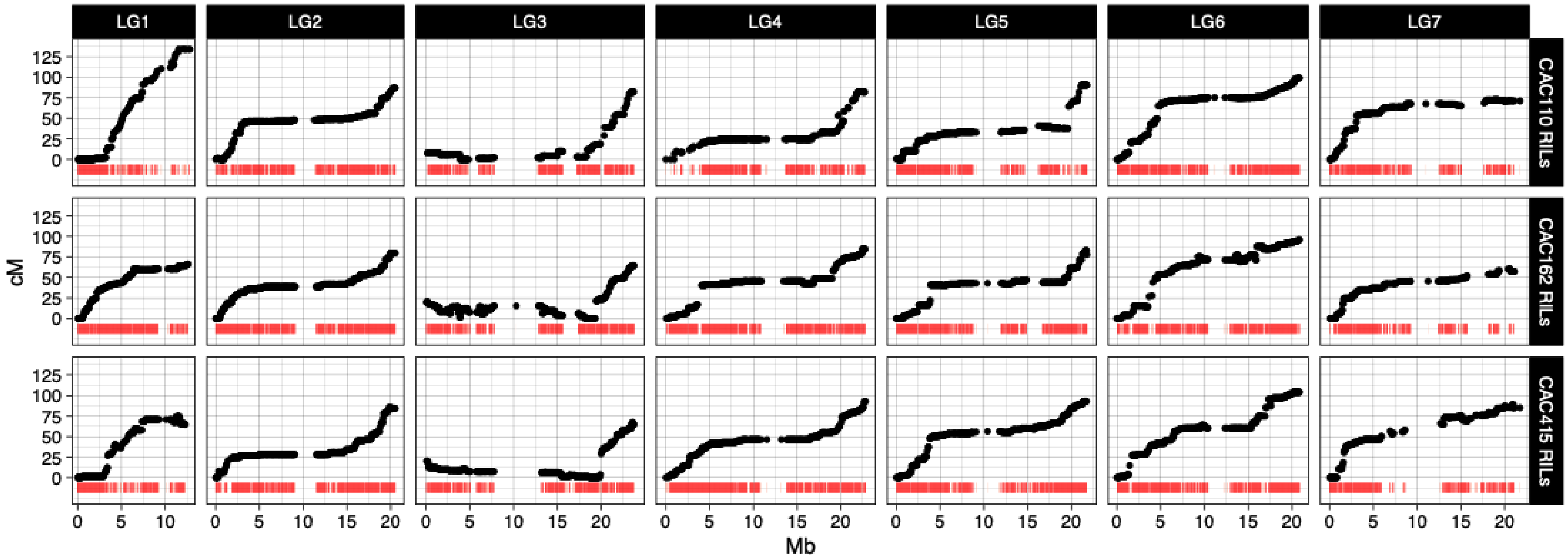

LG1
LG2
LG3
LG4
LG5
LG6
LG7
CAC110 RILs
CAC162 RILs
CAC415 RILs
cM
Mb
125
100
75
50
25
0
0
5
10
15
20

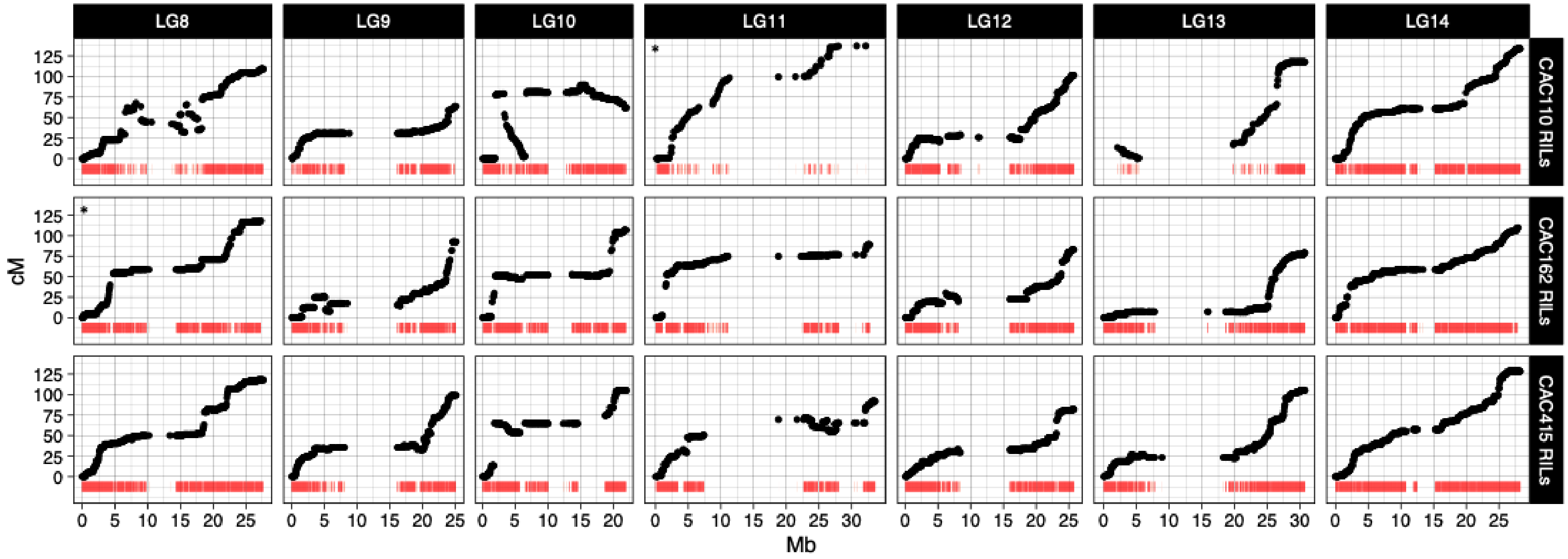


**Figure S5. Map position (cM) versus physical marker positions (Mb) of the IM62 v3 *Mimulus guttatus* reference genome in the three RIL populations. (a) Linkage group (LG) 1-LG7, (b) LG8-LG14.** Red heatmaps below each curve indicate the density of markers, measured as the distance in Mb to the nearest neighboring marker. Low SNP density, and the subsequent inability to genotype markers in areas of the *M. guttatus* parental genomes showing evidence of introgression from *M. nasutus,* manifests as regions of low marker density on some chromosomes. Recombination rate is substantially lower across the center of each chromosome, with a gap in genotyped markers that colocalize with a metacentric (dicentric in the case of Chr11) chromosomal structure, identified via a high density of centromeric repeats. Negative slope suggests inverted marker order relative to the IM62 v3 *M. guttatus* reference genome in both the *M. nasutus* and *M. guttatus* parents. Blocks of markers spanning large physical distances, but which map to nearly the same genetic position suggest inversions between the *M. guttatus* and *M. nasutus* parents used to produce that RIL population.

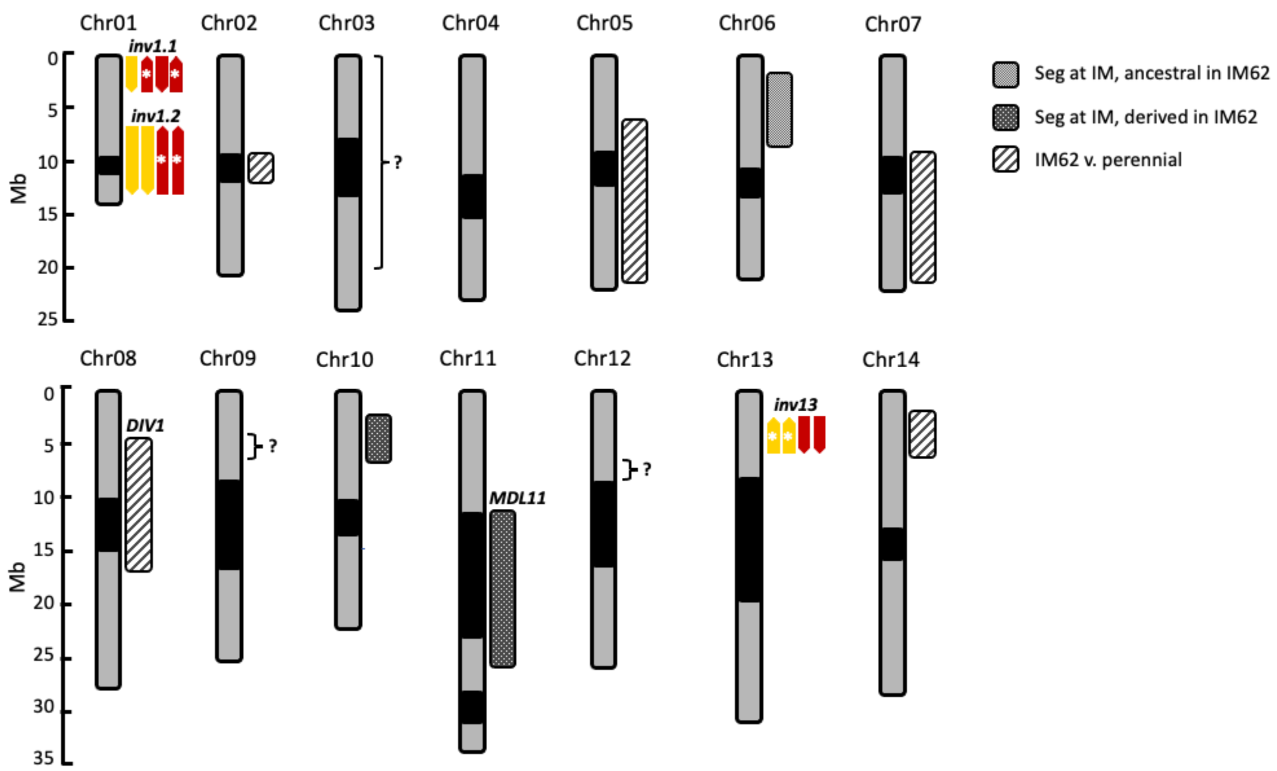


**Figure S6. Locations of newly identified and known inversions in *M. gutattus*.** Physical positions (Mb) of each of the identified putative chromosomal inversions among the CAC *M. nasutus* and *M. guttatus* lines crossed. Centromeres are indicated in black. Arrows to the right of each chromosome indicate the observed orientation of each inversion in CAC9, CAC110, CAC162, and CAC415 from left to right. Arrows in yellow show evidence of *M. nasutus* ancestry, arrows in red show evidence of *M. guttatus* ancestry. The locations of previously identified inversions in *M. guttatus* are shown to the right of each chromosome. Chr02: (Flagel *et al.*, 2019). Chr05: (Holeski *et al.*, 2014; Monnahan & Kelly, 2017; Flagel *et al.*, 2019; Kolis *et al.*, 2022). Chr06: (Lee, 2009; Fishman *et al.*, 2014; Lee *et al.*, 2016; Monnahan & Kelly, 2017). Chr07: (Flagel *et al.*, 2019). Chr08: (Lowry & Willis, 2010; Holeski *et al.*, 2014; Twyford & Friedman, 2015; Monnahan & Kelly, 2017; Flagel *et al.*, 2019; Kolis *et al.*, 2022). Chr10: (Lee, 2009; Fishman *et al.*, 2014; Holeski *et al.*, 2014; Lee *et al.*, 2016; Monnahan & Kelly, 2017; Flagel *et al.*, 2019). Chr11: (Fishman & Willis, 2005; Fishman & Saunders, 2008; Fishman & Kelly, 2015; Monnahan & Kelly, 2017; Flagel *et al.*, 2019; Finseth *et al.*, 2021). Chr14: (Flagel *et al.*, 2019). Brackets marked with question marks indicate further areas of interest, due to unique marker order, but which could not be confidently identified as inversions.

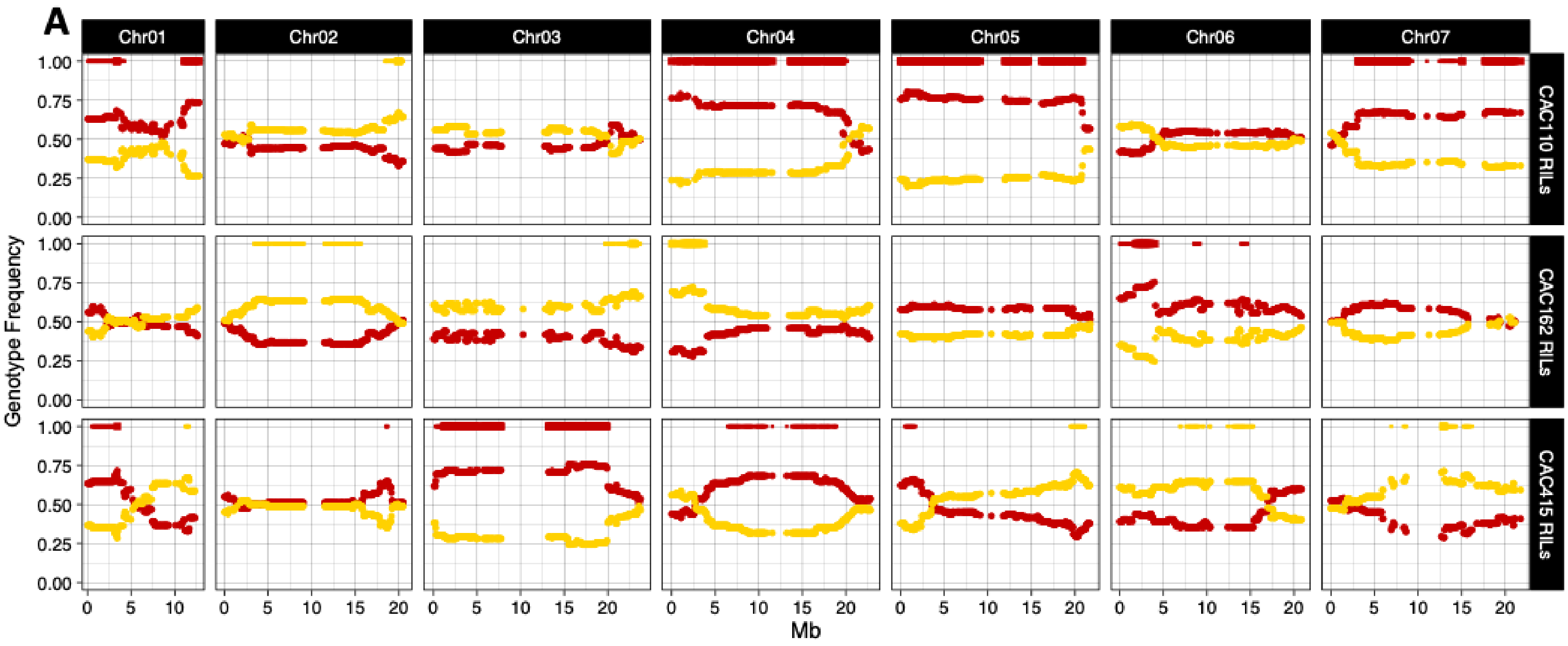

A
Chr01
Chr02
Chr03
Chr04
Chr05
Chr06
Chr07
CAC110 RILs
CAC162 RILs
CAC415 RILs
Genotype Frequency
1.00
0.75
0.50
0.25
0.00
0
5
10
15
20
Mb


183

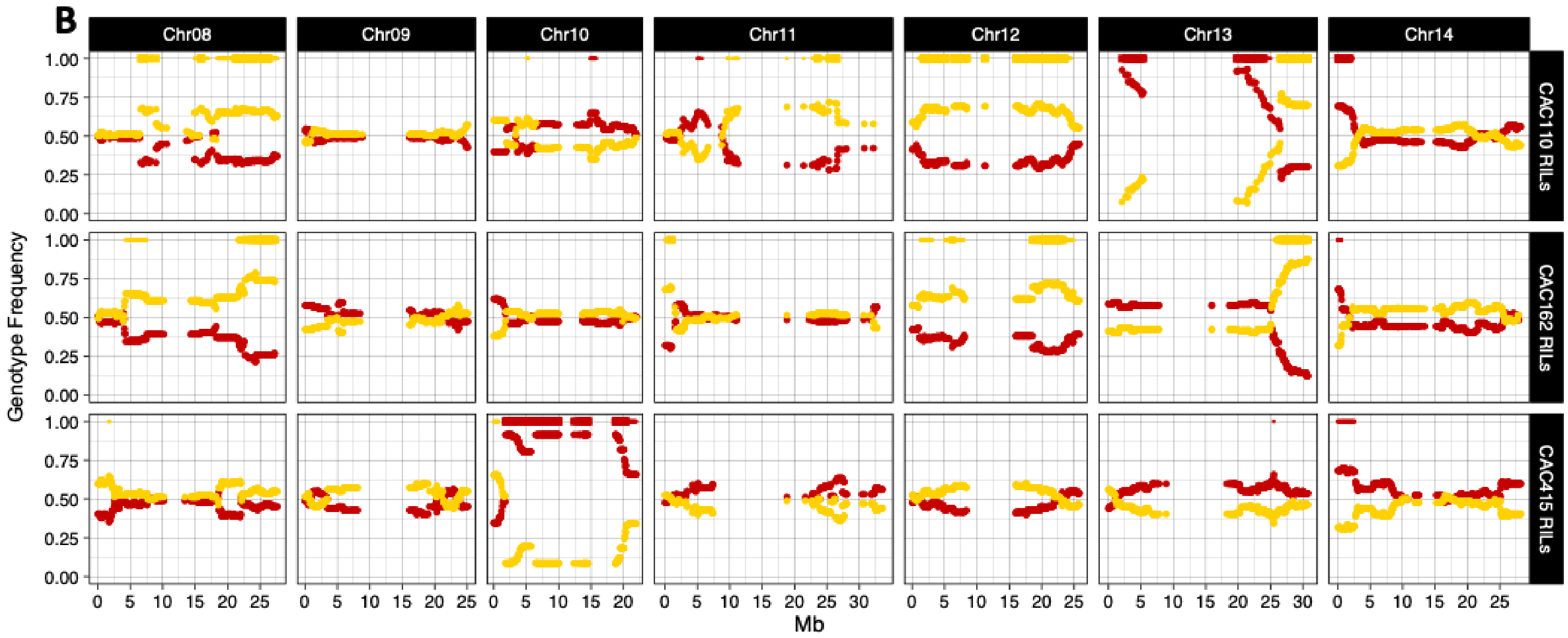


**Figure S7. Transmission ratio distortion by physical position, (a) Chromosome (Chr) 01-07, (b) Chr07-Chr14.** Genotype frequencies in windowed 50kb markers across the 14 linkage groups of the three RIL populations. Red dots indicate homozygous *M. guttatus* genotypes, yellow dots, homozygous *M. nasutus* genotypes. Bars at the top of each plot indicate regions showing significant transmission ratio distortion from the Mendelian 1:1 expectation by $\chi^2$ tests at $\alpha = 0.01$ (thin bars), and a more stringent Bonferonni-corrected $\alpha$ (thick bars, unique $\alpha$ for each chromosome in each RIL population depending on the number of markers tested, see Table S3 for values). Bar color indicates and excess of *M. guttatus* (red), or *M. nasutus* (yellow) genotypes. Markers are ordered by physical position (Mb) within the IM62 v3 *Mimulus guttatus* reference genome.

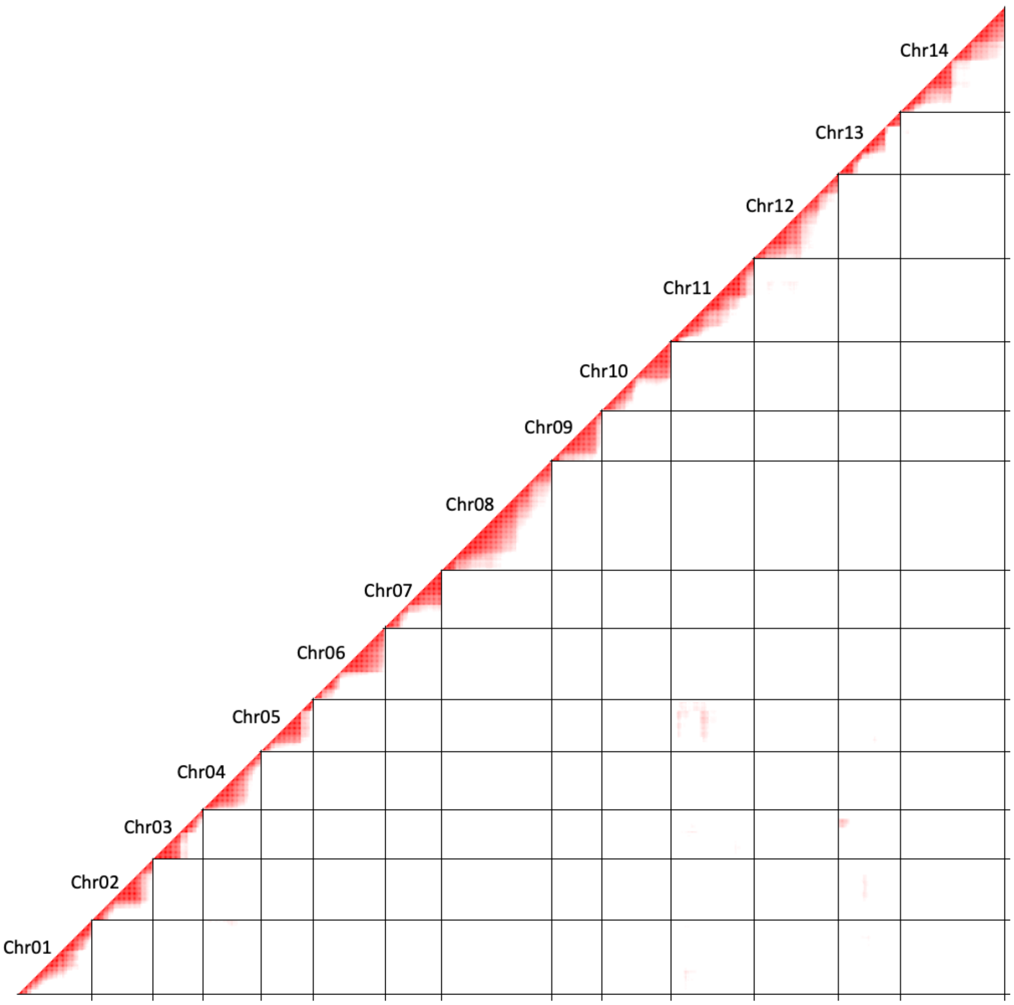


**Figure S8. CAC110 RILs linkage disequilibrium heatmap.** LD heatmap for each pairwise marker comparison across all 14 chromosomes, showing r values greater than the 95% quantile (>0.37) in the first RIL population. Values range from 0 (white) to 1 (red).

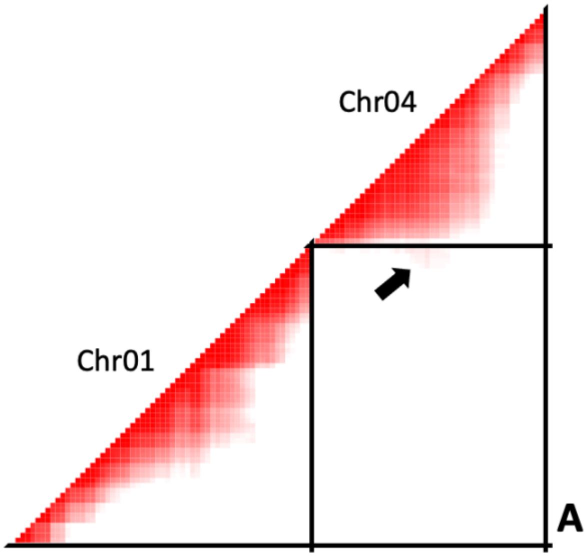


196

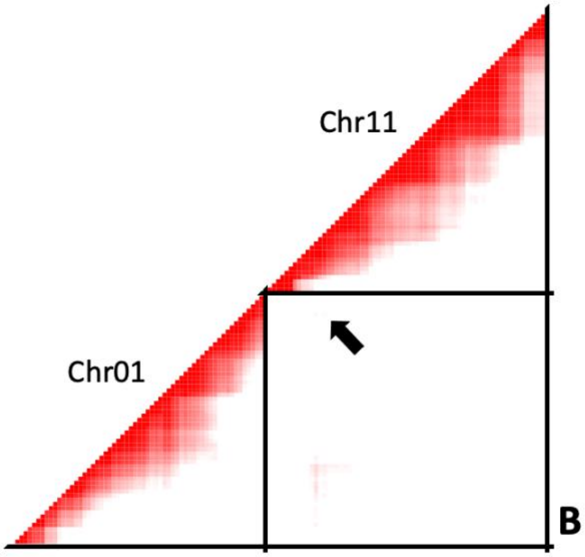


197

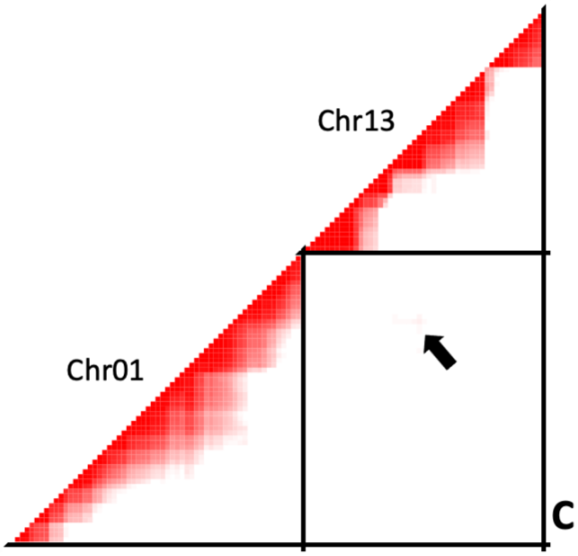
Chr13
Chr01
C

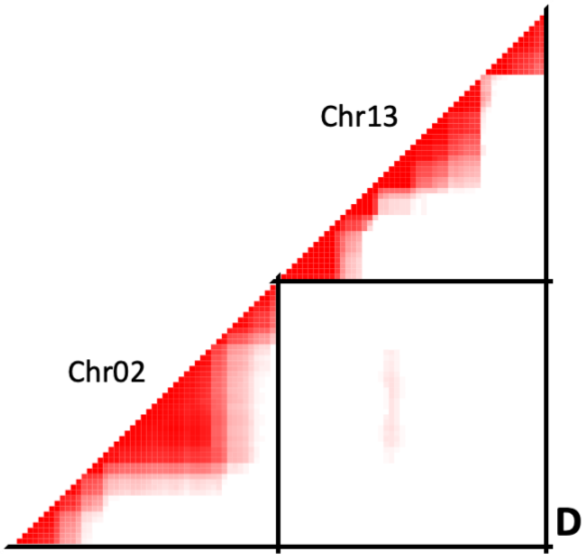
Chr13
Chr02
D

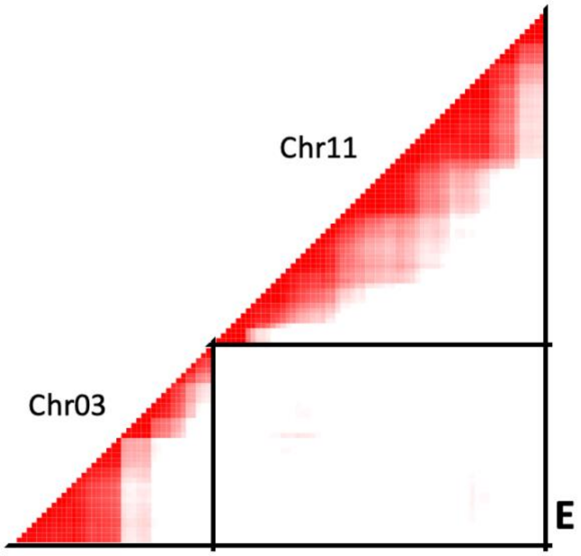
Chr11
Chr03
E

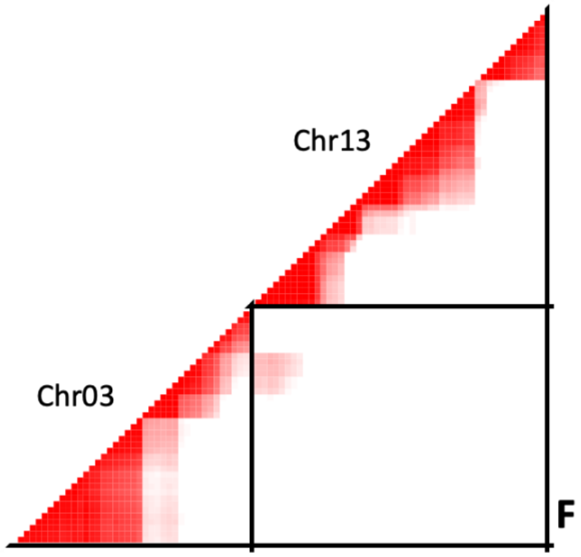
Chr13
Chr03
F

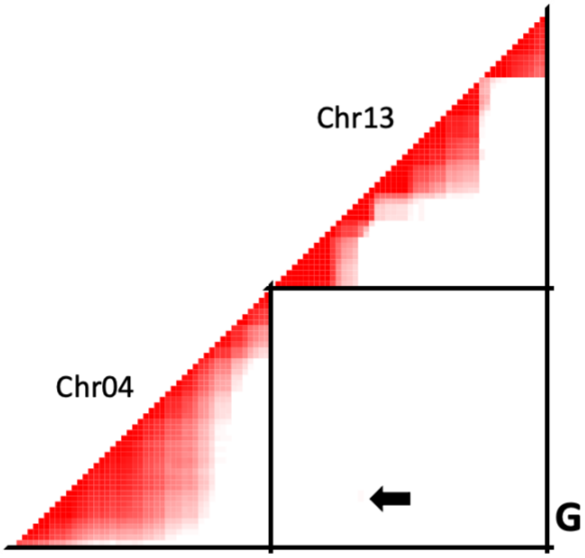


202

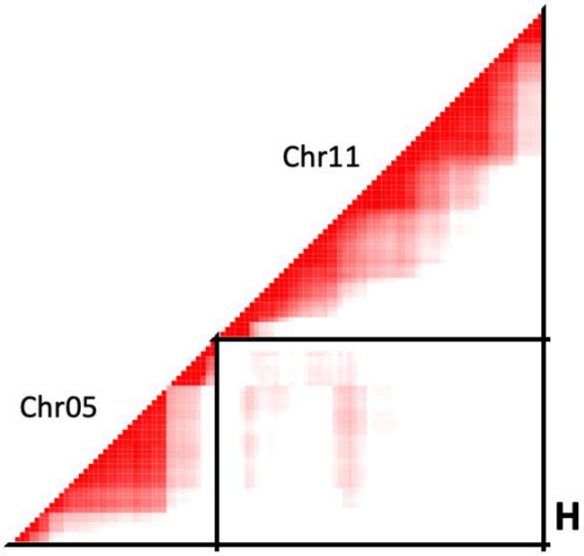


203

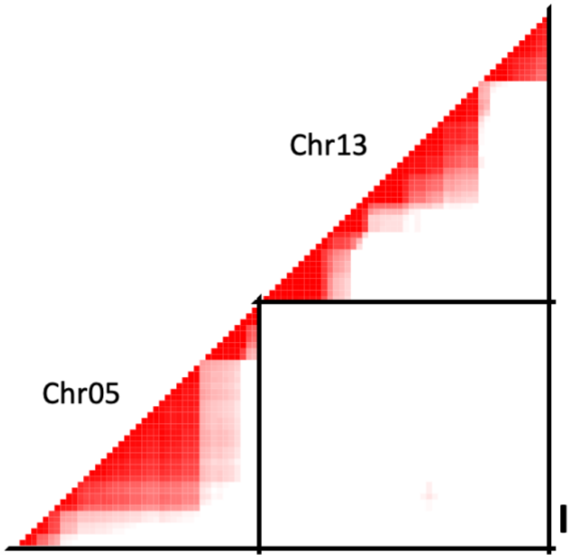
Chr13
Chr05
I

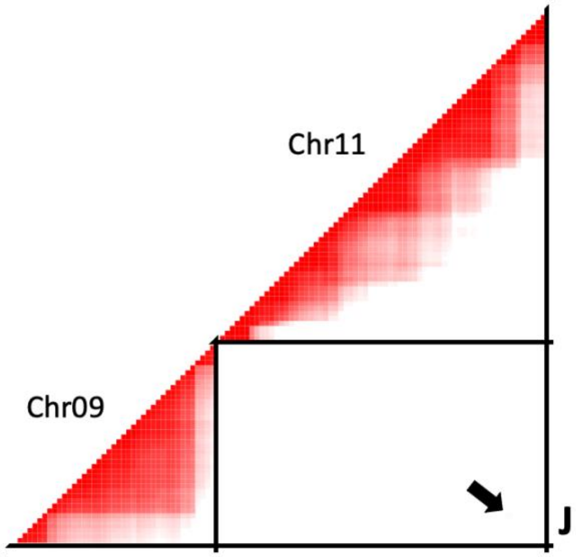
Chr11
Chr09
J

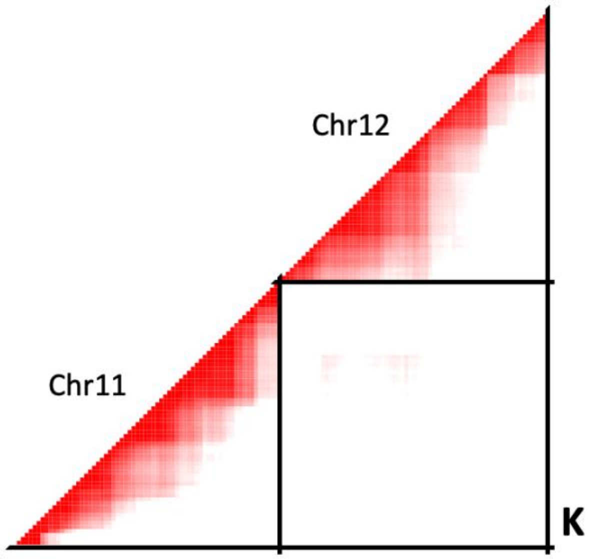


**Figure S9. CAC110 RILs linkage disequilibrium heatmap.** LD heatmaps for each pairwise marker comparison, showing r values greater than the 95% quantile (>0.37) in chromosome comparisons with at least 1 marker above the cutoff in the first RIL population. Values range from 0 (white) to 1 (red). Arrows indicates significant marker comparisons, when distorted regions are small. **(a)** Chr01 and Chr04 **(b)** Chr01 and Chr11 **(c)** Chr01 and Chr13 **(d)** Chr02 and Chr13 **(e)** Chr03 and Chr11 **(f)** Chr03 and Chr13 **(g)** Chr04 and Chr13 **(h)** Chr05 and Chr11 **(i)** Chr05 and Chr13 **(j)** Chr09 and Chr11 **(k)** Chr11 and Chr12.

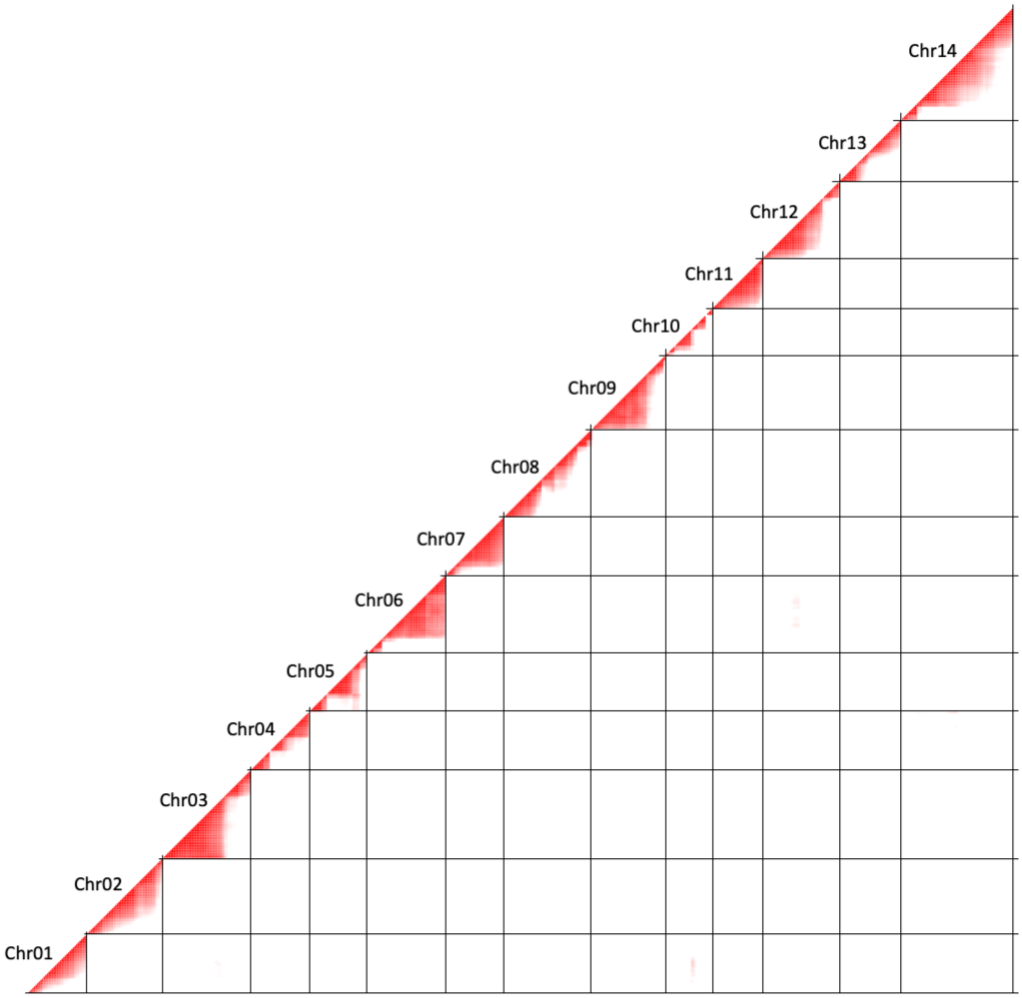


**Figure S10. CAC162 RILs linkage disequilibrium heatmap.** LD heatmap for each pairwise marker comparison across all 14 chromosomes, showing r values greater than the 95% quantile (>0.41) in the second RIL population. Values range from 0 (white) to 1 (red).

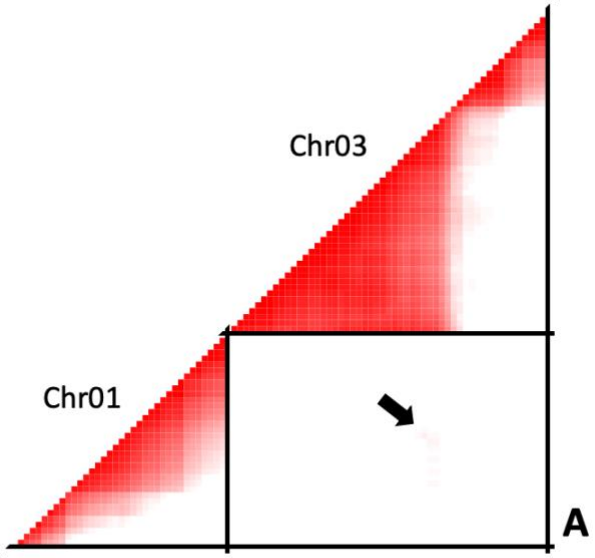
Chr03
Chr01
A

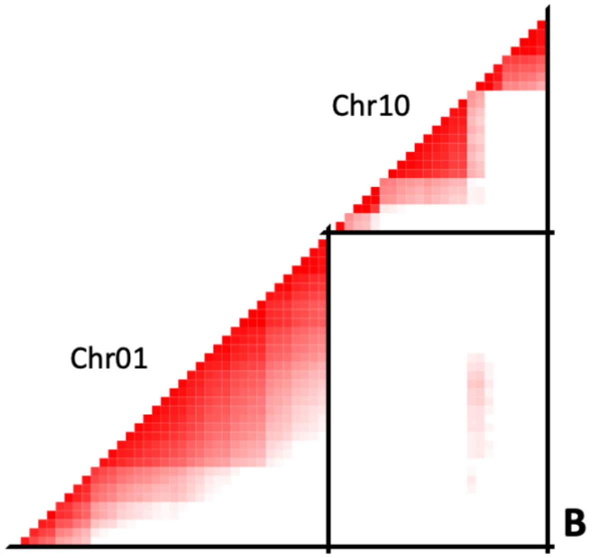
Chr10
Chr01
B

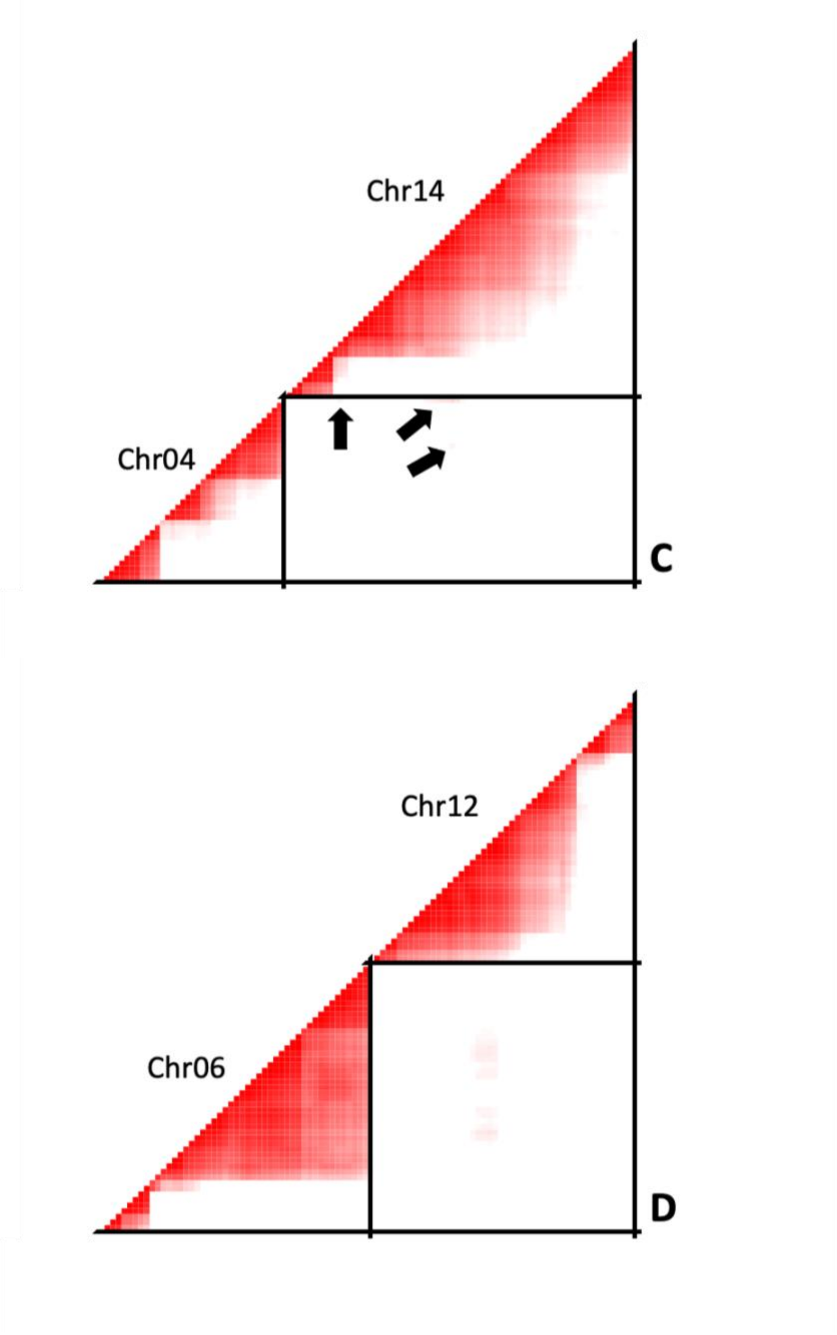


**Figure S11. CAC162 RILs linkage disequilibrium heatmap.** LD heatmaps for each pairwise marker comparison, showing r values greater than the 95% quantile (>0.41) in chromosome comparisons with at least 1 marker above the cutoff in the second RIL population. Values range from 0 (white) to 1 (red). Arrows indicates significant marker comparisons, when distorted regions are small. **(a)** Chr01 and Chr03 **(b)** Chr01 and Chr10 **(c)** Chr04 and Chr14 **(d)** Chr06 and Chr12.

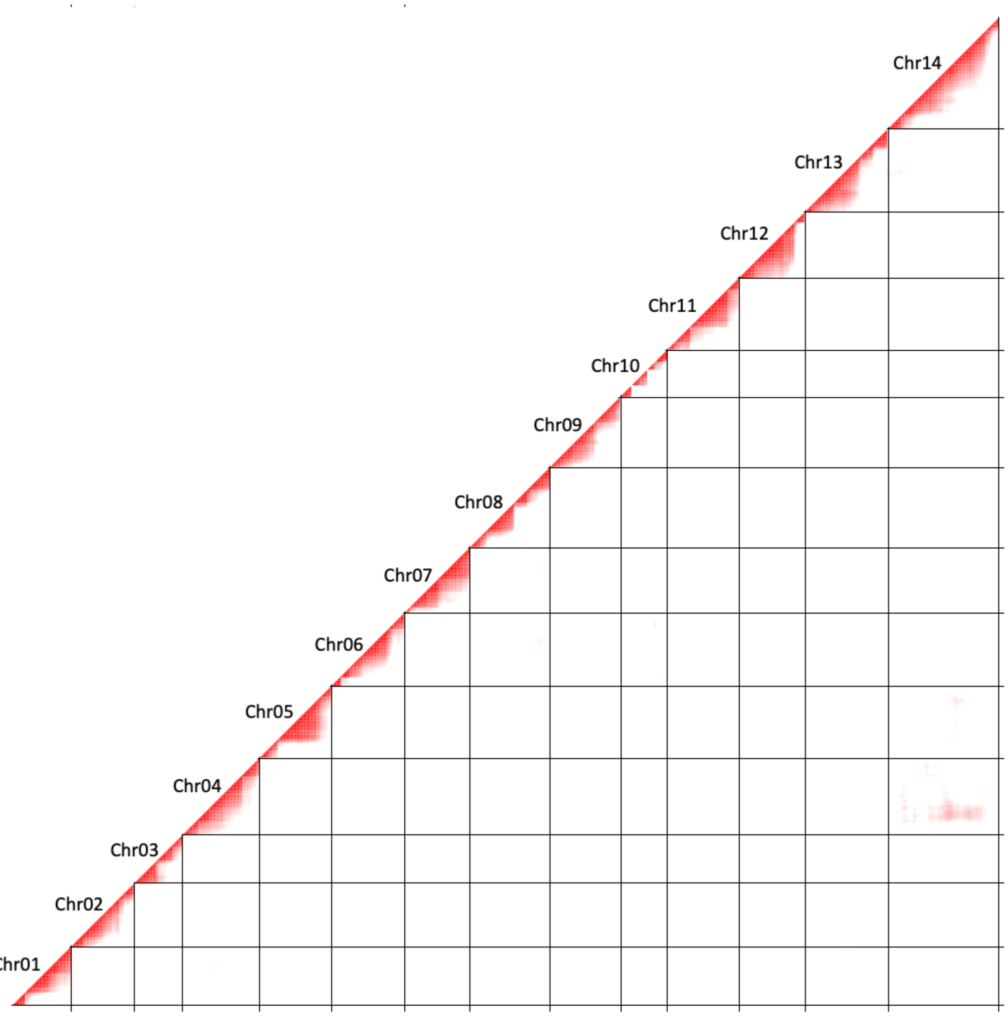


**Figure S12. CAC415 RILs linkage disequilibrium heatmap.** LD heatmap for each pairwise marker comparison across all 14 chromosomes, showing r values greater than the 95% quantile (>0.39) in the third RIL population. Values range from 0 (white) to 1 (red).

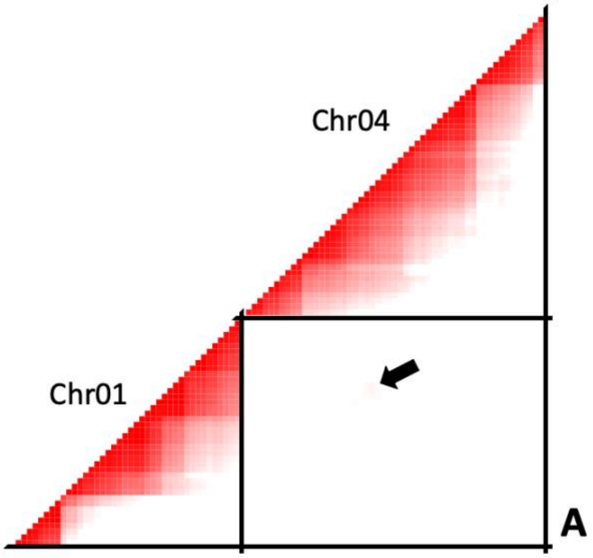
Chr04
Chr01
A

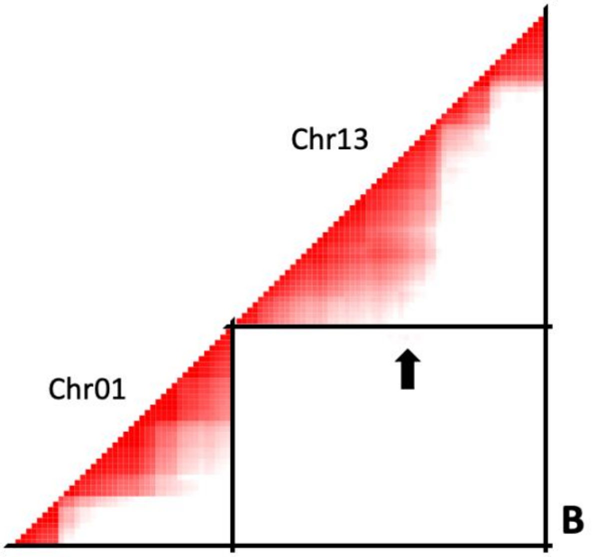
Chr13
Chr01
B

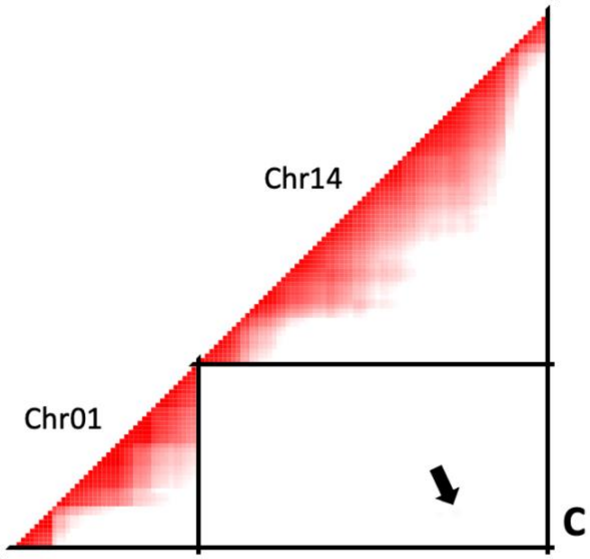
Chr14
Chr01
C

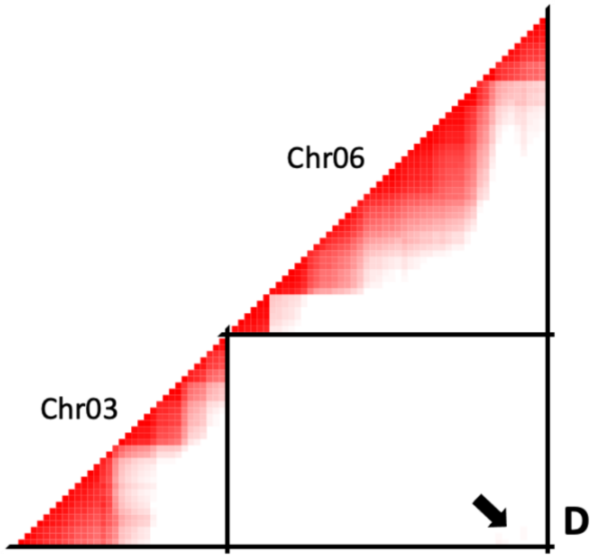
Chr06
Chr03
D

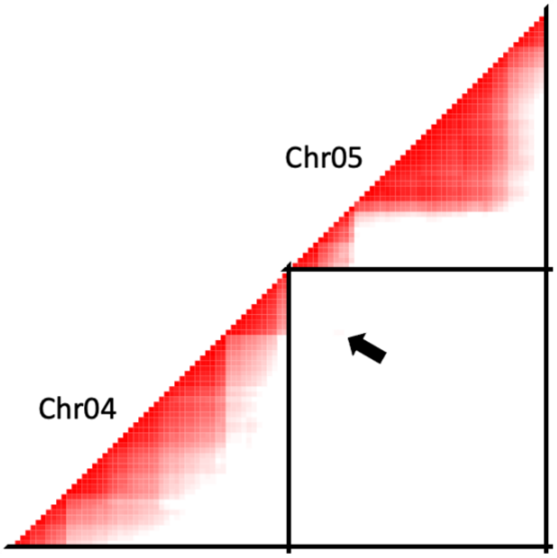


E

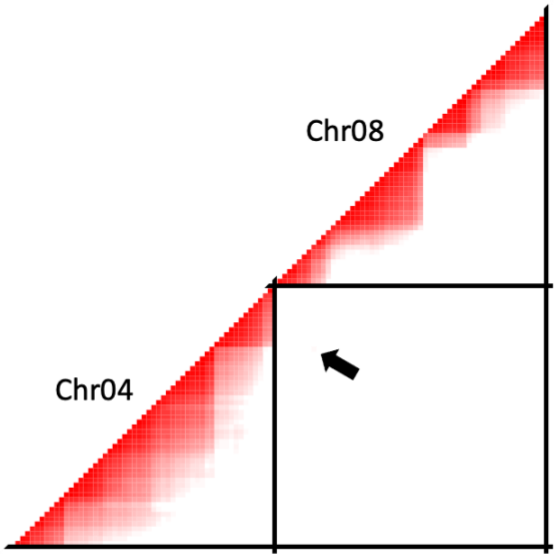


F

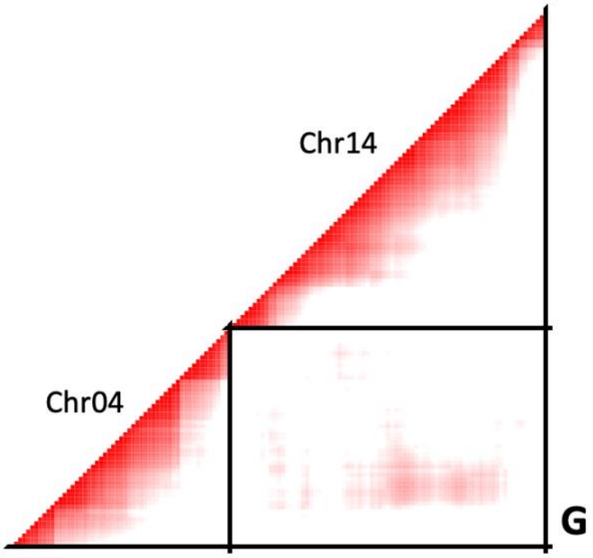


238

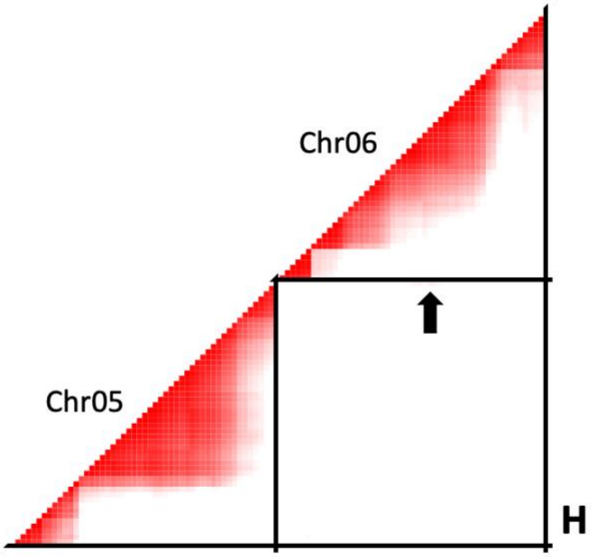


239

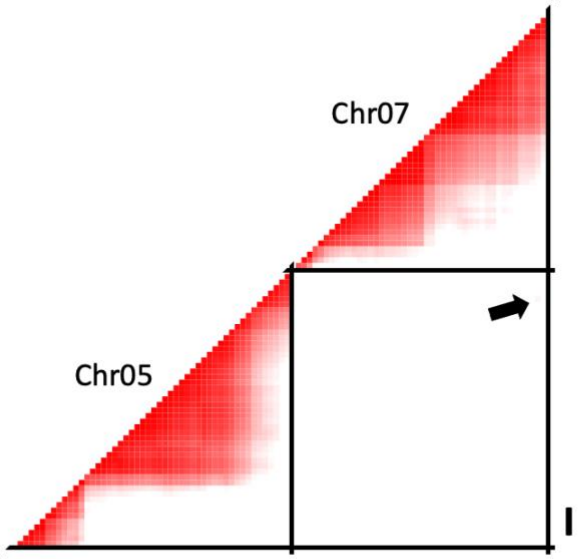


240

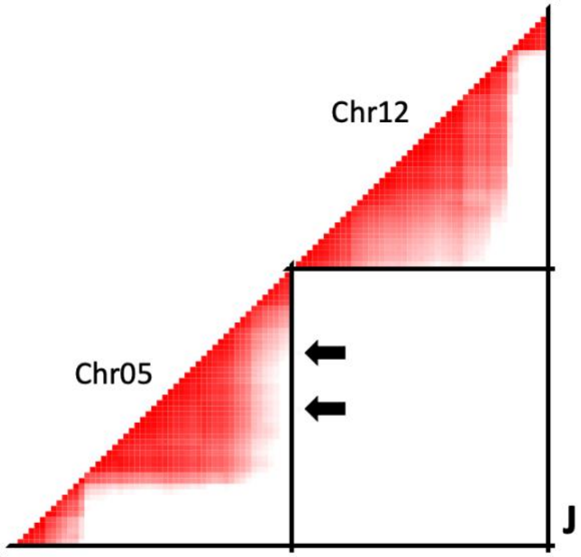


241

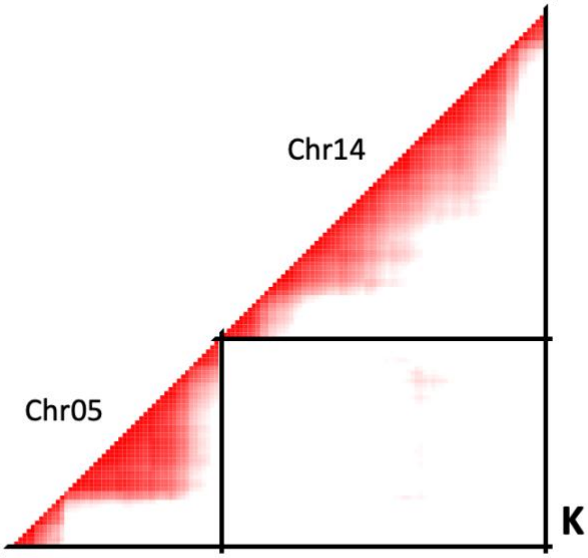
Chr14
Chr05
K

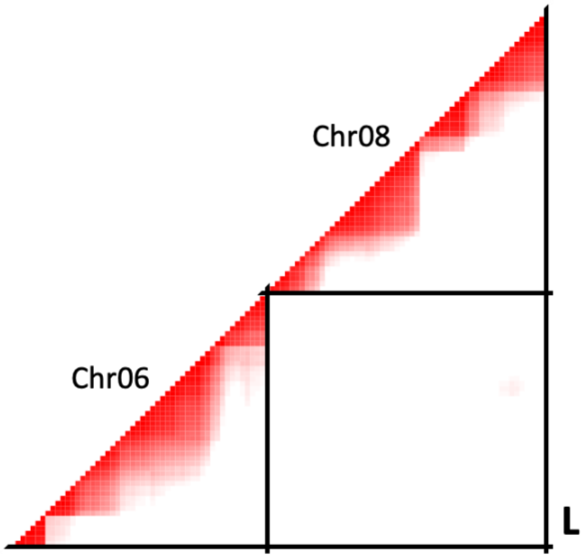
Chr08
Chr06
L

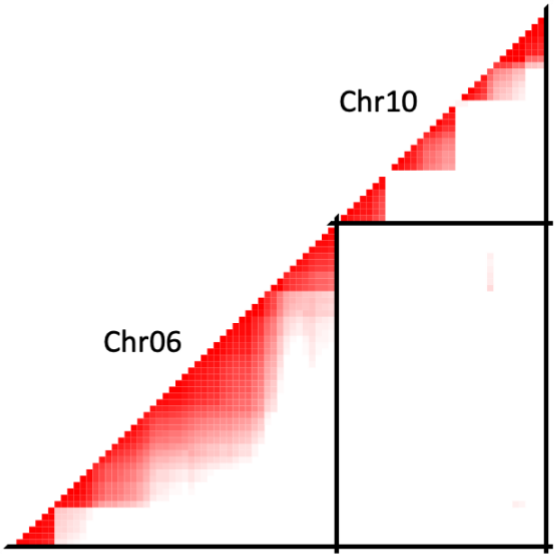


M

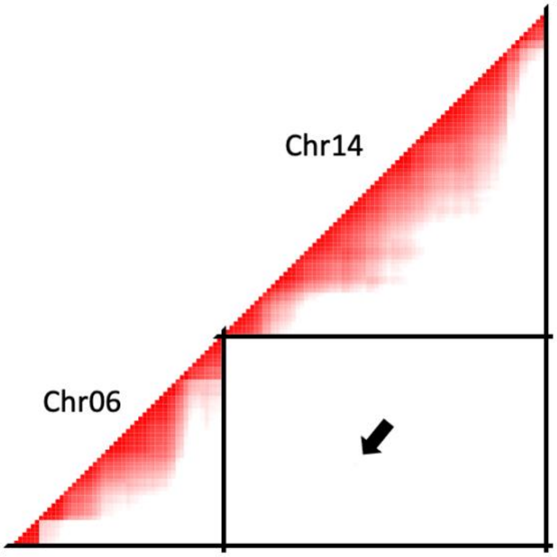


N

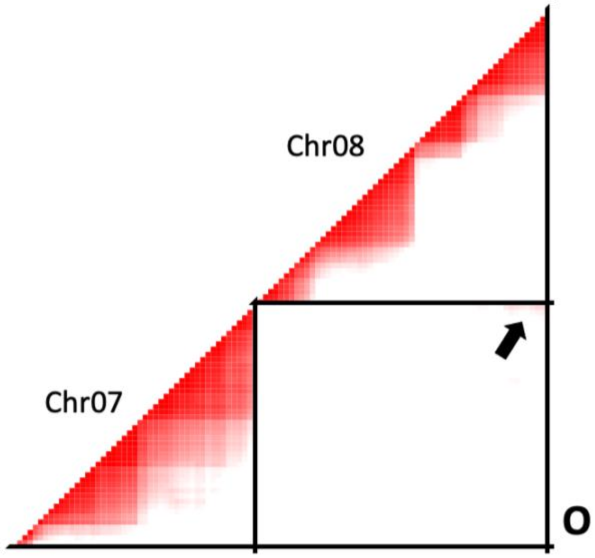


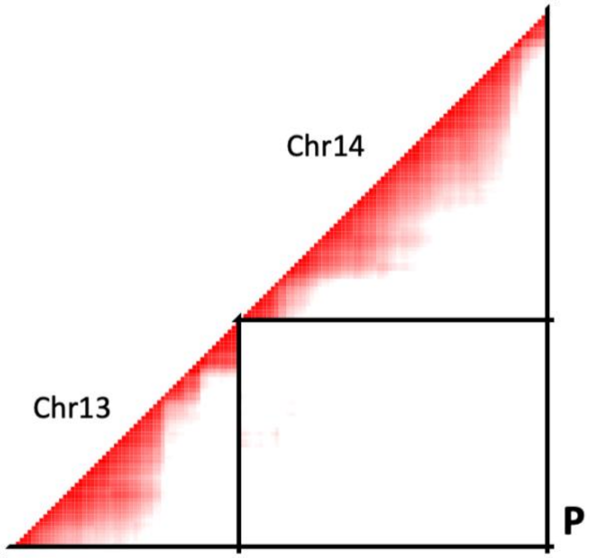


**Figure S13. CAC415 RILs linkage disequilibrium heatmap.**

LD heatmaps for each pairwise marker comparison, showing r values greater than the 95% quantile (>0.39) in chromosome comparisons with at least 1 marker above the cutoff in the third RIL population. Values range from 0 (white) to 1 (red). Arrows indicates significant marker comparisons, when distorted regions are small. (a) Chr01 and Chr04 (b) Chr01 and Chr13 (c) Chr01 and Chr14 (d) Chr03 and Chr06 (e) Chr04 and Chr05 (f) Chr04 and Chr08 (g) Chr04 and Chr14 (h) Chr05 and Chr06 (i) Chr05 and Chr07 (j) Chr05 and Chr12 (k) Chr05 and Chr14 (l) Chr06 and Chr08 (m) Chr06 and Chr10 (n) Chr06 and Chr14 (o) Chr07 and Chr08 (p) Chr13 and Chr14.

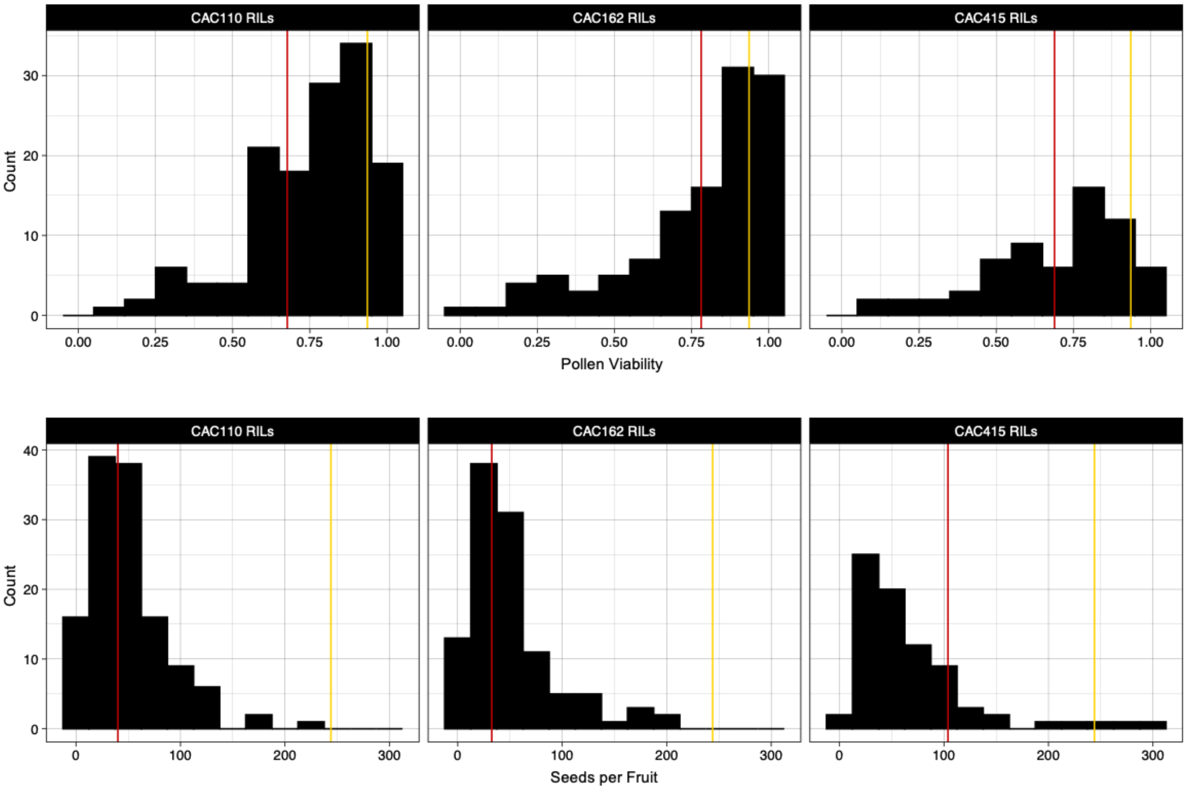


**Figure S14. Male and female fertility distributions in the three RIL populations. (a)** Male fertility (proportion viable pollen grains) in the CAC110 (N=138), CAC162 (N=116), and CAC415 RILs (N=65). Yellow lines indicate CAC9 average value (0.94), red lines indicate *M. guttatus* parent average values (0.68, 0.78, and 0.69 respectively). **(b)** Female fertility (seeds per fruit) in the CAC110 (N=127), CAC162 (N=109), and CAC415 RILs (N=78). Yellow lines indicate CAC9 average value (244.00), red lines indicate *M. guttatus* parent average values (40.08, 32.53, and 103.85 respectively).

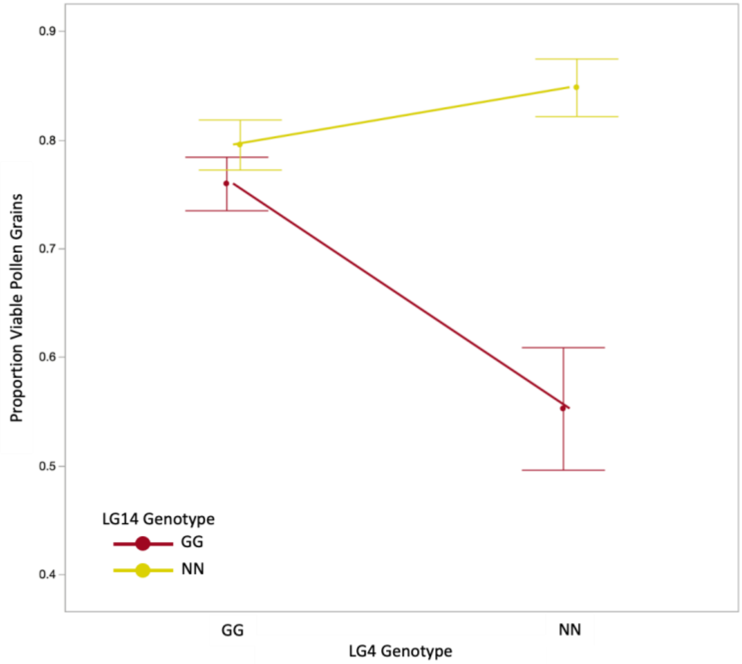
0.9
0.8
0.7
0.6
0.5
0.4
Proportion Viable Pollen Grains
LG14 Genotype
GG
NN
GG
NN
LG4 Genotype

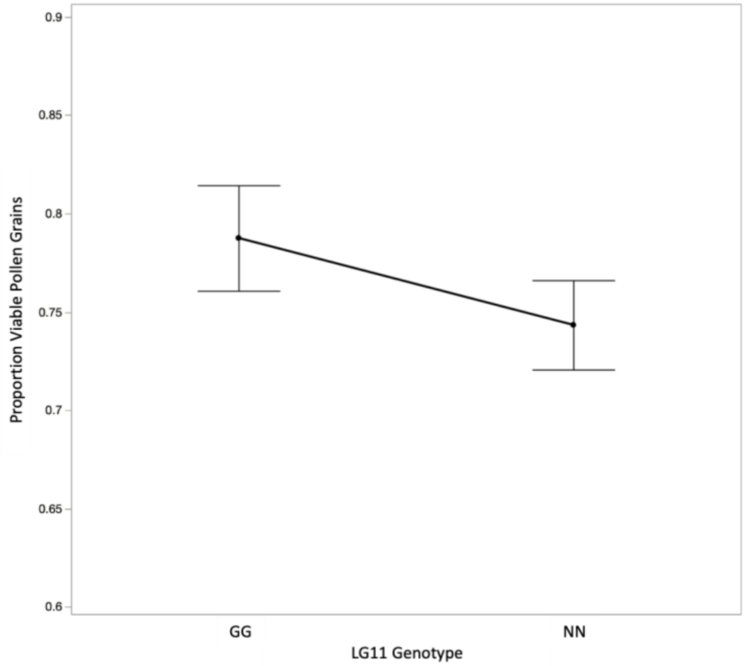
0.9
0.85
0.8
0.75
0.7
0.65
0.6
Proportion Viable Pollen Grains
GG
NN
LG11 Genotype

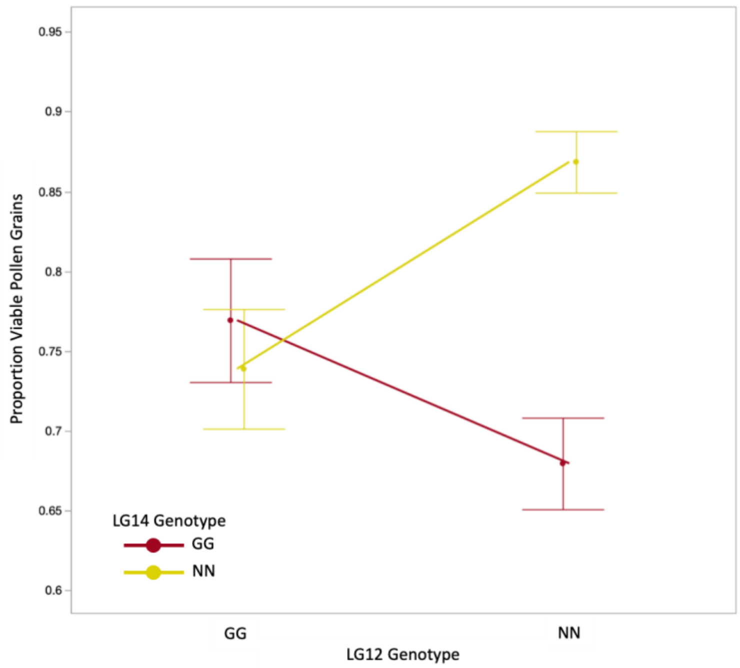


**Figure S15. CAC110 RILs male fertility effect plots.** Average proportion viable pollen grains of each genotypic class for each significant additive and epistatic QTL.

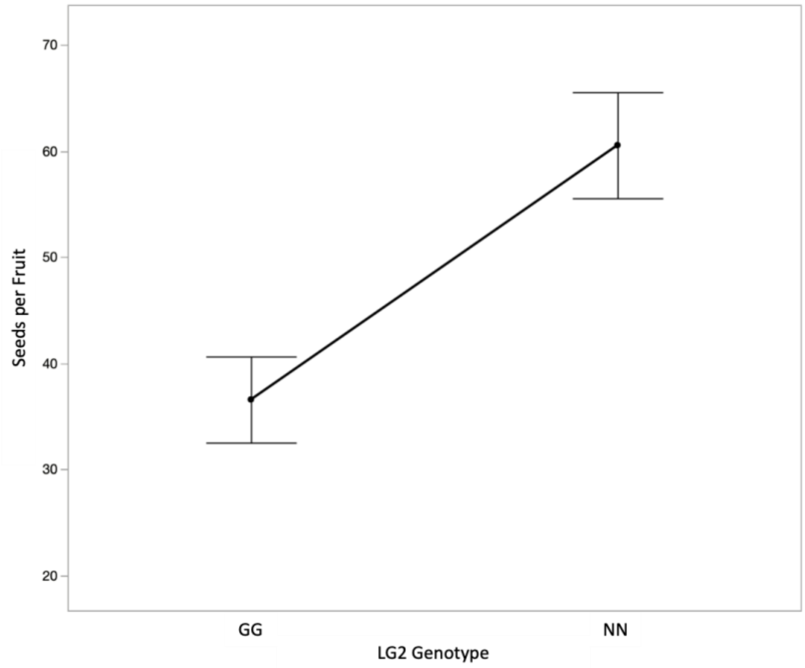
Seeds per Fruit
70
60
50
40
30
20
GG
NN
LG2 Genotype

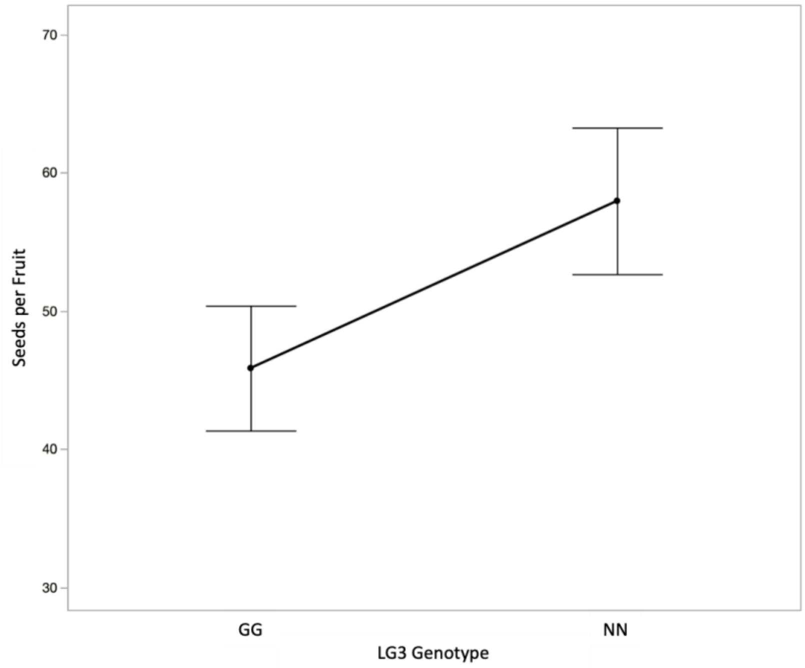
Seeds per Fruit
70
60
50
40
30
GG
NN
LG3 Genotype

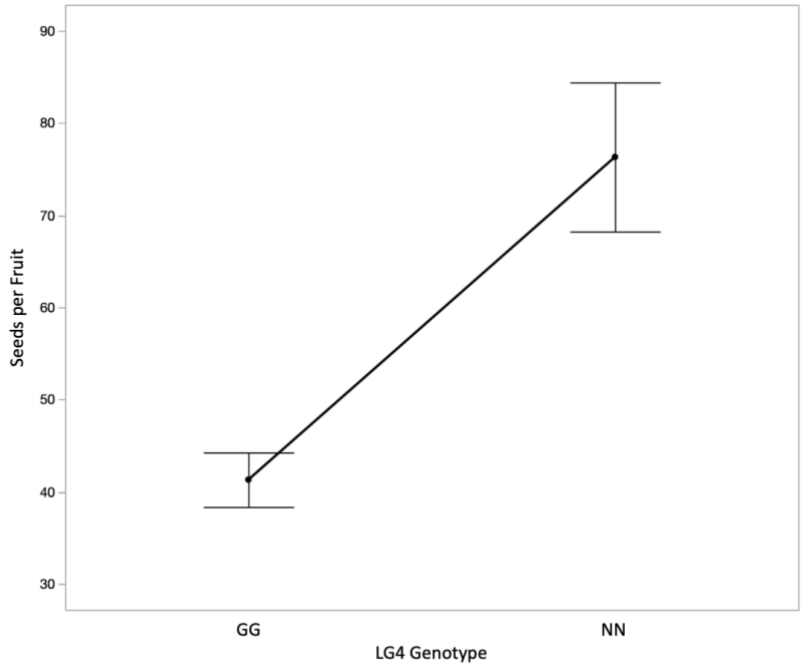


**Figure S16. CAC110 RILs female fertility effect plots.** Average seeds per fruit of each genotypic class for each significant additive QTL.

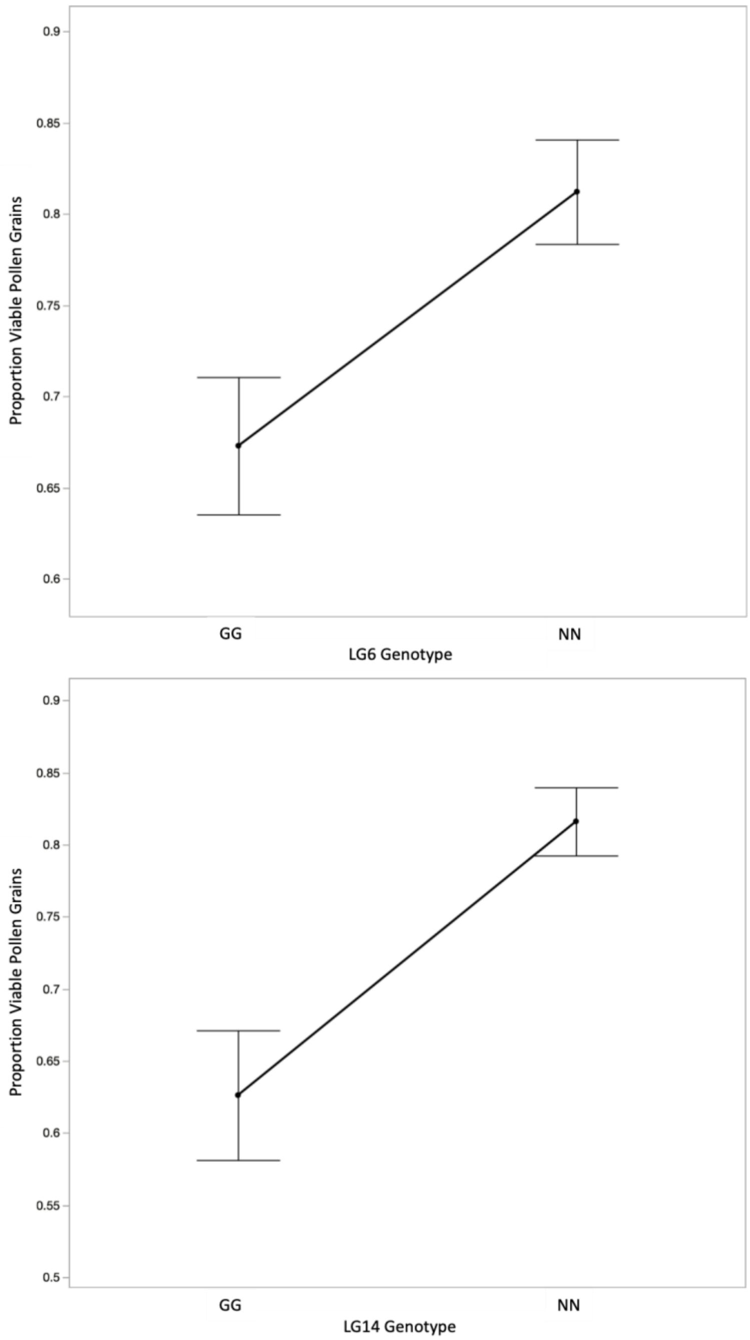


**Figure S17. CAC162 RILs male fertility effect plots.** Average proportion viable pollen grains of each genotypic class for each significant additive QTL.

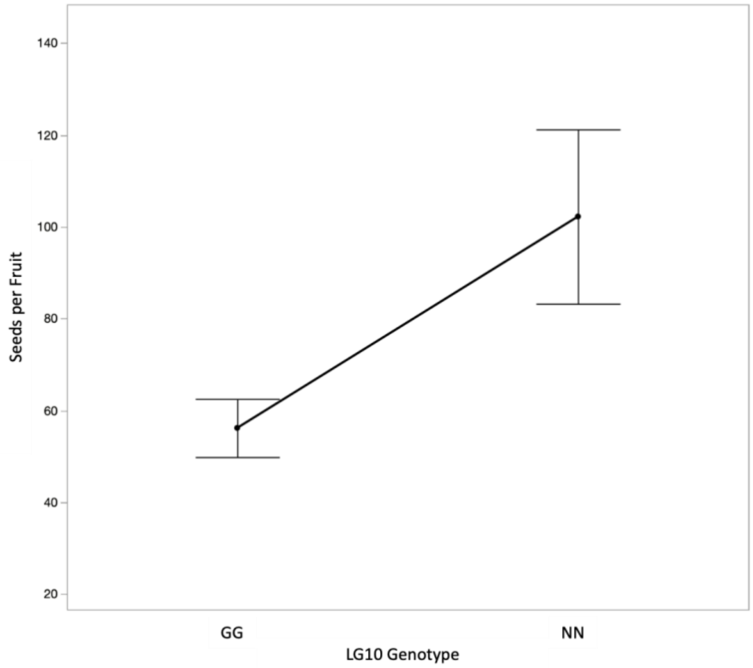


**Figure S18. CAC415 RILs female fertility effect plots.** Average seeds per fruit of each genotypic class for each significant additive QTL.

**Table S1. Geographic locations of *Mimulus* populations used.**
Information about the biology, geography, and sequencing of each sample used in this manuscript.

| Line abbreviation | Species/Patry | Population | Latitude (N) | Longitude (W) | NCBI SRA Accession No. |
|---|---|---|---|---|---|
| CAC110 | sympatric *M. guttatus* | Catherine Creek, Washington side of the Columbia River Gorge off of Hwy. 14 | 45°42′42′′ | 121°21′55′′ | TBA |
| CAC112 | | | | | TBA |
| CAC134 | | | | | TBA |
| CAC141 | | | | | TBA |
| CAC162 | | | | | TBA |
| CAC262 | | | | | TBA |
| CAC415 | | | | | TBA |
| CAC6 | | | | | SRX525044 |
| CAC9 | sympatric *M. nasutus* | Catherine Creek, Washington side of the Columbia River Gorge off of Hwy. 14 | | | SRX525048 |
| IM62 | allopatric *M. guttatus* | Iron Mountain, Hwy. 20, Linn County, Oregon | 44° 24' 03" | 122° 08' 16" | SRX115898 |
| DPRG102 | sympatric *M. guttatus* | Don Pedro Reservoir, Stanislaus National Forest, Junction of Hwy. 120 and Jacksonville Road, Tuolumne County, California | 37° 49' 44" | 120° 20' 38" | TBA |
| SF5 | allopatric *M. nasutus* | Sherar's Falls, Tygh Valley, Wasco County, Oregon | 45°15′52′′ | 121°01′21′′ | SRX116529 |
| KOOT | allopatric *M. nasutus* | West of Kootenai National Forest, off MT 200, north of Cabinet Gorge Reservoir | 48° 06' 14" | 115° 58' 59" | SRX525049 |
| NHN26 | sympatric *M. nasutus* | Notch Hill, north of Nanoose Harbour, Vancouver Island, BC | 49° 16' 23' | 124° 09' 36" | SRX525051 |

**Table S2. Linkage map construction details.** Information regarding the number of markers, and program options in LepMAP3 used to construct each of the three linkage maps featured in this manuscript. Each chromosome was either run through SeparateChromosomes2 on its own (independently) or as a group with other chromosomes in the same family (together). Physical orders with better higher likelihoods when run through OrderMarkers2 are shown in bold. The three linkage maps contain as total of 4,967 unique markers among them, 3,141 (~63%) of which are present in all three maps. 513 markers (~10%) are unique to a single map, leaving 1,330 markers (~27%) shared between two of the three maps.

| | | | SeparateChromosomes2 | | | OrderMarkers2 | | | |
|---|---|---|---|---|---|---|---|---|---|
| | LG | Number of markers | Run conditions | LodLimit | Theta | Highest Likelihood 10 iterations | Physical Order Likelihood | Retained positions | Unique positions |
| CAC110 RILs | 1 | 156 | independently | 0.1 | 0.15 | -7,008.42 | - | 156 | 60 |
| | 2 | 314 | independently | 0.1 | 0.15 | -6,946.23 | - | 314 | 49 |
| | 3 | 292 | together | 20 | 0.3 | -6,326.90 | - | 271 | 40 |
| | 4 | 279 | independently | 0.1 | 0.15 | -6,730.64 | - | 279 | 47 |
| | 5 | 305 | together | 20 | 0.3 | -10,574.70 | - | 277 | 42 |
| | 6 | 335 | together | 20 | 0.3 | -7,543.75 | - | 332 | 57 |
| | 7 | 279 | together | 20 | 0.3 | -4,606.16 | - | 252 | 46 |
| | 8 | 352 | independently | 0.1 | 0.15 | -15,013.73 | -15,514.09 | 349 | 88 |
| | 9 | 255 | together | 20 | 0.3 | -4,355.72 | - | 255 | 41 |
| | 10 | 322 | together | 20 | 0.3 | -8,102.44 | -16,185.25 | 319 | 55 |
| | 11 | 115 | independently | 0.1 | 0.15 | -9,735.51 | **-8,846.31** | 115 | 67 |
| | 12 | 317 | independently | 0.1 | 0.15 | -11,752.58 | - | 304 | 68 |
| | 13 | 166 | independently | 0.1 | 0.15 | -6,879.29 | -7,732.36 | 138 | 50 |
| | 14 | 467 | together | 20 | 0.3 | -14,236.16 | - | 467 | 84 |
| | Global | 3954 | - | - | - | - | - | 3828 | 794 |
| CAC162 RILs | 1 | 218 | together | 15 | 0.25 | -3,311.21 | - | 213 | 36 |
| | 2 | 340 | together | 15 | 0.25 | -5,011.07 | - | 340 | 47 |
| | 3 | 309 | together | 15 | 0.25 | -8,771.58 | - | 291 | 54 |
| | 4 | 359 | independently | 0.1 | 0.15 | -6,319.86 | - | 359 | 36 |
| | 5 | 331 | together | 15 | 0.25 | -7,611.04 | - | 307 | 36 |
| | 6 | 315 | independently | 0.1 | 0.15 | -7,782.91 | - | 315 | 48 |
| | 7 | 243 | together | 15 | 0.25 | -6,165.75 | - | 222 | 36 |
| | 8 | 392 | independently | 0.1 | 0.15 | -11,342.19 | **-11,224.64** | 392 | 54 |
| | 9 | 265 | together | 15 | 0.25 | -6,308.35 | - | 265 | 45 |
| | 10 | 327 | independently | 0.1 | 0.15 | -7,708.27 | - | 303 | 25 |
| | 11 | 260 | independently | 0.1 | 0.15 | -8,465.66 | -9,211.12 | 260 | 35 |
| | 12 | 338 | together | 15 | 0.25 | -9,874.55 | - | 316 | 47 |
| | 13 | 340 | together | 15 | 0.25 | -4,384.34 | - | 340 | 38 |
| | 14 | 466 | together | 15 | 0.25 | -12,223.42 | - | 466 | 69 |
| | Global | 4503 | - | - | - | - | - | 4389 | 606 |
| CAC415 RILs | 1 | 186 | together | 15 | 0.25 | -5,020.23 | - | 174 | 41 |
| | 2 | 314 | together | 15 | 0.25 | -6,706.58 | - | 314 | 44 |
| | 3 | 302 | independently | 0.1 | 0.15 | -7,280.74 | - | 289 | 34 |
| | 4 | 364 | independently | 0.1 | 0.15 | -8,592.37 | - | 363 | 54 |
| | 5 | 327 | independently | 0.1 | 0.15 | -7,690.35 | - | 325 | 50 |
| | 6 | 323 | independently | 0.1 | 0.15 | -7,628.57 | - | 322 | 51 |
| | 7 | 245 | together | 15 | 0.25 | -7,388.27 | - | 237 | 46 |
| | 8 | 438 | together | 15 | 0.25 | -12,658.23 | - | 436 | 56 |
| | 9 | 268 | independently | 0.1 | 0.15 | 6,441.53 | - | 266 | 49 |
| | 10 | 258 | together | 15 | 0.25 | -5,819.33 | - | 252 | 33 |
| | 11 | 289 | together | 15 | 0.25 | -10,483.63 | -11,605.05 | 255 | 50 |
| | 12 | 333 | independently | 0.1 | 0.15 | -7,276.95 | - | 322 | 46 |
| | 13 | 343 | independently | 0.1 | 0.15 | -8,136.78 | - | 343 | 58 |
| | 14 | 489 | independently | 0.1 | 0.15 | -15,535.39 | - | 481 | 78 |
| | Global | 4479 | - | - | - | - | - | 4379 | 690 |

**Table S3. Positions of significant transmission ratio distortion.** Genetic (cM) and physical (Mb) position on each chromosome in the three RIL populations that show significant distortion (Direction) via Chi-Square test at α = 0.01 and Bonferonni-corrected α. Shading indicates putative contiguous regions of TRD.

| Chromosome | cM | Mb | Direction | RIL Population | α Threshold |
|---|---|---|---|---|---|
| 01 | 0.000 - 15.128 | 0.05 - 3.90 | *M. guttatus* | CAC110 | 0.01 |
| 01 | 10.930 - 12.679 | 3.30 - 3.45 | *M. guttatus* | CAC110 | 0.000167 |
| 01 | 32.397 | 4.10 - 4.25 | *M. guttatus* | CAC110 | 0.01 |
| 01 | 122.469 - 134.368 | 11.05 - 12.85 | *M. guttatus* | CAC110 | 0.01 |
| 01 | 124.218 - 134.368 | 11.10 - 12.85 | *M. guttatus* | CAC110 | 0.000167 |
| 01 | NA | NA | NA | CAC162 | 0.01 |
| 01 | NA | NA | NA | CAC162 | 0.000278 |
| 01 | 1.220 - 28.693 | 0.50 - 3.60 | *M. guttatus* | CAC415 | 0.01 |
| 01 | 12.204 | 3.40 - 3.45 | *M. guttatus* | CAC415 | 0.000244 |
| 01 | 72.640 - 74.470 | 11.20 - 11.70 | *M. nasutus* | CAC415 | 0.01 |
| 02 | 64.202 - 64.551 | 18.40 - 18.80 | *M. nasutus* | CAC110 | 0.01 |
| 02 | 74.023 - 86.970 | 18.95 - 20.55 | *M. nasutus* | CAC110 | 0.01 |
| 02 | 81.370 - 83.468 | 19.90 - 20.10 | *M. nasutus* | CAC110 | 0.000204 |
| 02 | 34.741 - 36.783 | 3.40 - 5.25 | *M. nasutus* | CAC162 | 0.01 |
| 02 | 38.823 - 39.844 | 5.50 - 12.05 | *M. nasutus* | CAC162 | 0.01 |
| 02 | 40.864 - 43.926 | 12.15 - 15.70 | *M. nasutus* | CAC162 | 0.01 |
| 02 | NA | NA | NA | CAC162 | 0.000213 |
| 02 | 56.154 | 18.50 - 18.75 | *M. guttatus* | CAC415 | 0.01 |
| 02 | NA | NA | NA | CAC415 | 0.000227 |
| 03 | NA | NA | NA | CAC110 | 0.01 |
| 03 | NA | NA | NA | CAC110 | 0.000250 |
| 03 | 22.452 - 31.637 | 19.65 - 20.85 | *M. nasutus* | CAC162 | 0.01 |
| 03 | 44.449 - 64.360 | 21.15 - 23.80 | *M. nasutus* | CAC162 | 0.01 |
| 03 | 54.660 - 56.702 | 22.75 - 23.15 | *M. nasutus* | CAC162 | 0.000185 |
| 03 | 0.000 - 17.078 | 0.25 - 19.95 | *M. guttatus* | CAC415 | 0.01 |
| 03 | 0.000 - 10.978 | 1.20 - 19.95 | *M. guttatus* | CAC415 | 0.000294 |
| 04 | 0.000 - 35.719 | 0.00 - 19.65 | *M. guttatus* | CAC110 | 0.01 |
| 04 | 0.000 - 35.719 | 0.00 - 19.65 | *M. guttatus* | CAC110 | 0.000213 |
| 04 | 39.221 - 44.119 | 19.80 - 20.15 | *M. guttatus* | CAC110 | 0.01 |
| 04 | 0.000 - 16.873 | 0.05 - 3.70 | *M. nasutus* | CAC162 | 0.01 |
| 04 | 0.000 | 0.05 - 0.45 | *M. nasutus* | CAC162 | 0.000278 |
| 04 | 3.062 - 13.298 | 1.20 - 3.20 | *M. nasutus* | CAC162 | 0.000278 |
| 04 | 16.873 | 3.45 - 3.70 | *M. nasutus* | CAC162 | 0.000278 |
| 04 | 42.725 - 54.929 | 6.45 - 18.95 | *M. guttatus* | CAC415 | 0.01 |
| 04 | NA | NA | NA | CAC415 | 0.000185 |

| | | | | | |
|---|---|---|---|---|---|
| 05 | 0.000 - 82.002 | 0.00 - 20.90 | *M. guttatus* | CAC110 | 0.01 |
| 05 | 0.000 - 82.002 | 0.00 - 20.90 | *M. guttatus* | CAC110 | 0.000238 |
| 05 | NA | NA | NA | CAC162 | 0.01 |
| 05 | NA | NA | NA | CAC162 | 0.000278 |
| 05 | 2.441 - 9.765 | 0.50 - 1.75 | *M. guttatus* | CAC415 | 0.01 |
| 05 | 78.303 - 89.303 | 19.45 - 21.25 | *M. nasutus* | CAC415 | 0.01 |
| 05 | NA | NA | NA | CAC415 | 0.000200 |
| 06 | NA | NA | NA | CAC110 | 0.01 |
| 06 | NA | NA | NA | CAC110 | 0.000175 |
| 06 | 0.000 - 27.785 | 0.00 - 4.05 | *M. guttatus* | CAC162 | 0.01 |
| 06 | 8.689 - 27.785 | 1.80 - 4.05 | *M. guttatus* | CAC162 | 0.000208 |
| 06 | 67.161 | 8.50 - 9.25 | *M. guttatus* | CAC162 | 0.01 |
| 06 | 69.203 | 13.90 - 14.30 | *M. guttatus* | CAC162 | 0.01 |
| 06 | 69.713 | 14.35 - 14.40 | *M. guttatus* | CAC162 | 0.01 |
| 06 | 69.713 | 14.60 - 14.70 | *M. guttatus* | CAC162 | 0.01 |
| 06 | 57.782 | 6.9 - 7.05 | *M. nasutus* | CAC415 | 0.01 |
| 06 | 60.221 | 7.60 - 9.20 | *M. nasutus* | CAC415 | 0.01 |
| 06 | 60.221 | 9.5 - 9.65 | *M. nasutus* | CAC415 | 0.01 |
| 06 | 60.221 | 10.25 - 15.40 | *M. nasutus* | CAC415 | 0.01 |
| 06 | NA | NA | NA | CAC415 | 0.000196 |
| 07 | 45.655 - 72.643 | 3.05 - 21.80 | *M. guttatus* | CAC110 | 0.01 |
| 07 | 53.409 - 64.250 | 3.10 - 8.75 | *M. guttatus* | CAC110 | 0.000217 |
| 07 | 65.299 - 72.643 | 15.00 - 21.80 | *M. guttatus* | CAC110 | 0.000217 |
| 07 | NA | NA | NA | CAC162 | 0.01 |
| 07 | NA | NA | NA | CAC162 | 0.000278 |
| 07 | 56.933 | 6.90 - 6.95 | *M. nasutus* | CAC415 | 0.01 |
| 07 | 55.713 - 73.440 | 8.30 - 14.40 | *M. nasutus* | CAC415 | 0.01 |
| 07 | 64.900 | 12.85 - 13.05 | *M. nasutus* | CAC415 | 0.000217 |
| 07 | 68.561 - 72.22 | 15.20 - 16.25 | *M. nasutus* | CAC415 | 0.01 |
| 08 | 56.691 - 67.882 | 6.60 - 9.00 | *M. nasutus* | CAC110 | 0.01 |
| 08 | 56.691 - 62.287 | 6.60 - 7.20 | *M. nasutus* | CAC110 | 0.000114 |
| 08 | 63.686 | 7.90 - 7.95 | *M. nasutus* | CAC110 | 0.000114 |
| 08 | 63.686 | 8.95 - 9.00 | *M. nasutus* | CAC110 | 0.000114 |
| 08 | 53.543 - 53.893 | 14.90 - 15.15 | *M. nasutus* | CAC110 | 0.01 |
| 08 | 50.745 - 66.133 | 15.85 - 17.05 | *M. nasutus* | CAC110 | 0.01 |
| 08 | 65.084 - 66.133 | 15.85 - 16.05 | *M. nasutus* | CAC110 | 0.000114 |
| 08 | 72.088 - 109.184 | 18.35 - 27.70 | *M. nasutus* | CAC110 | 0.01 |
| 08 | 82.934 - 87.492 | 21.25 - 21.65 | *M. nasutus* | CAC110 | 0.000114 |
| 08 | 93.096 - 100.091 | 21.95 - 23.80 | *M. nasutus* | CAC110 | 0.000114 |
| 08 | 102.889 | 24.10 - 24.25 | *M. nasutus* | CAC110 | 0.000114 |
| 08 | 104.288 | 25.35 - 26.40 | *M. nasutus* | CAC110 | 0.000114 |
| 08 | 39.841 - 55.513 | 4.25 - 7.55 | *M. nasutus* | CAC162 | 0.01 |
| 08 | 71.385 - 117.963 | 21.50 - 27.35 | *M. nasutus* | CAC162 | 0.01 |
| 08 | 80.576 - 83.129 | 22.10 - 22.20 | *M. nasutus* | CAC162 | 0.000185 |
| 08 | 95.938 - 117.963 | 22.70 - 27.35 | *M. nasutus* | CAC162 | 0.000185 |
| 08 | 9.765 | 1.70 - 1.80 | *M. nasutus* | CAC415 | 0.01 |
| 08 | NA | NA | NA | CAC415 | 0.000179 |

| | | | | | |
|---|---|---|---|---|---|
| 09 | NA | NA | NA | CAC110 | 0.01 |
| 09 | NA | NA | NA | CAC110 | 0.000244 |
| 09 | NA | NA | NA | CAC162 | 0.01 |
| 09 | NA | NA | NA | CAC162 | 0.000222 |
| 09 | NA | NA | NA | CAC415 | 0.01 |
| 09 | NA | NA | NA | CAC415 | 0.000204 |
| 10 | 19.249 - 20.648 | 5.10 - 5.30 | *M. nasutus* | CAC110 | 0.01 |
| 10 | 88.257 - 89.656 | 14.90 - 15.80 | *M. guttatus* | CAC110 | 0.01 |
| 10 | NA | NA | NA | CAC110 | 0.000182 |
| 10 | NA | NA | NA | CAC162 | 0.01 |
| 10 | NA | NA | NA | CAC162 | 0.000400 |
| 10 | 0.000 - 1.220 | 0.00 - 0.77 | *M. nasutus* | CAC415 | 0.01 |
| 10 | 53.554 - 104.91 | 1.80 - 22.00 | *M. guttatus* | CAC415 | 0.01 |
| 10 | 53.554 - 95.746 | 1.80 - 20.25 | *M. guttatus* | CAC415 | 0.000303 |
| 10 | 98.797 | 20.30 - 20.40 | *M. guttatus* | CAC415 | 0.000303 |
| 11 | 54.170 - 56.618 | 5.05 - 5.75 | *M. guttatus* | CAC110 | 0.01 |
| 11 | 82.175 - 85.323 | 9.60 - 9.90 | *M. nasutus* | CAC110 | 0.01 |
| 11 | 94.429 - 132.951 | 10.35 - 26.70 | *M. nasutus* | CAC110 | 0.01 |
| 11 | 102.823 - 103.523 | 23.50 - 23.65 | *M. nasutus* | CAC110 | 0.000149 |
| 11 | 119.282 - 124.889 | 25.30 - 26.60 | *M. nasutus* | CAC110 | 0.000149 |
| 11 | 0.000 - 2.551 | 0.05 - 1.10 | *M. nasutus* | CAC162 | 0.01 |
| 11 | 1.021 - 2.551 | 0.75 - 1.10 | *M. nasutus* | CAC162 | 0.000286 |
| 11 | NA | NA | NA | CAC415 | 0.01 |
| 11 | NA | NA | NA | CAC415 | 0.000200 |
| 12 | 15.399 - 87.186 | 1.05 - 24.50 | *M. nasutus* | CAC110 | 0.01 |
| 12 | 21.705 - 25.202 | 1.70 - 5.20 | *M. nasutus* | CAC110 | 0.000147 |
| 12 | 22.404 - 29.398 | 5.30 - 17.60 | *M. nasutus* | CAC110 | 0.000147 |
| 12 | 38.49 - 60.193 | 18.60 - 21.70 | *M. nasutus* | CAC110 | 0.000147 |
| 12 | 65.090 | 22.55 - 22.70 | *M. nasutus* | CAC110 | 0.000147 |
| 12 | 68.239 - 80.538 | 22.75 - 23.60 | *M. nasutus* | CAC110 | 0.000147 |
| 12 | 8.168 - 9.188 | 1.30 - 1.90 | *M. nasutus* | CAC162 | 0.01 |
| 12 | 15.823 - 16.334 | 2.00 - 2.55 | *M. nasutus* | CAC162 | 0.01 |
| 12 | 18.375 - 18.885 | 2.70 - 3.10 | *M. nasutus* | CAC162 | 0.01 |
| 12 | 17.864 - 30.110 | 5.05 - 7.95 | *M. nasutus* | CAC162 | 0.01 |
| 12 | 30.110 | 6.20 - 6.25 | *M. nasutus* | CAC162 | 0.000213 |
| 12 | 30.621 | 18.55 - 18.60 | *M. nasutus* | CAC162 | 0.01 |
| 12 | 30.621 | 18.55 - 18.60 | *M. nasutus* | CAC162 | 0.000213 |
| 12 | 32.152 - 67.944 | 18.65 - 24.20 | *M. nasutus* | CAC162 | 0.01 |
| 12 | 32.152 - 32.152 | 18.65 - 19.20 | *M. nasutus* | CAC162 | 0.000213 |
| 12 | 37.259 - 45.425 | 19.80 - 23.40 | *M. nasutus* | CAC162 | 0.000213 |
| 12 | 69.986 - 75.611 | 24.30 - 24.90 | *M. nasutus* | CAC162 | 0.01 |
| 12 | NA | NA | NA | CAC415 | 0.01 |
| 12 | NA | NA | NA | CAC415 | 0.000217 |

| | | | | | |
|---|---|---|---|---|---|
| 13 | 0.000 - 53.932 | 2.10 - 25.00 | *M. guttatus* | CAC110 | 0.01 |
| 13 | 0.000 - 45.185 | 2.10 - 24.00 | *M. guttatus* | CAC110 | 0.000200 |
| 13 | 88.949 - 117.730 | 26.50 - 30.80 | *M. nasutus* | CAC110 | 0.01 |
| 13 | 88.949 - 117.730 | 26.50 - 30.80 | *M. nasutus* | CAC110 | 0.000200 |
| 13 | 38.986 - 78.866 | 25.55 - 30.80 | *M. nasutus* | CAC162 | 0.01 |
| 13 | 54.367 - 78.866 | 26.30 - 30.80 | *M. nasutus* | CAC162 | 0.000263 |
| 13 | 61.649 | 25.50 - 25.55 | *M. guttatus* | CAC415 | 0.01 |
| 13 | NA | NA | NA | CAC415 | 0.000172 |
| 14 | 0.000 - 22.761 | 0.00 - 2.35 | *M. guttatus* | CAC110 | 0.01 |
| 14 | 0.000 - 9.099 | 0.00 - 1.90 | *M. guttatus* | CAC110 | 0.000119 |
| 14 | 0.000 - 3.062 | 0.00 - 0.60 | *M. guttatus* | CAC162 | 0.01 |
| 14 | NA | NA | NA | CAC162 | 0.000145 |
| 14 | 0.000 - 13.422 | 0.00 - 2.65 | *M. guttatus* | CAC415 | 0.01 |
| 14 | NA | NA | NA | CAC415 | 0.000128 |

**Table S4. Average Interchromosomal linkage disequilibrium (r) across all pairwise marker comparisons for each chromosome pair in each RIL population.** N is the total number of pairwise marker comparisons. r95 indicates average ILD of those marker comparisons which are greater than the 95% quantile. N95 is the number of those pairwise comparisons that are greater than the 95% quantile. NA indicates there were no pairwise marker comparisons greater than the 95% quantile.

| LG Comparison | r | N | r95 | N95 | Proportion | RIL Population |
|---|---|---|---|---|---|---|
| 01,02 | 0.1003 | 2940 | NA | 0 | 0.0000 | CAC110 |
| 01,02 | 0.0871 | 1692 | NA | 0 | 0.0000 | CAC162 |
| 01,02 | 0.0822 | 1804 | NA | 0 | 0.0000 | CAC415 |
| 01,03 | 0.0948 | 2400 | NA | 0 | 0.0000 | CAC110 |
| **01,03** | **0.2244** | **1944** | **0.4251** | **25** | **0.0129** | **CAC162** |
| 01,03 | 0.0618 | 1394 | NA | 0 | 0.0000 | CAC415 |
| **01,04** | **0.0991** | **2820** | **0.4027** | **69** | **0.0245** | **CAC110** |
| 01,04 | 0.1335 | 1296 | NA | 0 | 0.0000 | CAC162 |
| **01,04** | **0.1431** | **2214** | **0.4041** | **19** | **0.0086** | **CAC415** |
| 01,05 | 0.1085 | 2520 | NA | 0 | 0.0000 | CAC110 |
| 01,05 | 0.1097 | 1296 | NA | 0 | 0.0000 | CAC162 |
| 01,05 | 0.1040 | 2050 | NA | 0 | 0.0000 | CAC415 |
| 01,06 | 0.0697 | 3420 | NA | 0 | 0.0000 | CAC110 |
| 01,06 | 0.0808 | 1728 | NA | 0 | 0.0000 | CAC162 |
| 01,06 | 0.0677 | 2091 | NA | 0 | 0.0000 | CAC415 |
| 01,07 | 0.1052 | 2760 | NA | 0 | 0.0000 | CAC110 |
| 01,07 | 0.0818 | 1296 | NA | 0 | 0.0000 | CAC162 |
| 01,07 | 0.0673 | 1886 | NA | 0 | 0.0000 | CAC415 |
| 01,08 | 0.0745 | 5280 | NA | 0 | 0.0000 | CAC110 |
| 01,08 | 0.1135 | 1944 | NA | 0 | 0.0000 | CAC162 |
| 01,08 | 0.1408 | 2296 | NA | 0 | 0.0000 | CAC415 |
| 01,09 | 0.0883 | 2460 | NA | 0 | 0.0000 | CAC110 |
| 01,09 | 0.1120 | 1620 | NA | 0 | 0.0000 | CAC162 |
| 01,09 | 0.0620 | 2009 | NA | 0 | 0.0000 | CAC415 |
| 01,10 | 0.0722 | 3300 | NA | 0 | 0.0000 | CAC110 |
| **01,10** | **0.1510** | **900** | **0.4638** | **43** | **0.0478** | **CAC162** |
| 01,10 | 0.0770 | 1353 | NA | 0 | 0.0000 | CAC415 |
| **01,11** | **0.1260** | **4020** | **0.3961** | **64** | **0.0159** | **CAC110** |
| 01,11 | 0.0539 | 1260 | NA | 0 | 0.0000 | CAC162 |
| 01,11 | 0.0785 | 2050 | NA | 0 | 0.0000 | CAC415 |
| 01,12 | 0.0832 | 4080 | NA | 0 | 0.0000 | CAC110 |

| | | | | | | |
|---|---|---|---|---|---|---|
| 01,12 | 0.1656 | 1692 | NA | 0 | 0.0000 | CAC162 |
| 01,12 | 0.0490 | 1886 | NA | 0 | 0.0000 | CAC415 |
| **01,13** | **0.1026** | **3000** | **0.3831** | **24** | **0.0080** | **CAC110** |
| 01,13 | 0.0816 | 1368 | NA | 0 | 0.0000 | CAC162 |
| **01,13** | **0.1492** | **2378** | **0.4022** | **9** | **0.0038** | **CAC415** |
| 01,14 | 0.0864 | 5040 | NA | 0 | 0.0000 | CAC110 |
| 01,14 | 0.1142 | 2484 | NA | 0 | 0.0000 | CAC162 |
| **01,14** | **0.1319** | **3198** | **0.3934** | **5** | **0.0016** | **CAC415** |
| 02,03 | 0.0867 | 1960 | NA | 0 | 0.0000 | CAC110 |
| 02,03 | 0.0784 | 2538 | NA | 0 | 0.0000 | CAC162 |
| 02,03 | 0.0911 | 1496 | NA | 0 | 0.0000 | CAC415 |
| 02,04 | 0.0808 | 2303 | NA | 0 | 0.0000 | CAC110 |
| 02,04 | 0.0946 | 1692 | NA | 0 | 0.0000 | CAC162 |
| 02,04 | 0.0634 | 2376 | NA | 0 | 0.0000 | CAC415 |
| 02,05 | 0.0616 | 2058 | NA | 0 | 0.0000 | CAC110 |
| 02,05 | 0.1019 | 1692 | NA | 0 | 0.0000 | CAC162 |
| 02,05 | 0.0842 | 2200 | NA | 0 | 0.0000 | CAC415 |
| 02,06 | 0.0682 | 2793 | NA | 0 | 0.0000 | CAC110 |
| 02,06 | 0.0975 | 2256 | NA | 0 | 0.0000 | CAC162 |
| 02,06 | 0.0884 | 2244 | NA | 0 | 0.0000 | CAC415 |
| 02,07 | 0.0848 | 2254 | NA | 0 | 0.0000 | CAC110 |
| 02,07 | 0.0742 | 1692 | NA | 0 | 0.0000 | CAC162 |
| 02,07 | 0.1123 | 2024 | NA | 0 | 0.0000 | CAC415 |
| 02,08 | 0.0958 | 4312 | NA | 0 | 0.0000 | CAC110 |
| 02,08 | 0.1219 | 2538 | NA | 0 | 0.0000 | CAC162 |
| 02,08 | 0.0753 | 2464 | NA | 0 | 0.0000 | CAC415 |
| 02,09 | 0.0952 | 2009 | NA | 0 | 0.0000 | CAC110 |
| 02,09 | 0.0915 | 2115 | NA | 0 | 0.0000 | CAC162 |
| 02,09 | 0.0974 | 2156 | NA | 0 | 0.0000 | CAC415 |
| 02,10 | 0.0520 | 2695 | NA | 0 | 0.0000 | CAC110 |
| 02,10 | 0.0531 | 1175 | NA | 0 | 0.0000 | CAC162 |
| 02,10 | 0.1337 | 1452 | NA | 0 | 0.0000 | CAC415 |
| 02,11 | 0.0792 | 3283 | NA | 0 | 0.0000 | CAC110 |
| 02,11 | 0.1718 | 1645 | NA | 0 | 0.0000 | CAC162 |
| 02,11 | 0.0674 | 2200 | NA | 0 | 0.0000 | CAC415 |
| 02,12 | 0.0401 | 3332 | NA | 0 | 0.0000 | CAC110 |
| 02,12 | 0.0882 | 2209 | NA | 0 | 0.0000 | CAC162 |
| 02,12 | 0.0780 | 2024 | NA | 0 | 0.0000 | CAC415 |
| **02,13** | **0.1408** | **2450** | **0.4083** | **90** | **0.0367** | **CAC110** |
| 02,13 | 0.1008 | 1786 | NA | 0 | 0.0000 | CAC162 |

| | | | | | | |
|---|---|---|---|---|---|---|
| 02,13 | 0.0684 | 2552 | NA | 0 | 0.0000 | CAC415 |
| 02,14 | 0.0945 | 4116 | NA | 0 | 0.0000 | CAC110 |
| 02,14 | 0.1041 | 3243 | NA | 0 | 0.0000 | CAC162 |
| 02,14 | 0.0707 | 3432 | NA | 0 | 0.0000 | CAC415 |
| 03,04 | 0.0496 | 1880 | NA | 0 | 0.0000 | CAC110 |
| 03,04 | 0.0856 | 1944 | NA | 0 | 0.0000 | CAC162 |
| 03,04 | 0.1287 | 1836 | NA | 0 | 0.0000 | CAC415 |
| 03,05 | 0.1039 | 1680 | NA | 0 | 0.0000 | CAC110 |
| 03,05 | 0.1435 | 1944 | NA | 0 | 0.0000 | CAC162 |
| 03,05 | 0.0768 | 1700 | NA | 0 | 0.0000 | CAC415 |
| 03,06 | 0.0940 | 2280 | NA | 0 | 0.0000 | CAC110 |
| 03,06 | 0.0809 | 2592 | NA | 0 | 0.0000 | CAC162 |
| **03,06** | **0.1284** | **1734** | **0.4024** | **5** | **0.0029** | **CAC415** |
| 03,07 | 0.0752 | 1840 | NA | 0 | 0.0000 | CAC110 |
| 03,07 | 0.0582 | 1944 | NA | 0 | 0.0000 | CAC162 |
| 03,07 | 0.1069 | 1564 | NA | 0 | 0.0000 | CAC415 |
| 03,08 | 0.0851 | 3520 | NA | 0 | 0.0000 | CAC110 |
| 03,08 | 0.1265 | 2916 | NA | 0 | 0.0000 | CAC162 |
| 03,08 | 0.0799 | 1904 | NA | 0 | 0.0000 | CAC415 |
| 03,09 | 0.0865 | 1640 | NA | 0 | 0.0000 | CAC110 |
| 03,09 | 0.1260 | 2430 | NA | 0 | 0.0000 | CAC162 |
| 03,09 | 0.1058 | 1666 | NA | 0 | 0.0000 | CAC415 |
| 03,10 | 0.0897 | 2200 | NA | 0 | 0.0000 | CAC110 |
| 03,10 | 0.1326 | 1350 | NA | 0 | 0.0000 | CAC162 |
| 03,10 | 0.0958 | 1122 | NA | 0 | 0.0000 | CAC415 |
| **03,11** | **0.1867** | **2680** | **0.3860** | **69** | **0.0257** | **CAC110** |
| 03,11 | 0.1129 | 1890 | NA | 0 | 0.0000 | CAC162 |
| 03,11 | 0.0647 | 1700 | NA | 0 | 0.0000 | CAC415 |
| 03,12 | 0.0964 | 2720 | NA | 0 | 0.0000 | CAC110 |
| 03,12 | 0.0840 | 2538 | NA | 0 | 0.0000 | CAC162 |
| 03,12 | 0.0761 | 1564 | NA | 0 | 0.0000 | CAC415 |
| **03,13** | **0.1453** | **2000** | **0.4824** | **64** | **0.0320** | **CAC110** |
| 03,13 | 0.0684 | 2052 | NA | 0 | 0.0000 | CAC162 |
| 03,13 | 0.1268 | 1972 | NA | 0 | 0.0000 | CAC415 |
| 03,14 | 0.0543 | 3360 | NA | 0 | 0.0000 | CAC110 |
| 03,14 | 0.1271 | 3726 | NA | 0 | 0.0000 | CAC162 |
| 03,14 | 0.0718 | 2652 | NA | 0 | 0.0000 | CAC415 |
| 04,05 | 0.0623 | 1974 | NA | 0 | 0.0000 | CAC110 |
| 04,05 | 0.0706 | 1296 | NA | 0 | 0.0000 | CAC162 |
| **04,05** | **0.1399** | **2700** | **0.3988** | **3** | **0.0011** | **CAC415** |

| | | | | | | |
|---|---|---|---|---|---|---|
| 04,06 | 0.0690 | 2679 | NA | 0 | 0.0000 | CAC110 |
| 04,06 | 0.0548 | 1728 | NA | 0 | 0.0000 | CAC162 |
| 04,06 | 0.0895 | 2754 | NA | 0 | 0.0000 | CAC415 |
| 04,07 | 0.0548 | 2162 | NA | 0 | 0.0000 | CAC110 |
| 04,07 | 0.1111 | 1296 | NA | 0 | 0.0000 | CAC162 |
| 04,07 | 0.0962 | 2484 | NA | 0 | 0.0000 | CAC415 |
| 04,08 | 0.0768 | 4136 | NA | 0 | 0.0000 | CAC110 |
| 04,08 | 0.1041 | 1944 | NA | 0 | 0.0000 | CAC162 |
| **04,08** | **0.1439** | **3024** | **0.3979** | **1** | **0.0003** | **CAC415** |
| 04,09 | 0.0735 | 1927 | NA | 0 | 0.0000 | CAC110 |
| 04,09 | 0.1238 | 1620 | NA | 0 | 0.0000 | CAC162 |
| 04,09 | 0.1040 | 2646 | NA | 0 | 0.0000 | CAC415 |
| 04,10 | 0.0626 | 2585 | NA | 0 | 0.0000 | CAC110 |
| 04,10 | 0.1461 | 900 | NA | 0 | 0.0000 | CAC162 |
| 04,10 | 0.0672 | 1782 | NA | 0 | 0.0000 | CAC415 |
| 04,11 | 0.0755 | 3149 | NA | 0 | 0.0000 | CAC110 |
| 04,11 | 0.0728 | 1260 | NA | 0 | 0.0000 | CAC162 |
| 04,11 | 0.1005 | 2700 | NA | 0 | 0.0000 | CAC415 |
| 04,12 | 0.0470 | 3196 | NA | 0 | 0.0000 | CAC110 |
| 04,12 | 0.1031 | 1692 | NA | 0 | 0.0000 | CAC162 |
| 04,12 | 0.1035 | 2484 | NA | 0 | 0.0000 | CAC415 |
| **04,13** | **0.1303** | **2350** | **0.3716** | **2** | **0.0009** | **CAC110** |
| 04,13 | 0.1510 | 1368 | NA | 0 | 0.0000 | CAC162 |
| 04,13 | 0.1289 | 3132 | NA | 0 | 0.0000 | CAC415 |
| 04,14 | 0.0531 | 3948 | NA | 0 | 0.0000 | CAC110 |
| **04,14** | **0.1708** | **2484** | **0.4401** | **17** | **0.0068** | **CAC162** |
| **04,14** | **0.2961** | **4212** | **0.4512** | **832** | **0.1975** | **CAC415** |
| 05,06 | 0.0675 | 2394 | NA | 0 | 0.0000 | CAC110 |
| 05,06 | 0.1297 | 1728 | NA | 0 | 0.0000 | CAC162 |
| **05,06** | **0.0983** | **2550** | **0.4037** | **15** | **0.0059** | **CAC415** |
| 05,07 | 0.0628 | 1932 | NA | 0 | 0.0000 | CAC110 |
| 05,07 | 0.1130 | 1296 | NA | 0 | 0.0000 | CAC162 |
| **05,07** | **0.1240** | **2300** | **0.4012** | **4** | **0.0017** | **CAC415** |
| 05,08 | 0.0508 | 3696 | NA | 0 | 0.0000 | CAC110 |
| 05,08 | 0.0812 | 1944 | NA | 0 | 0.0000 | CAC162 |
| **05,08** | **0.1105** | **2800** | **0.3940** | **1** | **0.0004** | **CAC415** |
| 05,09 | 0.0724 | 1722 | NA | 0 | 0.0000 | CAC110 |
| 05,09 | 0.1111 | 1620 | NA | 0 | 0.0000 | CAC162 |
| 05,09 | 0.1016 | 2450 | NA | 0 | 0.0000 | CAC415 |
| 05,10 | 0.0599 | 2310 | NA | 0 | 0.0000 | CAC110 |

| | | | | | | |
|---|---|---|---|---|---|---|
| 05,10 | 0.0734 | 900 | NA | 0 | 0.0000 | CAC162 |
| 05,10 | 0.1091 | 1650 | NA | 0 | 0.0000 | CAC415 |
| **05,11** | **0.2175** | **2814** | **0.4226** | **474** | **0.1684** | **CAC110** |
| 05,11 | 0.1003 | 1260 | NA | 0 | 0.0000 | CAC162 |
| 05,11 | 0.1079 | 2500 | NA | 0 | 0.0000 | CAC415 |
| 05,12 | 0.0704 | 2856 | NA | 0 | 0.0000 | CAC110 |
| 05,12 | 0.1033 | 1692 | NA | 0 | 0.0000 | CAC162 |
| **05,12** | **0.1176** | **2300** | **0.3957** | **4** | **0.0017** | **CAC415** |
| **05,13** | **0.1133** | **2100** | **0.3957** | **13** | **0.0062** | **CAC110** |
| 05,13 | 0.0733 | 1368 | NA | 0 | 0.0000 | CAC162 |
| 05,13 | 0.1121 | 2900 | NA | 0 | 0.0000 | CAC415 |
| 05,14 | 0.0616 | 3528 | NA | 0 | 0.0000 | CAC110 |
| 05,14 | 0.0864 | 2484 | NA | 0 | 0.0000 | CAC162 |
| **05,14** | **0.1589** | **3900** | **0.4167** | **94** | **0.0241** | **CAC415** |
| 06,07 | 0.0763 | 2622 | NA | 0 | 0.0000 | CAC110 |
| 06,07 | 0.0821 | 1728 | NA | 0 | 0.0000 | CAC162 |
| 06,07 | 0.1154 | 2346 | NA | 0 | 0.0000 | CAC415 |
| 06,08 | 0.0677 | 5016 | NA | 0 | 0.0000 | CAC110 |
| 06,08 | 0.1192 | 2592 | NA | 0 | 0.0000 | CAC162 |
| **06,08** | **0.1273** | **2856** | **0.4106** | **23** | **0.0081** | **CAC415** |
| 06,09 | 0.0496 | 2337 | NA | 0 | 0.0000 | CAC110 |
| 06,09 | 0.1248 | 2160 | NA | 0 | 0.0000 | CAC162 |
| 06,09 | 0.0741 | 2499 | NA | 0 | 0.0000 | CAC415 |
| 06,10 | 0.0717 | 3135 | NA | 0 | 0.0000 | CAC110 |
| 06,10 | 0.0845 | 1200 | NA | 0 | 0.0000 | CAC162 |
| **06,10** | **0.1326** | **1683** | **0.4304** | **10** | **0.0059** | **CAC415** |
| 06,11 | 0.1348 | 3819 | NA | 0 | 0.0000 | CAC110 |
| 06,11 | 0.0890 | 1680 | NA | 0 | 0.0000 | CAC162 |
| 06,11 | 0.0867 | 2550 | NA | 0 | 0.0000 | CAC415 |
| 06,12 | 0.0634 | 3876 | NA | 0 | 0.0000 | CAC110 |
| **06,12** | **0.2184** | **2256** | **0.4341** | **83** | **0.0368** | **CAC162** |
| 06,12 | 0.1248 | 2346 | NA | 0 | 0.0000 | CAC415 |
| 06,13 | 0.0843 | 2850 | NA | 0 | 0.0000 | CAC110 |
| 06,13 | 0.0812 | 1824 | NA | 0 | 0.0000 | CAC162 |
| 06,13 | 0.0995 | 2958 | NA | 0 | 0.0000 | CAC415 |
| 06,14 | 0.0485 | 4788 | NA | 0 | 0.0000 | CAC110 |
| 06,14 | 0.0892 | 3312 | NA | 0 | 0.0000 | CAC162 |
| **06,14** | **0.1300** | **3978** | **0.4029** | **1** | **0.0003** | **CAC415** |
| 07,08 | 0.0530 | 4048 | NA | 0 | 0.0000 | CAC110 |
| 07,08 | 0.1047 | 1944 | NA | 0 | 0.0000 | CAC162 |

| | | | | | | |
|---|---|---|---|---|---|---|
| **07,08** | **0.1350** | **2576** | **0.4180** | **32** | **0.0124** | **CAC415** |
| 07,09 | 0.0974 | 1886 | NA | 0 | 0.0000 | CAC110 |
| 07,09 | 0.1520 | 1620 | NA | 0 | 0.0000 | CAC162 |
| 07,09 | 0.0977 | 2254 | NA | 0 | 0.0000 | CAC415 |
| 07,10 | 0.0691 | 2530 | NA | 0 | 0.0000 | CAC110 |
| 07,10 | 0.0948 | 900 | NA | 0 | 0.0000 | CAC162 |
| 07,10 | 0.0828 | 1518 | NA | 0 | 0.0000 | CAC415 |
| 07,11 | 0.0750 | 3082 | NA | 0 | 0.0000 | CAC110 |
| 07,11 | 0.0862 | 1260 | NA | 0 | 0.0000 | CAC162 |
| 07,11 | 0.1393 | 2300 | NA | 0 | 0.0000 | CAC415 |
| 07,12 | 0.0747 | 3128 | NA | 0 | 0.0000 | CAC110 |
| 07,12 | 0.0811 | 1692 | NA | 0 | 0.0000 | CAC162 |
| 07,12 | 0.0939 | 2116 | NA | 0 | 0.0000 | CAC415 |
| 07,13 | 0.0689 | 2300 | NA | 0 | 0.0000 | CAC110 |
| 07,13 | 0.1137 | 1368 | NA | 0 | 0.0000 | CAC162 |
| 07,13 | 0.0472 | 2668 | NA | 0 | 0.0000 | CAC415 |
| 07,14 | 0.0760 | 3864 | NA | 0 | 0.0000 | CAC110 |
| 07,14 | 0.1120 | 2484 | NA | 0 | 0.0000 | CAC162 |
| 07,14 | 0.0740 | 3588 | NA | 0 | 0.0000 | CAC415 |
| 08,09 | 0.0781 | 3608 | NA | 0 | 0.0000 | CAC110 |
| 08,09 | 0.1185 | 2430 | NA | 0 | 0.0000 | CAC162 |
| 08,09 | 0.1120 | 2744 | NA | 0 | 0.0000 | CAC415 |
| 08,10 | 0.0597 | 4840 | NA | 0 | 0.0000 | CAC110 |
| 08,10 | 0.1139 | 1350 | NA | 0 | 0.0000 | CAC162 |
| 08,10 | 0.0935 | 1848 | NA | 0 | 0.0000 | CAC415 |
| 08,11 | 0.0939 | 5896 | NA | 0 | 0.0000 | CAC110 |
| 08,11 | 0.0979 | 1890 | NA | 0 | 0.0000 | CAC162 |
| 08,11 | 0.0908 | 2800 | NA | 0 | 0.0000 | CAC415 |
| 08,12 | 0.0849 | 5984 | NA | 0 | 0.0000 | CAC110 |
| 08,12 | 0.0863 | 2538 | NA | 0 | 0.0000 | CAC162 |
| 08,12 | 0.0740 | 2576 | NA | 0 | 0.0000 | CAC415 |
| 08,13 | 0.1100 | 4400 | NA | 0 | 0.0000 | CAC110 |
| 08,13 | 0.0866 | 2052 | NA | 0 | 0.0000 | CAC162 |
| 08,13 | 0.1011 | 3248 | NA | 0 | 0.0000 | CAC415 |
| 08,14 | 0.0686 | 7392 | NA | 0 | 0.0000 | CAC110 |
| 08,14 | 0.1274 | 3726 | NA | 0 | 0.0000 | CAC162 |
| 08,14 | 0.0995 | 4368 | NA | 0 | 0.0000 | CAC415 |
| 09,10 | 0.0904 | 2255 | NA | 0 | 0.0000 | CAC110 |
| 09,10 | 0.1074 | 1125 | NA | 0 | 0.0000 | CAC162 |
| 09,10 | 0.0758 | 1617 | NA | 0 | 0.0000 | CAC415 |

| | | | | | | |
|---|---|---|---|---|---|---|
| **09,11** | **0.1469** | **2747** | **0.3752** | **2** | **0.0007** | **CAC110** |
| 09,11 | 0.0711 | 1575 | NA | 0 | 0.0000 | CAC162 |
| 09,11 | 0.0896 | 2450 | NA | 0 | 0.0000 | CAC415 |
| 09,12 | 0.0639 | 2788 | NA | 0 | 0.0000 | CAC110 |
| 09,12 | 0.1226 | 2115 | NA | 0 | 0.0000 | CAC162 |
| 09,12 | 0.1003 | 2254 | NA | 0 | 0.0000 | CAC415 |
| 09,13 | 0.0699 | 2050 | NA | 0 | 0.0000 | CAC110 |
| 09,13 | 0.0717 | 1710 | NA | 0 | 0.0000 | CAC162 |
| **09,13** | **0.1417** | **2842** | **0.4009** | **1** | **0.0004** | **CAC415** |
| 09,14 | 0.0612 | 3444 | NA | 0 | 0.0000 | CAC110 |
| 09,14 | 0.1531 | 3105 | NA | 0 | 0.0000 | CAC162 |
| 09,14 | 0.0679 | 3822 | NA | 0 | 0.0000 | CAC415 |
| 10,11 | 0.0930 | 3685 | NA | 0 | 0.0000 | CAC110 |
| 10,11 | 0.1011 | 875 | NA | 0 | 0.0000 | CAC162 |
| 10,11 | 0.0721 | 1650 | NA | 0 | 0.0000 | CAC415 |
| 10,12 | 0.0570 | 3740 | NA | 0 | 0.0000 | CAC110 |
| 10,12 | 0.0980 | 1175 | NA | 0 | 0.0000 | CAC162 |
| 10,12 | 0.1084 | 1518 | NA | 0 | 0.0000 | CAC415 |
| 10,13 | 0.0757 | 2750 | NA | 0 | 0.0000 | CAC110 |
| 10,13 | 0.1372 | 950 | NA | 0 | 0.0000 | CAC162 |
| 10,13 | 0.0864 | 1914 | NA | 0 | 0.0000 | CAC415 |
| 10,14 | 0.0664 | 4620 | NA | 0 | 0.0000 | CAC110 |
| 10,14 | 0.1520 | 1725 | NA | 0 | 0.0000 | CAC162 |
| 10,14 | 0.1194 | 2574 | NA | 0 | 0.0000 | CAC415 |
| **11,12** | **0.1682** | **4556** | **0.3902** | **135** | **0.0296** | **CAC110** |
| 11,12 | 0.0680 | 1645 | NA | 0 | 0.0000 | CAC162 |
| 11,12 | 0.0685 | 2300 | NA | 0 | 0.0000 | CAC415 |
| 11,13 | 0.0919 | 3350 | NA | 0 | 0.0000 | CAC110 |
| 11,13 | 0.0740 | 1330 | NA | 0 | 0.0000 | CAC162 |
| 11,13 | 0.0794 | 2900 | NA | 0 | 0.0000 | CAC415 |
| 11,14 | 0.0930 | 5628 | NA | 0 | 0.0000 | CAC110 |
| 11,14 | 0.1356 | 2415 | NA | 0 | 0.0000 | CAC162 |
| **11,14** | **0.1458** | **3900** | **0.3904** | **1** | **0.0003** | **CAC415** |
| 12,13 | 0.0909 | 3400 | NA | 0 | 0.0000 | CAC110 |
| 12,13 | 0.0714 | 1786 | NA | 0 | 0.0000 | CAC162 |
| 12,13 | 0.0941 | 2668 | NA | 0 | 0.0000 | CAC415 |
| 12,14 | 0.1328 | 5712 | NA | 0 | 0.0000 | CAC110 |
| 12,14 | 0.1018 | 3243 | NA | 0 | 0.0000 | CAC162 |
| 12,14 | 0.1522 | 3588 | NA | 0 | 0.0000 | CAC415 |
| **13,14** | **0.1047** | **4200** | **0.3966** | **10** | **0.0024** | **CAC110** |

| | | | | | | |
|---|---|---|---|---|---|---|
| 13,14 | 0.1072 | 2622 | NA | 0 | 0.0000 | CAC162 |
| **13,14** | **0.1665** | **4524** | **0.4149** | **50** | **0.0111** | **CAC415** |

**Table S5. Positions of two-locus epistasis.** Physical (Mb) positions of pairs of loci on each linkage group (LG) in the three RIL populations that show LD (r) in the 95th percentile. Counts of each genotypic class in the population and Chi-Square statistics from likelihood ratio tests, with p<0.0001 are shown in bold. Loci marked with an asterisk have peak r and $\chi^2$ values at adjacent markers. Single locus proportions of *M. guttatus* alleles (G Prop.) were used to determine if the pattern of TRD at each epistatic locus could be explained by a two-locus hybrid incompatibility (NG- or GN-), or if epistatic selection favoring the same parental genotypes better explains the pattern (GG or NN). The best candidates for two locus hybrid incompatibilities are shaded grey.

| | Locus 1 | | Locus 2 | | | | | | | | | | Locus 1 | Locus 2 | |
|---|---|---|---|---|---|---|---|---|---|---|---|---|---|---|---|
| Population | LG | Mb | LG | Mb | r | GG | GN | NG | NN | N | χ2 | p | G Prop. | G Prop. | Pattern |
| | 03 | 20.40 - 22.65 | 13 | 3.05 - 5.20 | 0.56 | 64 | 3 | 17 | 19 | 103 | 32.552 | <0.0001 | 0.65 | 0.79 | GG |
| | 05 | 3.10 - 20.75 | 11 | 2.55 - 4.90 | 0.54 | 39 | 21 | 1 | 22 | 83 | 29.035 | <0.0001 | 0.72 | 0.48 | NG- |
| | 01 | 11.15 - 12.85 | 04 | 1.20 - 19.65 | 0.46 | 73 | 13 | 12 | 19 | 117 | 22.858 | <0.0001 | 0.74 | 0.73 | GG |
| | 02 | 2.85 - 18.95 | 13 | 21.75 - 23.65 | 0.45 | 54 | 1 | 26 | 14 | 95 | 21.079 | <0.0001 | 0.58 | 0.84 | GG |
| | 09 | 0.80 - 0.90 | 11 | 26.30 - 26.60 | 0.38 | 29 | 37 | 6 | 54 | 126 | 19.359 | <0.0001 | 0.52 | 0.28 | NG- |
| | 05 | 2.40 - 20.90 | 11 | 5.05 - 10.20* | 0.50 | 43 | 14 | 5 | 17 | 79 | 18.697 | <0.0001 | 0.72 | 0.61 | GG |
| | 03 | 0.15 - 18.95 | 11 | 24.35 - 25.30 | 0.40 | 29 | 24 | 12 | 56 | 121 | 18.567 | <0.0001 | 0.44 | 0.34 | NN |
| CAC110 | 11 | 8.80 - 24.40 | 12 | 1.00 - 19.2 | 0.44 | 18 | 12 | 12 | 59 | 101 | 17.988 | <0.0001 | 0.30 | 0.30 | NN |
| | 01 | 3.30 - 5.50 | 11 | 4.00 - 5.75 | 0.47 | 29 | 9 | 10 | 24 | 72 | 16.515 | <0.0001 | 0.53 | 0.54 | GG |
| | 05 | 2.25 - 3.50 | 13 | 24.00 - 25.00 | 0.44 | 54 | 13 | 7 | 14 | 88 | 15.846 | <0.0001 | 0.76 | 0.69 | GG |
| | 13 | 25.00 - 25.70 | 14 | 1.25 - 2.15 | 0.41 | 41 | 7 | 15 | 17 | 80 | 13.622 | 0.0002 | 0.60 | 0.70 | GG |
| | 03 | 18.40 - 21.55 | 11 | 3.35 - 6.25 | 0.42 | 34 | 7 | 12 | 17 | 70 | 13.194 | 0.0003 | 0.59 | 0.66 | GG |
| | 01 | 7.80 - 9.60 | 13 | 21.75 - 23.65 | 0.42 | 40 | 2 | 25 | 12 | 79 | 11.101 | 0.0009 | 0.53 | 0.82 | GG |
| | 04 | 3.80 - 3.90 | 13 | 21.40 - 21.55 | 0.37 | 60 | 1 | 16 | 5 | 82 | 9.671 | 0.0019 | 0.74 | 0.93 | GG |
| | 01 | 11.10 - 11.15 | 11 | 4.10 - 4.55 | 0.39 | 26 | 13 | 5 | 15 | 59 | 9.497 | 0.0021 | 0.66 | 0.53 | GG |
| | 04 | 19.90 - 22.80 | 14 | 2.35 - 18.15 | 0.50 | 29 | 10 | 14 | 44 | 97 | 24.708 | <0.0001 | 0.40 | 0.44 | NN |
| CAC162 | 01 | 1.40 - 5.55 | 10 | 19.65 - 19.95 | 0.54 | 27 | 9 | 8 | 31 | 75 | 23.571 | <0.0001 | 0.48 | 0.47 | NN |
| | 06 | 6.30 - 15.10 | 12 | 6.20 - 7.75 | 0.47 | 9 | 50 | 23 | 15 | 97 | 21.637 | <0.0001 | 0.61 | 0.33 | - |
| | 01 | 2.35 - 5.45 | 03 | 0.00 - 20.65 | 0.45 | 28 | 21 | 7 | 42 | 98 | 20.628 | <0.0001 | 0.50 | 0.36 | NG- |
| | 04 | 1.90 - 22.30 | 14 | 2.15 - 25.60 | 0.61 | 34 | 9 | 7 | 31 | 81 | 31.85 | <0.0001 | 0.53 | 0.51 | GG |
| | 06 | 17.00 - 19.25 | 10 | 19.50 - 20.25 | 0.43 | 38 | 0 | 22 | 13 | 73 | 22.217 | <0.0001 | 0.52 | 0.82 | GG |
| | 05 | 3.50 - 20.40 | 14 | 16.50 - 23.00 | 0.49 | 26 | 5 | 17 | 32 | 80 | 19.799 | <0.0001 | 0.39 | 0.54 | GN- |
| | 07 | 13.20 - 21.80 | 08 | 22.30 - 27.70 | 0.48 | 23 | 7 | 14 | 38 | 82 | 19.719 | <0.0001 | 0.37 | 0.45 | NN |
| | 13 | 22.05 - 25.55 | 14 | 0.00 - 3.00 | 0.47 | 41 | 6 | 15 | 20 | 82 | 18.738 | <0.0001 | 0.57 | 0.68 | GG |
| | 06 | 9.65 - 16.00 | 08 | 23.60 - 25.55 | 0.44 | 22 | 10 | 12 | 37 | 81 | 15.892 | <0.0001 | 0.40 | 0.42 | NN |
| | 01 | 5.85 - 7.40 | 04 | 4.10 - 16.55 | 0.43 | 32 | 6 | 18 | 24 | 80 | 15.338 | <0.0001 | 0.48 | 0.63 | GN- |
| | 01 | 7.70 - 11.70 | 13 | 24.20 - 25.15 | 0.42 | 24 | 4 | 22 | 30 | 80 | 15.279 | <0.0001 | 0.35 | 0.58 | GN- |
| | 05 | 20.85 - 21.75 | 06 | 7.10 - 16.05 | 0.43 | 20 | 11 | 11 | 40 | 82 | 15.242 | <0.0001 | 0.38 | 0.38 | NN |
| | 05 | 20.05 - 20.40 | 07 | 20.05 - 21.80 | 0.42 | 17 | 7 | 15 | 42 | 81 | 14.019 | 0.0002 | 0.30 | 0.40 | NN |
| CAC415 | 06 | 6.30 - 6.45 | 14 | 8.85 - 9.80 | 0.49 | 24 | 7 | 18 | 32 | 81 | 13.719 | 0.0002 | 0.38 | 0.52 | GN- |
| | 05 | 8.50 - 17.80 | 12 | 0.30 - 0.70 | 0.40 | 23 | 7 | 18 | 33 | 81 | 13.458 | 0.0002 | 0.37 | 0.51 | GN- |
| | 03 | 15.95 - 19.85 | 06 | 17.65 - 20.05 | 0.41 | 44 | 18 | 5 | 15 | 82 | 13.338 | 0.0003 | 0.76 | 0.60 | GG |
| | 04 | 19.75 - 19.90 | 08 | 2.20 - 2.50 | 0.40 | 13 | 36 | 22 | 11 | 82 | 13.207 | 0.0003 | 0.60 | 0.43 | - |
| | 05 | 21.40 - 21.75 | 08 | 22.10 - 22.15* | 0.39 | 22 | 9 | 16 | 35 | 82 | 12.436 | 0.0004 | 0.38 | 0.46 | NN |
| | 01 | 3.35 - 3.40 | 14 | 20.65 - 22.70 | 0.40 | 35 | 18 | 5 | 17 | 75 | 12.133 | 0.0005 | 0.71 | 0.53 | GG |
| | 11 | 24.70 - 25.50 | 14 | 25.10 - 25.15* | 0.39 | 32 | 15 | 10 | 24 | 81 | 12.119 | 0.0005 | 0.58 | 0.52 | GG |
| | 07 | 1.00 - 1.45 | 08 | 18.50 - 18.55* | 0.40 | 13 | 30 | 26 | 12 | 81 | 12.078 | 0.0005 | 0.53 | 0.48 | - |
| | 04 | 20.05 - 20.10 | 05 | 2.40 - 3.50 | 0.40 | 33 | 11 | 10 | 19 | 73 | 12.024 | 0.0005 | 0.60 | 0.59 | GG |
| | 05 | 0.50 - 1.20 | 06 | 1.45 - 1.50 | 0.40 | 23 | 21 | 2 | 18 | 64 | 11.726 | 0.0006 | 0.69 | 0.39 | NG- |
| | 09 | 23.70 - 23.85 | 13 | 26.20 - 27.70* | 0.40 | 14 | 25 | 30 | 11 | 80 | 11.494 | 0.0007 | 0.49 | 0.55 | - |
